# Single-cell multiomics of neuron activation reveals context-specific genetics of brain disorders

**DOI:** 10.1101/2025.02.17.638682

**Authors:** Lifan Liang, Siwei Zhang, Zicheng Wang, Hanwen Zhang, Chuxuan Li, Christina Thapa, Emily K. Oh, David Sirkin, Xiaotong Sun, Alexandra Barishman, Ada McCarroll, Alexandra C. Duhe, Sheng Qian, Xiaoyuan Zhong, Brendan Jamison, Whitney Wood, Alena Kozlova, Zhiping P. Pang, Alan R. Sanders, Xin He, Jubao Duan

**Affiliations:** Department of Human Genetics, The University of Chicago, Chicago, IL 60637, USA; Center for Psychiatric Genetics, Endeavor Health Research Institute, Evanston, IL 60201, USA; Graduate Group in Genomics and Computational Biology, Perelman School of Medicine, University of Pennsylvania, Philadelphia, PA 19104, USA; Department of Neuroscience and Cell Biology, Child Health Institute of New Jersey, Rutgers Robert Wood Johnson Medical School, New Brunswick, NJ 08901, USA; Department of Psychiatry and Behavioral Neuroscience, The University of Chicago, Chicago, IL 60637, USA

## Abstract

Most causal variants for neuropsychiatric disorders (NPD) remain unknown. A major hurdle is that disease variants may act in specific contexts, such as during neuronal activation, which is difficult to study in vivo at the population level. We profiled single-nucleus neuron-activation multiomics in human induced pluripotent stem cell (iPSC)-derived neurons from 100 donors, revealing the NPD-relevant transcriptomic/epigenomic landscape of neuronal activation. We identified abundant genetic variants associated with activity-dependent gene expression (eQTL) and chromatin accessibility (caQTL), the latter explaining larger proportions of NPD heritability. Integrating multiomic data with genome-wide association studies (GWAS) further revealed NPD risk variants and genes with effects detected only upon stimulation, such as activity-dependent cholesterol metabolism. Our work highlights the power of cell stimulation to reveal context-specific “hidden” genetic effects.

## Main Text

GWAS have identified hundreds of NPD risk loci, with over 280 for schizophrenia (SCZ) (*1–7*). However, despite extensive functional genomics study in postmortem brains (*8–11*) and in human iPSC-induced neurons from large cohorts (*12, 13*), causal variants/genes for NPD remain largely unknown. Most GWAS risk variants are in noncoding regions that lack functional interpretation and likely act in specific biological contexts (*14–17*), meaning that disease variants may only show detectable function in response to stimuli, as found for some other disorders (*14, 18*). It is thus important to understand NPD genetic risk in specific biological contexts such as neuronal activation.

Neuronal activity regulates neurodevelopment and synaptic plasticity (*19*), processes important in developing NPD. Neuronal activation leads to Ca^2+^ influx, activating early response genes (ERGs such as *Fos*), further activating late response genes (LRGs such as *Bdnf*) (*19, 20*). In mouse, neuronal activity alters open chromatin regions (OCRs), accompanied by expression changes of LRGs within 4-6 h (*21–25*). Stimuli such as membrane-depolarizing levels of potassium chloride (KCl) (*19, 21, 22, 26, 27*) activate neurons, mimicking the in vivo effects of social experiences, stress, or drugs of abuse (*19*). KCl stimulation of iPSC-induced neurons (iNs) elicits extensive activity-dependent transcriptomic and epigenomic alterations (*28, 29*). However, genetic regulation of cell type- and individual-specific responses to neuronal stimulation as well as its functional relevance to NPD remain elusive.

We profiled single-nucleus multiomics (sn-multiomics) (RNA sequencing and assay for transposase-accessible chromatin with sequencing; snRNA-seq/ATAC-seq) of neuronal activity-dependent transcriptomic and epigenomic changes in co-cultured human excitatory and inhibitory iNs from 100 donors (Fig. 1A). The multi-modality assay allowed us to identify gene regulatory networks (GRN) controlling neuronal activation, shedding light on the role of activity-dependent transcriptional factors (TFs) in NPD. We identified thousands of cell-type-specific and activity-dependent quantitative trait loci for gene expression (eQTLs) and chromatin accessibility (caQTLs), helping prioritize NPD risk variants and genes that manifested functional effects only upon neuronal stimulation.

**Fig. 1.**
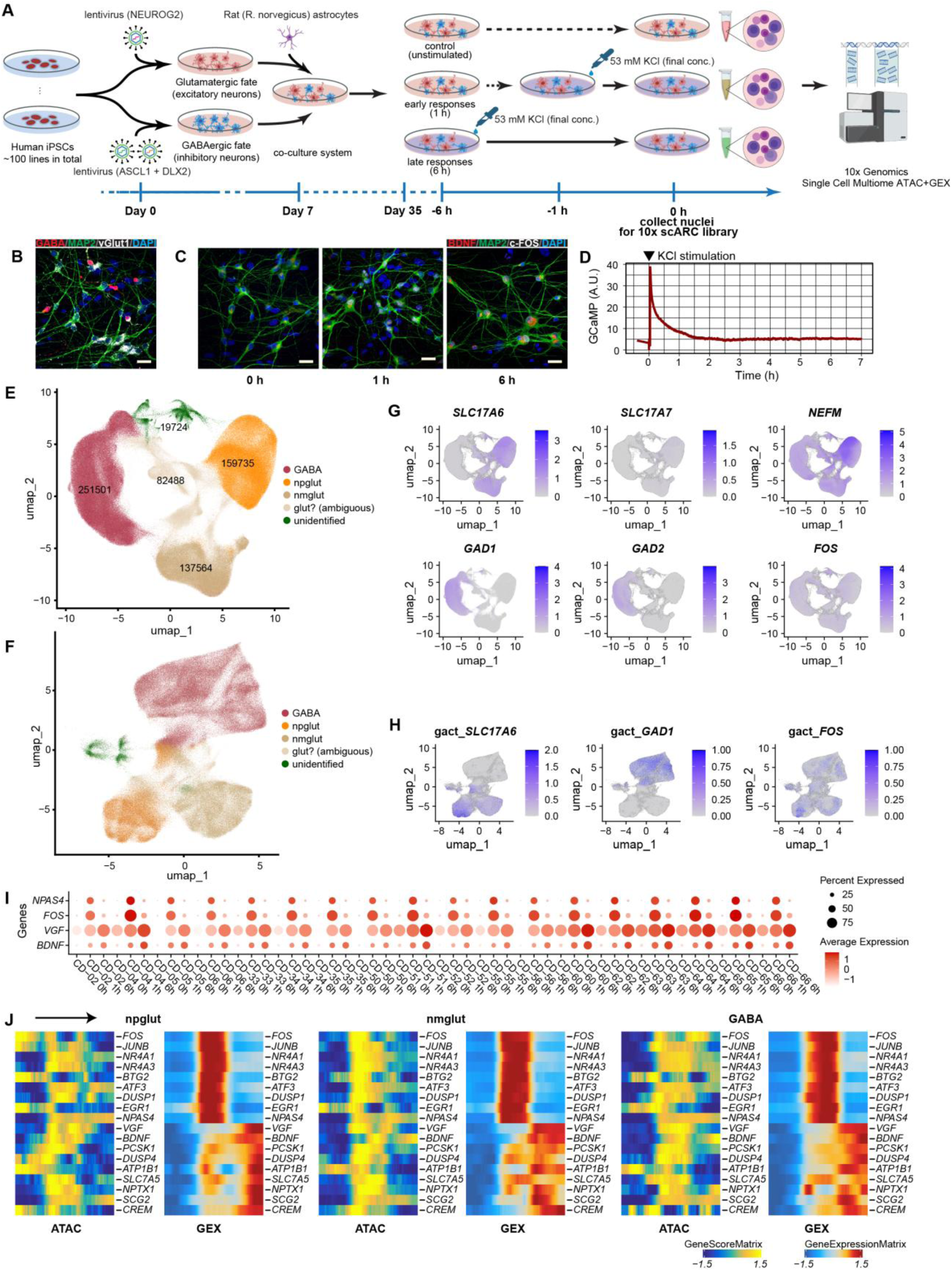
Single-nucleus multiomic assay of cell co-cultures that model neuronal activation. **(A)** Schematic of the experimental design. Co-cultures of excitatory and inhibitory neurons with rat astrocytes were stimulated by KCl to mimic neuronal activation. **(B)** IF staining of the co-cultures shows glutamatergic excitatory neurons (vGlut1+) and GABAergic inhibitory neurons (GABA+). MAP2+, neurons; DAPI, nuclei. **(C)** IF staining of early (1 h) response gene *c-FOS* and late (6 h) response gene *BDNF*. DAPI, nuclei. Scale bar=25 µm. **(D)** Ca^2+^ influx spike (fluorescence intensity of GCaMP) upon KCl stimulation. **(E)** UMAP projection and cell identities for snRNA-seq data of 100 lines. **(F)** UMAP projection for snATAC-seq data (cell identify label was transferred from snRNA-seq). **(G)** Feature plots of gene expression (snRNA-seq) for cell-type-specific genes (*SLC17A6* and *SLC17A7* for iGlut, *GAD1* and *GAD2* for iGABA, *NEFM* for two subclusters of iGlut) and *FOS*. **(H)** Feature plots of gene activity score (snATAC-seq) for cell-type-specific genes (*SLC17A6* for iGlut, *GAD1* for iGABA) and *FOS*. **(I)** Reproducible expression dynamics of early (*NPAS4, FOS*) and late (*VGF, BDNF*) response genes from 0 h, 1 h, to 6 h across cell lines (shown are 18 lines). **(J)** Pseudotime trajectories of gene expression and chromatin accessibility (GeneScore) of selected early and later response genes in each cell type.

## Results

### Sn-Multiomic profiling of human neuronal activation

To model neuronal activation, we differentiated a large cohort of human iPSC lines into glutamatergic neurons (iGlut, excitatory) (*30*) and GABAergic neurons (iGABA, inhibitory) (*31, 32*) (Fig. 1A). Day-35 iGlut and iGABA neurons, co-cultured with rat astrocytes, were stimulated by KCl (53 mM) for 1 or 6 h (*24, 25, 27, 33*) to capture changes of ERGs and LRGs, respectively. We verified the iGlut or iGABA neuronal identity, purity, and the expected expression dynamics of ERG *FOS* and LRG *BDNF* by immunostaining (Fig. 1B-C). We also confirmed effective neuronal activation by showing robust Ca^2+^ spikes immediately after KCl stimulation in GCaMP-infected neurons (Fig. 1D).

We obtained sn-multiomics data from the neural co-cultures of 100 iPSC lines at 0, 1, and 6 h after KCl stimulation (Table S1). We analyzed 1,053,422 nuclei (Tables S1-2), of which 651,012 passed snRNA/ATAC-seq quality control (QC) (Fig. S1-S3, S4A-D, Tables S2-3, Supplementary text). We defined three major neuronal subtypes: GABA (*n*=251,501), *NEFM*+ Glut (npglut; with stronger *NEFM* expression) (*n*=159,735), and *NEFM*-Glut (nmglut; with weaker *NEFM* expression) (*n*=137,564) (Fig. 1E-H, Fig. S4E-G, Table S1), showing reproducible cell type clustering patterns across sequencing libraries (Fig. S3A-E). Compared to single-cell transcriptomic profiles of human brains (*34*), our iGlut and iGABA are mostly similar to neurons of early brain developmental stages (from second trimester to 2 years old) (Fig. S5). We also found comparable proportions of neuronal subtypes between time points (Fig. S6A).

We first verified the changes of expression and chromatin accessibility of some known ERGs and LRGs. As expected, ERG expression rapidly increased from 0 to 1 h and diminished at 6 h of KCl stimulation, while LRG expression often peaked at 6 h (Fig. 1I, Fig. S6B-C). However, their chromatin accessibility did not seem to follow expression dynamics, with most ERGs still exhibiting a prolonged chromatin openness at 6 h while most LRGs showed robust chromatin openness at 1 h before their expression peaked at 6 h (Fig. S6B-C). The observed discordance between gene expression and chromatin accessibility at discrete time points was further confirmed by a continuous pseudotime trajectory analysis of single-cell gene expression and chromatin-accessibility score (Fig. S6D). ERGs and LRGs exhibited the expected expression dynamics while their gene score matrix often showed a discordant pattern (Fig. 1J). These results validated our cellular model for studying neuronal activation and highlighted the complexity of activity-dependent chromatin regulation.

We next surveyed transcriptomic and epigenomic landscapes of cell-type-specific neuronal activation and their relevance to NPD, by analyzing snRNA-seq/ATAC-seq data of the 18 lines from the sequencing batch 24 to minimize batch effects (Fig. S7A-D). About 36-65% of the expressed genes showed differential expression (DEG) (log_2_-fold change, FC, > 0.25 or < -0.25; false discovery rate, FDR < 0.05) at 1 or 6 h in any cell type (Fig. S7E, Table S4, Supplementary text). About 81-83% of ERGs and 45-74% of LRGs from an earlier study of KCl-stimulated iNs (*29*) were also DEGs at 1 h (45.5 to 60.2-fold enrichment) or 6 h (7.2 to 9.9-fold enrichment), respectively (Supplementary text) (Fig. S8A-E). Only the upregulated genes showed enrichment for synaptic genes (Fig. S9A), rare risk genes of SCZ (*35*) and autism spectrum disorder (ASD) (*36*) (Fig. S9B-E, Table S5), and for common GWAS risk of SCZ and other NPD (Fig. S10A). For chromatin landscape (Fig. 1F, Fig. S1H), we identified 150K to 300K OCR peaks across cell types/timepoints (Fig. S10B), of which 19-22% of the peaks were stimulation-specific (Fig. S10C). About 26-34% and 40-51% of the OCR peaks showed differential accessibility (DA) at 1 and 6 h, respectively (Fig. S10D-F; Table S6). For *FOS*, an ERG, we identified a stimulation-specific DA peak ∼4.2 kb downstream of its transcription start site (TSS) that may drive its early response through CEBP binding (Fig. S11A). For *BDNF*, an LRG and a SCZ risk gene (*7*), we identified DA peaks that may regulate activity-dependent *BDNF* expression (Fig. S11B, Fig. S12A-E, Table S6, Supplementary text). Interestingly, only DA peaks at 1 or 6 h of stimulation but not static (unchanged) peaks showed NPD GWAS enrichment (Fig. S12F-G).

To ascertain the in vivo relevance of our findings, we compared our results to those from mouse models (*21, 37*). We found that mouse neuronal activity-dependent genes were enriched in our KCl-stimulated DEGs (15.6-fold for ERGs, 4.4-fold for LRGs) with well-correlated log_2_FC (Supplementary text) (Fig. S8B-E). Consistent with previous in vivo studies (*21, 29, 37*), our study also showed that LRGs tend to be more cell type-specific than ERGs. Taken together, our results highlight the widespread effects of neuronal stimulation on cell-type-specific transcriptomes and chromatin accessibility as well as their relevance to NPD.

### Complex patterns of transcriptomic and epigenomic regulation of neuronal activation

To systematically characterize activity-dependent transcriptional dynamics, we performed clustering analysis to group 5,221 highly variable DEGs (FC≥2, from the 18 lines) with similar expression patterns across contexts into “modules”. We applied expression trajectory analysis on each cell type and obtained pseudotime of each cell, capturing the extent of cellular activation. We divided pseudotime into 100 bins, and clustered genes based on 300 expression measurements (100 bins, 3 cell types), which revealed 15 clusters (Fig. S13A, Table S7). While ERG clusters tended to show similar expression dynamics across cell types, LRGs had more variable patterns as previously reported (*19*): C3 was GABA-specific and C8 was Glut-specific (Fig. 2A).

**Fig. 2.**
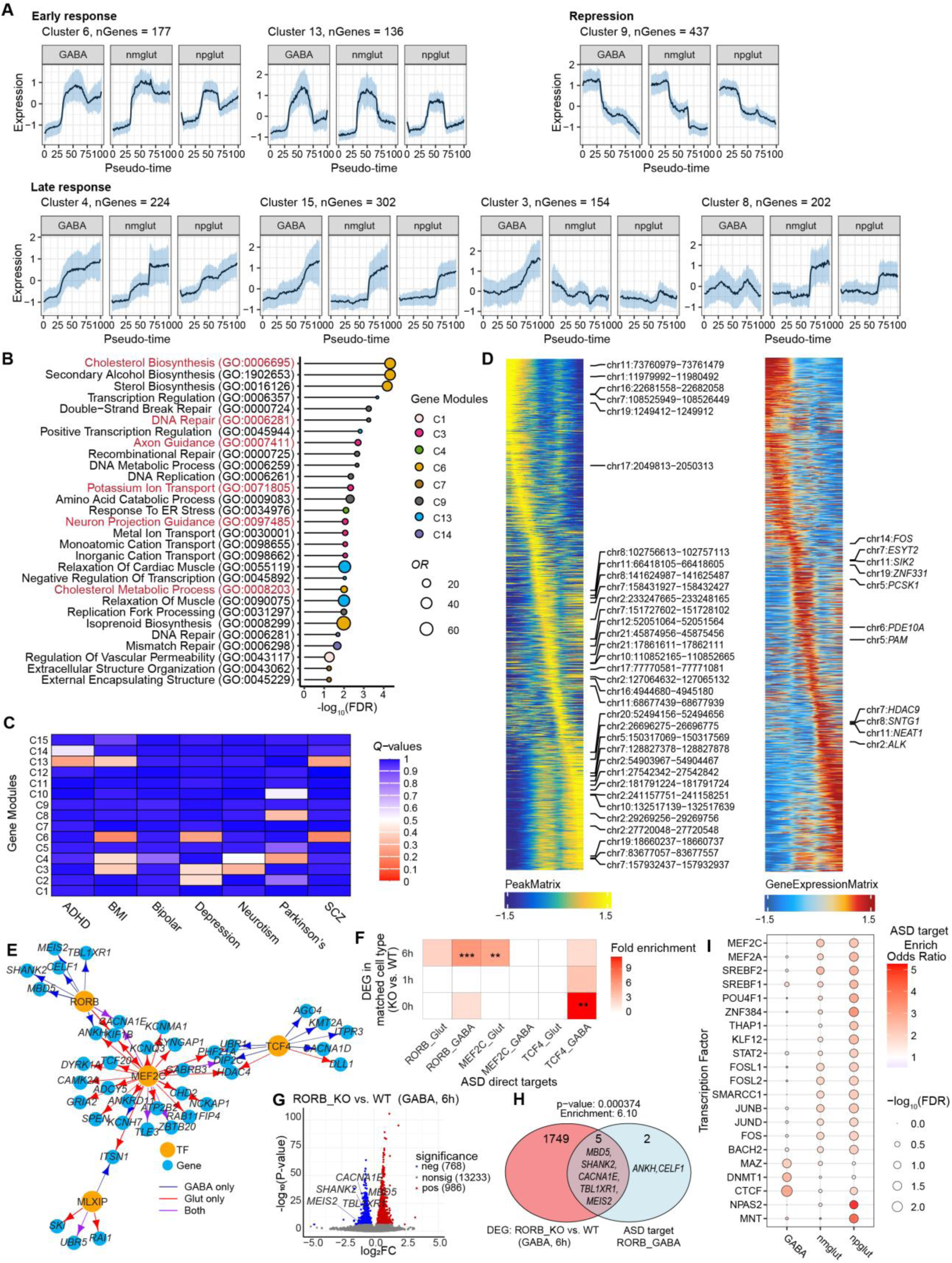
Complex transcriptomic and epigenomics regulation of neuronal activation and activity-dependent gene regulatory network (GRN). **(A)** Pseudotime trajectories of gene expression of some selected gene modules (clusters) during neuronal activation. Black line, average normalized expression of all genes in a cluster; blue shading, 50% quantile. **(B)** GO term (biological processes) enrichment for gene clusters. Terms with FDR < 0.05 are shown. **(C)** Enrichment of GWAS genes of seven complex traits in 15 gene clusters. Shown are q-values from MAGMA gene set test. (**D)** Pseudotime OCR peak activity (left) and gene expression (right) for OCR peak-gene pairs in npglut. The most variable features are labeled. (**E)** ASD-related GRN that consists of 4 selected TFs known to be ASD risk genes and their ASD risk gene targets. Arrow colors, cell type in which the associations were identified. Arrow length, inversely correlated with the magnitude of Spearman’s correlation between the motif activity of a TF and the expression of its target gene (only pairs with absolute value > 0.5). TF-target gene pairs were defined by the motif presence of the TF (*P* < 1 x 10^-5^ based on ArchR) in at least one OCRs linked to the gene (at FDR ≤ 0.1 from co-activation analysis). **(F)** to **(H)** Functional validation of ASD subnetwork (E) by CRISPR-knockout (KO) of 3 TFs (RORB, MEF2C and TCF4) in KCl-stimulated Glut/GABA co-cultures analyzed by scRNA-seq (also see Fig. S15). **(F)** Enrichment of the predicted ASD subnetwork target genes of each TF in DEGs (KO vs. WT) of their matched cell types. **, *P*<0.01, *** *P*<0.001, hypergeometric test. (**G)** Volcano plot of DEGs in RORB-KO neurons (vs. WT). Five ASD target genes are highlighted. **(H)** Venn diagram shows the overlap of the predicted ASD target genes of RORB in GABA and DEGs in GABA (RORB-KO). Enrichment *P* values were derived from hypergeometric tests. **(I)** The 21 TFs with targets enriched for ASD risk genes in at least one cell type (FDR ≤ 0.05).

We next examined the biological function of these clusters and their relevance to NPD (Fig. 2B, Table S8). Notably, the early response C6 has the most enriched gene ontology (GO) terms, with top-ranking terms related to cholesterol biosynthesis (Fig. 2B), which may contribute to NPD through affecting synaptic integrity, membrane fluidity, and neuronal signaling (*38–41*). A late response cluster, C3, was enriched for processes related to axon guidance and ion transport (Fig. 2B). These processes are relevant to NPD and body mass index (BMI) through modulating brain wiring and synaptic function important for feeding behavior and energy homeostasis (*42, 43*). The “repressive” clusters 9 and 14 were enriched for genes related to DNA repair, a process coupled to *NPAS4*-activated synaptic activity (*44*). ERG clusters C6 and C13 were enriched for GWAS signals of SCZ, major depressive disorder (MDD), BMI, and attention-deficit/hyperactivity disorder; while LRG clusters, C3 and C8, were enriched for GWAS signals for MDD, Parkinson’s disease (PD), neuroticism score, and BMI (Fig. 2C). These results support the biological relevance of these clusters to NPD genetics, and the importance of cholesterol metabolism to neuronal activation.

To characterize the activity-dependent chromatin accessibility dynamics, we first linked genes with the OCRs that likely regulate their expression. By correlating single-cell chromatin accessibility with gene expression, we obtained 8,522 OCR-gene pairs in 4,017 genes in GABA cells, 6,921 OCR-gene pairs in 3,625 genes in nmglut cells, and 7,492 OCR-gene pairs in 3,651 genes in npglut cells (Fig. 2D, Fig. S13B, Table S9). To confirm these OCR-gene pairs, we performed Micro-C in neuron co-cultures at 0, 1, and 6 h of KCl stimulation (Fig. S13C, Table S10) to identify promoter-interacting OCRs. We then carried out activity-by-contact (ABC) (*45*) analysis of the Micro-C chromatin contacts (Table S11) and our snATAC-seq data, and identified ∼370K enhancer-gene pairs in each cell type (Table S12). We found that 43-49% of the “co-activation”-based OCR-gene pairs (*FDR* ≤ 0.05) overlapped with ABC enhancer-gene pairs, representing a >2.5-fold enrichment (Fisher’s exact test *P* < 2.2 x 10^-16^) (Fig. S13D).

For most gene expression clusters, we observed concordant epigenome-transcriptome changes (Fig. S13A-B). However, several clusters showed notable differences. For ERG cluster C6, while gene expression dropped at 6 h, nearby chromatin regions remained largely open, suggesting “epigenetic memory” (Fig. S13A) (*19*). Similar epigenetic memory has been observed in mouse neurons in an early study: ∼36.5% of the activity-induced OCR peaks at 1 h remained open at 4 h while ∼5% remained open at 24 h despite the diminished gene expression (*21*). For LRG clusters C8 and C15, while gene expression peaked at 6 h (for C8, only in Glut), chromatin activation occurred at 1 h, suggesting “chromatin priming” (Fig. S13A). Such “epigenomic memory” and “chromatin priming” was exemplified at *FOS* locus, where nearby OCRs remained open at 6 h despite the diminished *FOS* expression while the OCRs were already open at 0 h without *FOS* expression (Fig. S11A). Similar chromatin priming has also been reported for some cell-type-specific enhancers in response to neuronal activation (*19*) or during neuron differentiation in which TF-binding sites exhibit a gain of chromatin openness prior to the expression of their associated genes (*46*). To further characterize the chromatin state of “primed” chromatin, we annotated KCl-responsive OCRs for neuronal active enhancer marked by H3K27ac (*29*) (Supplementary text). About 28% of KCl-responsive OCRs were found “primed”, meaning partially accessible without H3K27ac mark at 0 h, and then became fully marked by H3K27ac upon stimulation, likely enabling the access to TF-binding that fully activated the OCR (Fig. S14A-B). These epigenome-transcriptome “discordances” expanded our observation earlier for some known ERGs and LRGs (Fig. 1J), adding to the growing picture of the complexity of epigenome regulation (*47, 48*).

To mechanistically understand the complex pattern of epigenome regulation of neuron activation, we identified putative TF regulators of early and late response. These regulators were defined based on their motif enrichment patterns and differential expression during neuronal activation (Supplementary text, Table S13). We identified 145 candidate TF regulators of early response, half of which are shared by all cell types (Fig. S14C). While the expression of the shared TFs like *FOS* and *NPAS4* often elevated transiently at 1 h, their motifs remained enriched at 6 h (Fig. S14D), suggesting that the epigenomic state of these early response TFs were maintained at a later stage. Late response TF regulators showed a very different pattern: only 6 (out of 64) late response TFs were shared across cell types (Fig. S14D-F). Some GABA-specific TFs with highest motif enrichment at 6 h, including TCF4, a master regulator in SCZ (*49*), showed high expression and strong motif enrichment even before stimulation (Fig. S14D-F).

### Gene regulatory network inference of neuronal activation shed light on ASD genetics

Leveraging our sn-Multiomics data of the 18 lines (sequencing batch 24), we reconstructed gene regulatory networks (GRNs) that modulate neural transcriptional response (Methods) (*50*). Briefly, for each DEG at one condition, we defined candidate TF regulators, based on the presence of their motifs (*P* < 1 x 10^-5^ based on ArchR) in the OCRs linked to that gene (Fig. 2D). Among these candidate TFs, we then correlated the motif activity of these TFs with the target genes’ expression across pseudotime (Spearman’s *r* > 0.5), one cell type at a time. Our GRN inference resulted in 198 TFs, each having 100 or more targets in at least one cell type; the top TFs included well-known early response TFs, such as FOS and JUNB (Fig. S14G, Table S14).

These GRNs provide a framework to understand the functions of TFs in neuronal response and disease relevance. We illustrated the use of GRN in studying genetic regulation of ASD that has many known risk genes (*n*=185), including eight TFs (*51*). We focused on four TFs that likely played a role in neuronal response (Table S13): *MEF2C*, an early and late response TF in all cell types; *MLXIP*, an early response TF in all cell types; *RORB*, an early response TF in GABA and late response TF in npglut; and *TCF4*, a late response TF in GABA. These four TFs regulate 199-956 genes across cell types (Table S15). GO enrichment analysis of their target genes revealed ASD-relevant biological processes, such as synaptic transmission and neural development (Table S16). In the case of *RORB*, the enriched GO also included “lipid droplet (LD) formation” (Table S16). These results suggest convergent as well as distinct processes regulated by these TFs.

To further explore the functional relevance of these TFs, we created an “ASD subnetwork” consisting of the 4 TFs and 40 ASD risk genes that were targets of at least one TF (Fig. 2E). This network highlighted extensive cell-type-specific cross-regulation of ASD risk genes by the four TFs. For examples, *MEF2C* regulated 26 ASD genes some of which were coregulated by other TFs, while *UBR1*, a ubiquitination gene, was regulated by *MEF2C* and *TCF4* (Fig. 2E). Indeed, the shared target genes (not limiting to known ASD genes) of the 4 TFs were enriched for GO terms related to ubiquitin conjugating enzyme activity (Table S17), highlighting the importance of ubiquitin function in ASD.

To empirically validate the ASD subnetwork (Fig. 2E), we generated loss-of-function alleles for *RORB*, *MEF2C*, and *TCF4* by introducing a premature stop-codon in each TF using CRISPR DNA base editing in two donor iPSC lines (Methods). We then performed scRNA-seq of iPSC-derived Glut/GABA co-cultures upon KCl-stimulation to examine whether the predicted ASD target genes of a TF (Fig. 2E) were dysregulated by TF-knockout (KO) (Fig. 2F, Fig. S15A-M). We found that the predicted direct ASD target genes of all three TFs, RORB in GABA, MEF2C in Glut, and TCF4 in GABA, were enriched in DEGs (TF-KO vs. wild type) of their matched cell types (Fig. 2F, Fig. S15J-M). The strongest enrichment was found with *RORB*-KO, with 5/7 predicted ASD targets reduced by TF-KO in GABA at 6 h of stimulation (Fig. 2G-H). *UBR1*, the predicted target of both MEF2C and TCF4 (Fig. 2E), showed reduced expression in both *MEF2C*-KO and *TCF4*-KO (Fig. S15J-M). These functional results support the validity of our reconstructed ASD subnetwork.

To infer additional TFs important for ASD, we tested each TF’s targets for the enrichment of ASD risk genes. This analysis identified 21 TFs enriched (FDR ≤ 0.05) in at least one cell type (Table S18), most of which were Glut-specific (Fig. 2I, Fig. S14G). This list included the ASD risk gene *MEF2C* in the validated ASD subnetwork (Fig. 2E-F) and several important early response TFs like FOS. Notably, this list also included sterol regulatory element-binding proteins (SREBF1, SREBF2), which are important for regulating lipid and cholesterol synthesis (*52, 53*). Together with the enrichment of GO term “LD formation” among targets of RORB, an ASD risk gene (Fig. 2E), these results supported a possible link between lipid/cholesterol-related processes and ASD.

### eQTL mapping revealed stimulation-specific effects of genetic variants

Genetic variations associated with neuron activity-dependent expression are unknown. With the full iPSC cohort, we mapped eQTLs for each context (cell type x time point). We identified 1,316 to 4,113 genes with at least one eQTL (eGenes) across contexts (Fig. 3A, Table S19). As expected, eGenes were highly neuronal subtype-specific (30-58%) (Fig. S16A-C). About 8-11% and 15-18% of eGenes across contexts were also in vivo ERGs and LRGs (*21*), respectively (Fig. S16D-E, Table S19). The numbers of eGenes from stimulated conditions were generally larger than those from 0 h (Fig. 3A). Indeed, large fractions of eQTLs were identified only in stimulated states (Fig. S16F-J, Fig. S17), highlighting the value of using stimulation to reveal genetic effects that would otherwise be missed (*54*).

**Fig. 3.**
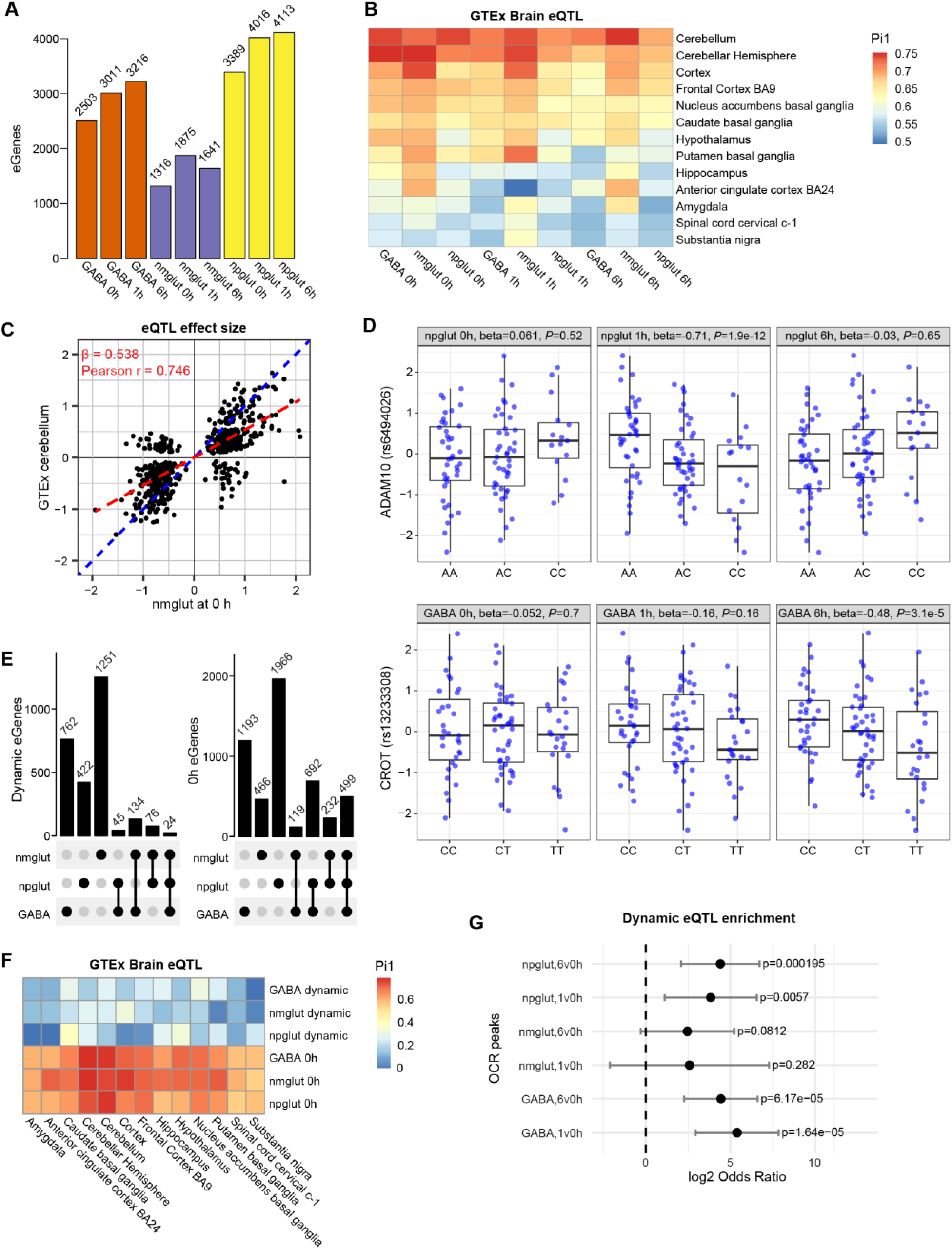
eQTL mapping identified neuronal stimulation-dependent effects of genetic variants on expression. **(A)** The number of eGenes of each context (cell type x time point). **(B)** Proportion of neuronal activity eQTL shared with GTEx brain eQTL from Pi1 analysis. **(C)** Effect size concordance of eQTL between 0 h nmglut and GTEx cerebellum. Each dot, an eQTL. Red line, fitted line with intercept=0; blue line, slope=1. **(D**) Examples of dynamic eQTL. eQTL of ADAM10 (top panel) is only significant in npglut at 1 h. eQTL of CROT (bottom panel) is only significant in GABA at 6 h. **(E)** The number of dynamic eGenes (left) and those in unstimulated neurons (0 h) (right) across cell types. **(F)** Proportion of shared neuron activity eQTL in GTEx eQTL from Pi1 analysis. Top 3, eGenes from dynamic test; bottom 3, eGenes from neurons at 0 h. **(G)** Enrichment (from TORUS analysis) of upregulated OCR peaks in dynamic eQTL.

We compared our eQTLs to GTEx brain eQTLs (*55*). Pi1 analysis showed stronger sharing (>0.65) with GTEx data for cerebellum, cerebellar hemisphere, and cortex than other regions, with strongest sharing found with 0 h eGenes (Fig. 3B). This suggested neuronal stimulation allows detection of eGenes that may be missed in postmortem brains. Overall, 19-61% of our eGenes were shared with GTEx brain eGenes (Fig. S16I), which showed a strong correlation of the effect sizes (Pearson’s *r* = 0.71∼0.76) (Fig. 3C, Fig. S18A). As expected, we observed lower correlation of the effect sizes of our eQTLs with GTEx whole blood eQTLs (Pearson’s *r* = 0.50∼0.59) (Fig. S18B). The proportion of GTEx brain eQTLs sharing effect directions with our eQTLs was substantially higher than that of GTEx blood eQTLs (Fig. S16J). These results thus supported the validity of our eQTLs.

We also compared our eQTLs with other brain eQTL datasets: PsychENCODE2 (*56*), ROSMAP (*57*), and a single-cell eQTL study of iPSC-derived dopaminergic neurons (*58*). These comparisons revealed largely similar patterns to GTEx (Fig. S19A-D): the effect sizes of our eQTLs show modest to high correlations (0.5-0.8) with others’ effect sizes. The proportions of shared eQTLs between our study and others range from 0.3 to 0.8 (Fig. S19A-D). These comparisons further support the validity of our neuron activity-dependent eQTLs and their complementarity to the existing brain and neuronal eQTLs.

To determine the “dynamic eQTLs” showing different effect sizes upon neuronal stimulation, we focused on all eGenes with at least one eQTL combining all 9 conditions (Methods). To assess the difference of eQTL effect size between 0 and 1 h or 6 h, we performed interaction testing. We identified 288 to 976 dynamic eGenes across cell types (Fig. 3D, Table S20). Compared to “static eQTLs” at 0 h, dynamic eQTLs were more likely to be cell-type-specific (Fig. 3E). We next compared the extent of overlap of our neuronal eQTLs with GTEx brain eQTLs. We found that while static eQTLs showed considerable overlap with GTEx eQTLs, much smaller proportions of dynamic eQTLs were shared with GTEx (Fig. 3F). These results suggested that neuronal stimulation revealed eQTLs missed by brain eQTL mapping. To test whether dynamic eQTLs were driven by activity-dependent epigenomic changes, we assessed the enrichment of dynamic eQTLs in OCRs. We found broad enrichment of dynamic eQTLs, compared to all eQTLs, in upregulated OCRs upon stimulation in the corresponding cell types (Fig. 3G).

### Joint analysis of eQTL and GWAS identified stimulation-specific NPD risk genes

To examine whether stimulation-dependent eQTL can help map GWAS risk genes of NPD, we used our recently developed method causal-TWAS (cTWAS) (*59*). It integrates the Transcriptome-wide Association Study (TWAS) (*60, 61*) with genetic fine-mapping, and computes the Posterior Inclusion Probability (PIP), the probability of a gene/variant being causal to a disease or trait (Fig. 4A) (*59*). We first applied cTWAS to our eQTL data at each cellular context to derive PIP of each expression trait being likely causal to a phenotype, and then computed the aggregated PIPs of the same gene across all 9 contexts (gene PIP) as the probability of a gene influencing a phenotype (Methods).

**Fig. 4.**
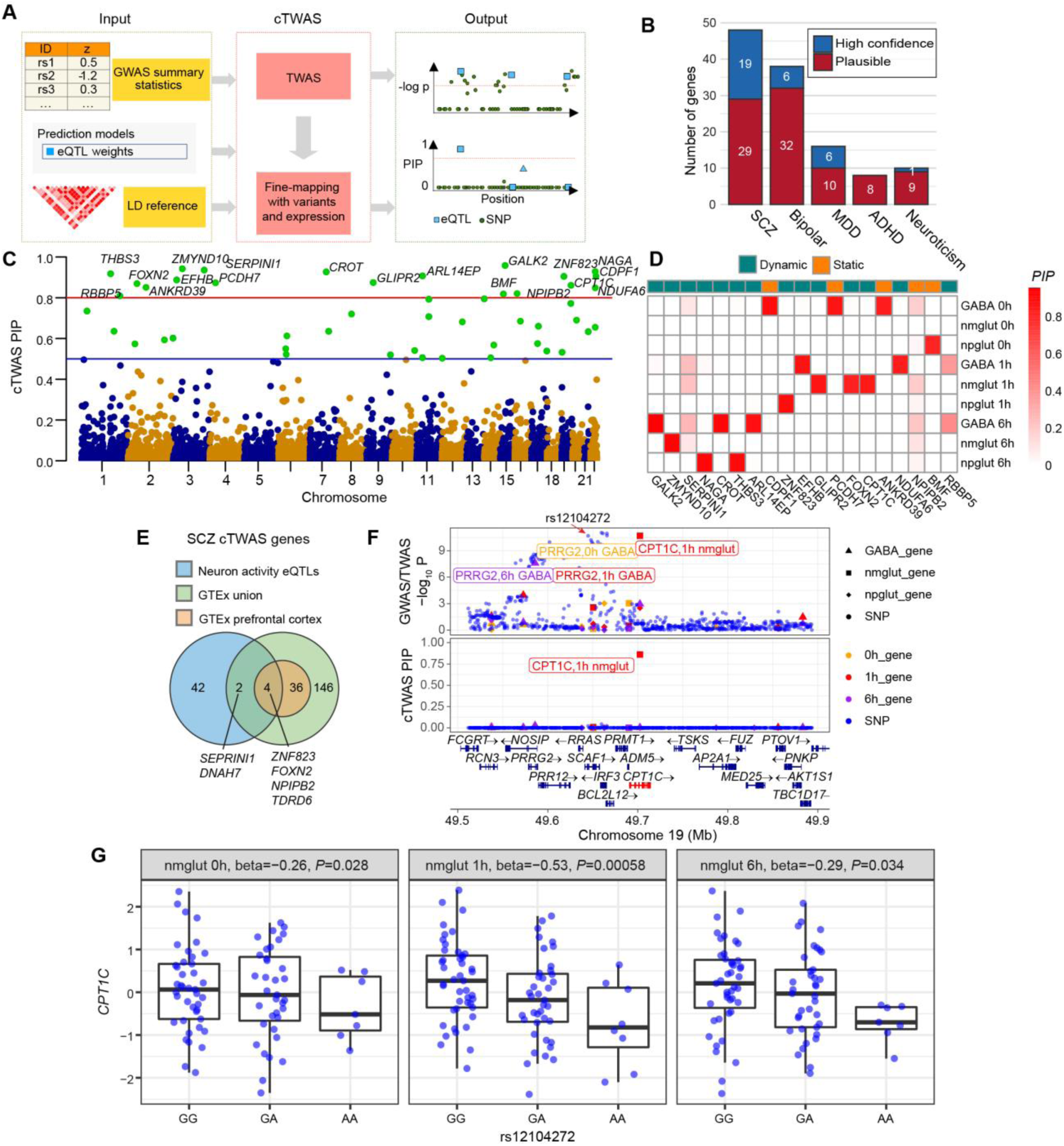
eQTL-based causal transcriptome-wide association study (cTWAS) identified putative neuronal stimulation-specific NPD risk genes. **(A)** Workflow of cTWAS analysis. PIP, Posterior Inclusion Probability of being causal. **(B)** Counts of candidate eGenes across 5 NPD phenotypes stratified by cTWAS PIP. High confidence, PIP > 0.8; plausible, 0.5 < PIP < 0.8. **(C)** Manhattan plot for SCZ cTWAS with PIP as the y coordinates and genomic position as the x coordinates. Red line, PIP = 0.8; blue line, PIP = 0.5. Highlighted are candidate risk genes of high confidence with PIP > 0.8. **(D)** PIPs across contexts for high confidence SCZ risk genes. A gene is “dynamic” if the sum of PIPs in stimulating states (1 h or 6 h) is larger than the PIP at 0 h by 0.5. **(E)** Sharing of SCZ risk genes from cTWAS between our neuronal activity eQTL and GTEx brain eQTL. **(F)** cTWAS Locus plot of *CPT1C*. Top panel, *P* values of GWAS SNPs and gene expression traits (from standard TWAS); bottom panel, cTWAS PIPs of SNPs and gene expression traits. SNPs shown as dots, and eGenes as other shapes. Red arrow points to the eQTL SNP rs12104272 (also a leading GWAS SNP) for *CPT1C*. **(G)** Box plot of *CPT1C* expression in nmglut with different genotypes of the eQTL SNP rs12104272 across time points. Note the larger estimated genetic effect at 1h than 0h or 6h (*P* = 0.08, likelihood ratio test).

We found that SCZ showed the largest number of candidate genes (Table S21), with 48 genes at PIP > 0.5 (Fig. 4B-C). Focusing on the 19 high confidence (PIP > 0.8) SCZ genes (Fig. 4C), we assessed which context drive the gene PIPs. We found that PIPs are highly context-specific, with driving contexts often being stimulated states (Fig. 4D). To better quantify this trend, we classified a cTWAS gene as “dynamic” if the total PIP from the stimulated states is larger than the PIP at 0 h by 0.5, and “static” otherwise. We found the majority of cTWAS genes for SCZ and other NPD were “dynamic” (Fig. 4C-D, Fig. S19E).

We compared our cTWAS results for SCZ to those from using the GTEx brain eQTLs. At PIP > 0.5, we found only 4 shared genes (*FOXN2*, *NPIPB2*, *TDRD6*, *ZNF823*) with cTWAS results from prefrontal cortex (Fig. 4E). Both *FOXN2* and *ZNF823* are credible SCZ risk genes (*7, 62, 63*). Even with cTWAS genes from all GTEx brain tissues, only two more genes (*DNAH7*, *SERPINI1*) were shared (Fig. 4E). These results highlight the utility of stimulation-driven eQTLs in discovering risk genes.

To understand the biological relevance of the identified cTWAS genes, we performed GO enrichment analysis on the union of candidates (PIP > 0.8) for SCZ, BP, and MDD. The top GO included “amino acid betaine metabolic processes”, “carnitine metabolic process”, and "nervous system development” (Table S22). The amino acid metabolic processes term (q = 0.01) was driven by two genes for SCZ, *CROT* (PIP = 0.86) and *CPT1C* (PIP = 0.93), with the latter driven by its eQTL at 1 h of stimulated nmglut cells (Fig. 4F). Indeed, the eQTL (rs12104272, also a leading SCZ GWAS risk SNP; Fig. 4F) of *CPT1C* driving the cTWAS results showed larger effects at 1 h vs. other time points (Fig. 4G). *CPT1C* plays an important role in neuronal lipid metabolism to maintain synaptic activity endurance (*64*). These results are reminiscent of our earlier findings that suggested lipid metabolism as a key process activated during neuronal stimulation (Fig. 2B, Cluster 6) and in ASD gene network (RORB targets, Table S16), further implying possible lipid dysregulation in NPD.

### CaQTL mapping uncovered abundant neuronal activation-dependent regulatory variants

Chromatin accessibility of cis*-*regulatory sequences controls gene expression and is influenced by individual genetic variation. Using the full iPSC cohort, we carried out caQTL mapping to identify genetic variants associated with chromatin accessibility for each context. We found 1.8∼11.6K peaks with at least one caQTL (cPeaks) in *cis* within 25 kb of a SNP (Fig. 5A, Table S23). Using a statistical interaction test, we found that 5,461 to 9,832 caQTLs were dynamic, showing different effect sizes in 1 or 6 h than 0 h (Table S24). We identified substantially more (>2-fold) cPeaks upon neuronal stimulation (Fig. 5A). The effect sizes of our caQTLs were well-correlated with PsychENCODE2 (*56*), with slightly weaker correlation for 1 or 6 h caQTLs (Fig. S20A), suggesting that the stimulation-specific effects may be missed by brain caQTLs. As another way of mapping genetic variants affecting chromatin accessibility, we analyzed allele imbalance of chromatin accessibility, or allele-specific open chromatin (ASoC), of heterozygous variants as described (*65, 66*) (Methods). We identified 4.9∼18.7K ASoC variants across 9 contexts, with more ASoC variants under stimulation (Fig. 5B, Figs. S20B, 21A-C; Table S25). Because ASoC analysis used intra-individual allelic differences while caQTL analysis was based on inter-individual variations, we assessed the agreement between the two analyses. We observed highly correlated effect sizes between the two (Pearson’s *R* = 0.82) (Fig. 5C, Fig. S20B), and top dynamic caQTL often showed strong stimulation-specific ASoC (Fig. 5D, Fig. S21D), supporting the consistency of the two analyses. These results highlight the dynamic nature of variant effects on chromatin accessibility.

**Fig. 5.**
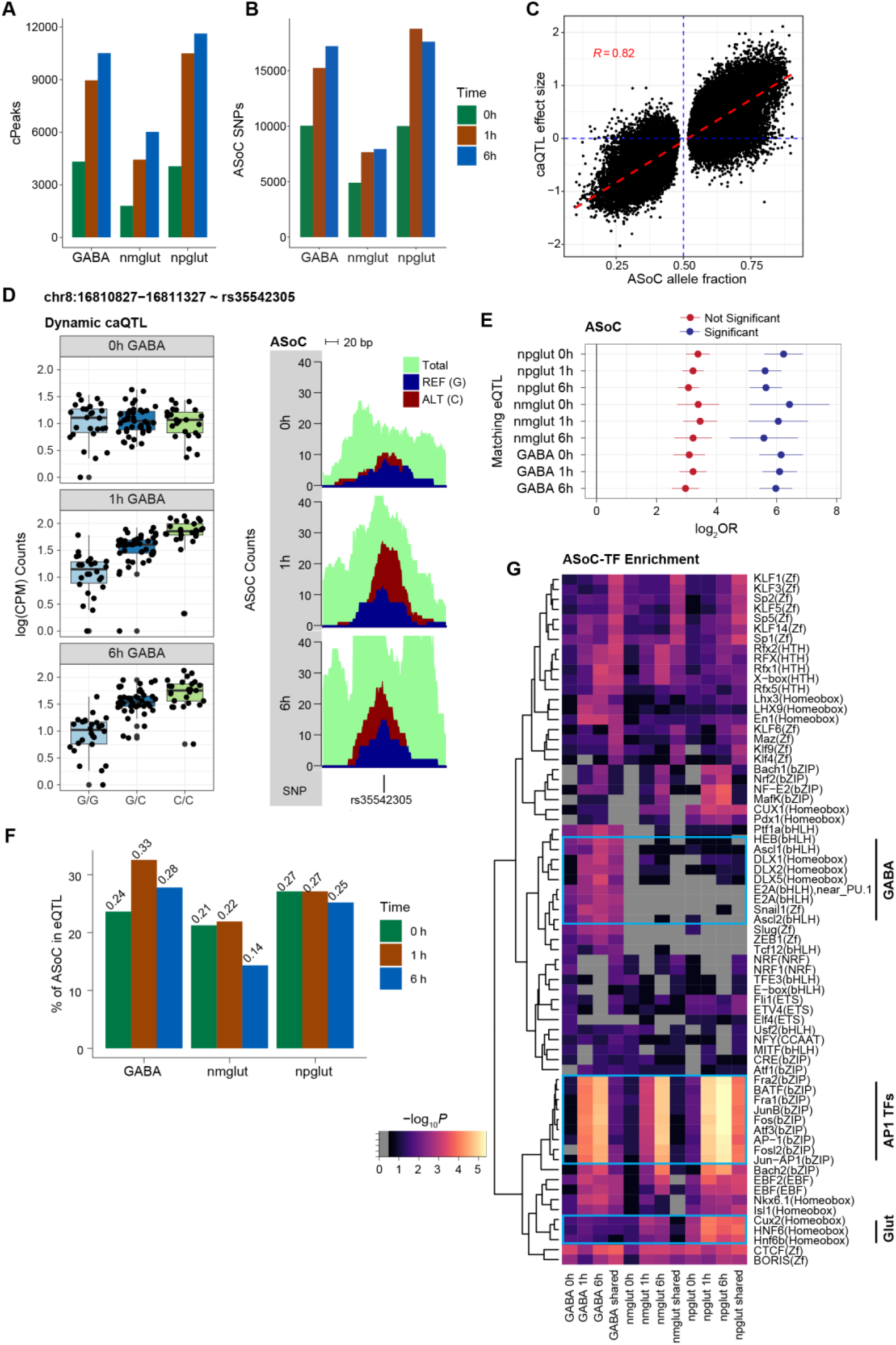
caQTL mapping uncovered neuronal stimulation-specific regulatory variants. **(A)** Counts of cPeaks (caQTL peaks) by context (cell type x time point). **(B)** Counts of ASoC SNPs by context. **(C)** Comparison of ASoC allele fractions and caQTL effect sizes across nine contexts. Red line, fitted slope; Pearson’s *r* = 0.78. **(D)** A dynamic caQTL (rs35542305) for chromatin accessibility at chr8:16810827−16811327 in GABA (left) shows consistent allelic imbalance (ASoC, right). Each dot (left panels), a cell line; CPM, count per million sequencing reads. **(E)** Fold enrichment of ASoC in eQTL in each matched context using TORUS analysis. FDR < 5% as a cutoff for significant ASoC. **(F)** Non-null proportion (Pi1) of ASoC SNPs in eQTL from the match context. **(G)** Heatmap shows the enrichment of TF motifs in ASoC SNP-flanking sequences (+/- 25 bp) at each context.

We next tested whether caQTLs likely influenced gene expression. We first assessed the enrichment of caQTL variants (for simplicity, ASoC variants were used) in eQTLs. We found strong enrichment (30 to 80-fold) of ASoC variants in eQTLs of the matching context (Fig. 5E). Our Pi1 analysis showed that 15-30% of ASoC variants were likely eQTLs in the matching contexts (Fig. 5F). Further comparison to brain eQTLs showed that ASoC SNPs were strongly enriched for frontal cortex eQTLs in both GTEx and PsychENCODE datasets (*55, 67*) (Fig. S22A-C), with 0 h ASoC SNPs showing the largest overlap with brain eQTLs. Finally, as we previously described for functional assay of ASoC SNPs in iPSC-derived neural progenitor cells (*65*), we conducted reporter gene assay of 6 ASoC SNPs (in GABA at 6 h) associated with SCZ or BP. Four of them showed differential allelic effect on reporter expression (Fig. S22D-K). These results support that caQTL variants likely affect gene expression.

To assign putative *cis*-target genes of ASoC SNPs, we examined whether an ASoC SNP was located inside a promoter or a promoter-interacting OCR in our Micro-C dataset (Fig. S13D, Table S10). With the 79-89K interacting bins across time points (Table S11), 24-44% of ASoC SNPs could be assigned to one or more target genes (Fig. S22L, Table S26). We next examined whether stimulation-specific ASoC SNPs were more likely in enhancers or promoters and enriched for specific TF-binding sites. Using GREAT (*68*), we found that stimulation-specific ASoC (vs. static ones) were more enriched in enhancers (vs. promoters) (Fig. S23). To identify specific TFs that might drive ASoC in each context, we examined the TF-binding motif enrichment at ASoC SNP sites. We found cell-type-specific TF enrichment, such as ASCL1 and DLX1/2/5 for GABA neurons and CUX2 for Glut neurons (Fig. 5G). The enriched TFs clearly distinguished stimulation phase: early response TFs such as FOS and JUNB were strongly enriched in ASoC SNPs at 1 and 6 h but not at 0 h (Fig. 5G). These results suggest that cell-type and activation-specific TF-binding in enhancers/promoters may drive context-specific caQTLs.

### Activity-dependent caQTLs explained NPD heritability missed by eQTLs and static caQTLs

To assess the role of activity-dependent chromatin accessibility variants in NPD genetics, we started with enrichment analysis of GWAS signals in ASoC variants. For each cell type, ASoC SNPs at 1 or 6 h of stimulation generally showed stronger enrichment (for SCZ, BP, MDD, neuroticism) than 0 h ASoC SNPs (Fig. 6A), suggesting stimulation helps uncover functional risk variants. To assess if neuron stimulation could help prioritize functional GWAS risk variants for SCZ (*7*), BP (*3*), and MDD (*4*), we intersected ASoC SNPs to GWAS variants and estimated the number of disease loci whose lead SNPs or linkage-disequilibrium proxies (r^2^>0.8) overlapped with at least one ASoC SNP. We found that stimulation substantially increased the number of GWAS risk loci that have risk SNPs showing ASoC (from 23 to 55 for SCZ, 20 to 38 for BP, and 45 to 100 for MDD) (Fig. 6B, Tables S27-29). Compared to GWAS loci with ASoC SNPs specific to or shared with 0 h, GWAS loci with stimulation-specific ASoC SNPs were more likely to overlap between SCZ, BP, and MDD (2.9-fold enrichment, two-tailed Fisher’s exact test *P*-value = 0.04) (Fig. S24A). *DRD2* was one of the two overlapping loci across NPD (Fig. S24A), consistent with the notion that *DRD2* is a pleiotropy hotspot for NPD (*69*). For the ASoC SNPs that might be the functional GWAS risk variants, many (73% for SCZ, 63% for BP, 68% for MDD) could be assigned to a Micro-C ci*s*-target gene (Tables S27-29). Thus, our caQTL mapping, especially the activity-dependent ASoC, substantially increased the putatively functional GWAS risk variants of NPD.

**Fig. 6.**
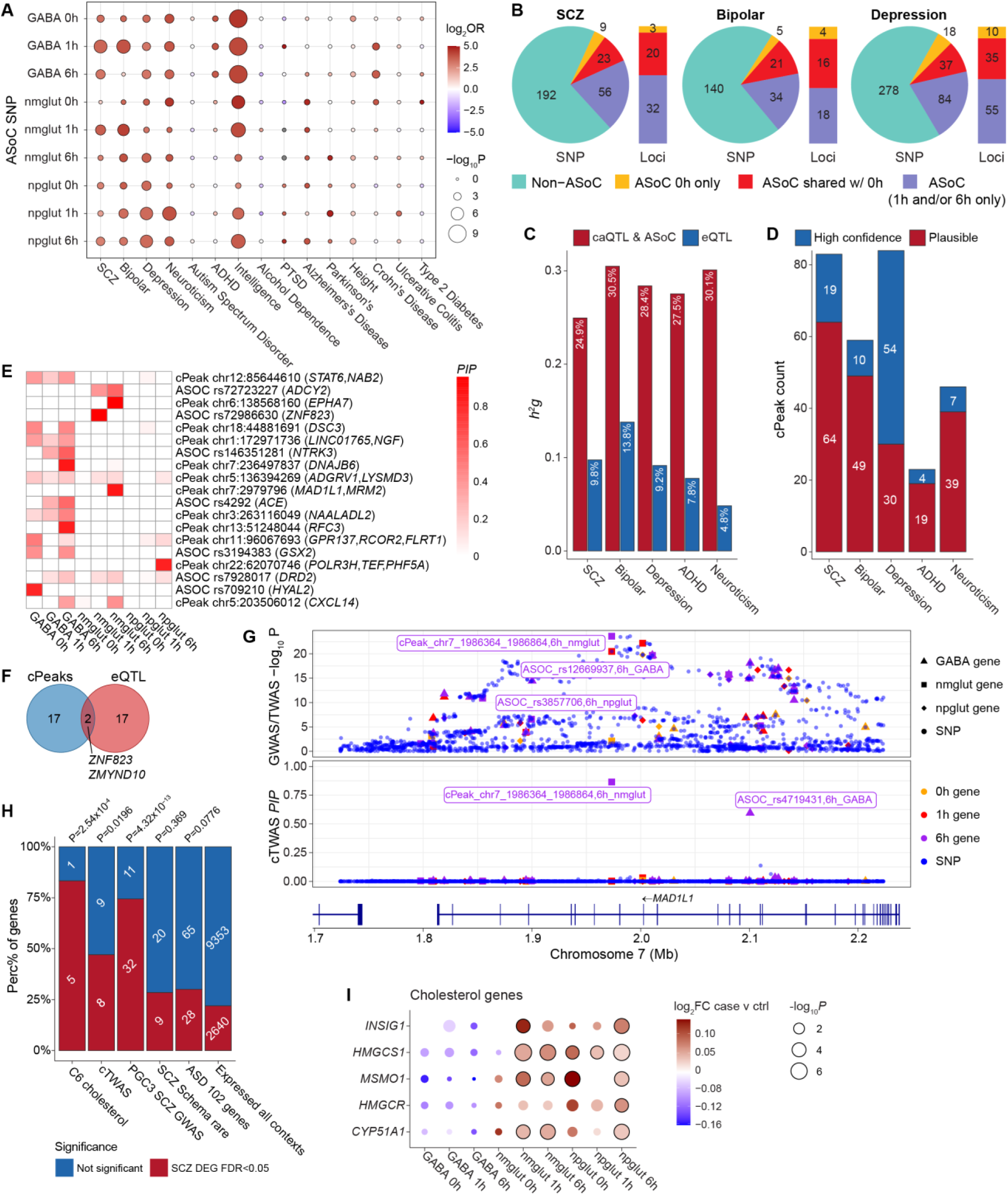
Activity-dependent caQTL explain genetic risk of NPD. **(A)** TORUS analysis of GWAS enrichment for ASoC SNPs of each context (vs. background SNPs). **(B)** Number of GWAS index SNPs and their linkage-disequilibrium proxies (R^2^>0.8) that also show ASoC (or not) for SCZ, BP, and MDD. The vertical bars show the number of putative functional GWAS risk loci that can be potentially explained by ASoC at 0 h or upon stimulation. **(C)** Comparison of h^2^_g_ explained by caQTL and eQTL for each phenotype. **(D)** Causal cPeak counts from cTWAS for each phenotype, stratified by PIPs. **(E)** Distribution of the PIPs of high confidence cPeaks across contexts. Note the predominant contribution of PIP from the stimulated conditions. Target genes of a cPeak are defined as the nearest gene (TSS) and/or eGene for the caQTL SNPs. **(F)** Venn diagram showing the overlap between high-confidence cTWAS eGenes and cPeaks. Overlap is defined by eQTL SNPs within 500 kb range of the caQTL SNPs. **(G)** Locus plot of cTWAS cPeak (chr7:1986364-1986864) in *MAD1L1*. Top panel, *P* values of GWAS SNPs and peak accessibility traits (from standard TWAS); bottom panel, cTWAS PIPs of SNPs and cPeaks. SNPs: dots, cPeaks: other shapes. **(H)** Gene set enrichment (vs. expressed in all contexts) for SCZ-associated DEGs of all contexts. Fisher’s exact test. **(I)** Cholesterol genes (in module C6 of Fig. 2B) showed SCZ-associated DE (FDR<0.05, circled). Bubble plot shows the log_2_FC and -log_10_*P* of DE from MAST test.

We then systematically analyzed the contribution of caQTLs, including ASoC, to genetic risk of NPD using cTWAS (*59*) (Methods). Here, we considered a cPeak (chromatin peak associated with caQTL or showing ASoC) as an analysis unit, similar to eGene in our eQTL-based cTWAS. For a total of 15-38K cPeaks across contexts (Table S30), we used the top caQTLs or ASoC variants (based on *p*-values) for each cPeak as the prediction model of that peak in cTWAS. We found that cPeaks explained considerably larger proportions of NPD heritability than eQTL (25% for cPeaks vs. 10% for eQTLs for SCZ) (Fig. 6C), even after controlling for eQTL effects in cTWAS (Fig. S24B). This highlighted the important contributions of genetic variants acting on epigenomes during neuron activation to NPD risk.

Given our observed larger contribution of caQTLs (vs. eQTLs) to NPD risk (Fig. 6C), we next used cTWAS to identify “causal” cPeaks. We found 4 to 54 high-confidence (PIP > 0.8) causal cPeaks across NPD (Table S30), with 19 for SCZ. Using a relaxed PIP (> 0.5) gave a larger number of cPeaks (83 for SCZ) (Fig. 6D). To identify the cellular contexts driving these results, we partitioned the PIPs of SCZ cPeaks across 9 contexts. In most cases, PIPs were contributed by stimulated conditions (Fig. 6E, Fig. S24C). Comparing with the caQTL-based cTWAS results for SCZ with earlier eQTL-based results, we found only 2 out of 19 cPeaks at PIP > 0.8 (14 out of 83 with PIP > 0.5) were shared (Fig. 6F, Fig. S24D). One of the two overlapping genes, *ZNF823,* is a likely SCZ risk gene (*7, 62, 63*). These results suggest that caQTLs, especially activity-dependent ones, complement eQTLs for identifying risk genes.

Lastly, we examined the functional relevance of the cPeaks. We linked cPeaks to their putative target genes through several means: co-activation of peaks with gene expression, eQTL targets of the caQTL or ASoC variants, genes overlapping with the cPeaks, and ABC scores computed from ATAC-seq and Micro-C data. Focusing on the potential eQTL target genes of cPeaks (FDR < 0.2 for eQTL), we found 7 out of 19 cPeaks of SCZ (PIP > 0.8) were linked to at least one target gene, with five linked to a unique gene (Table S30). Except for one lncRNA gene, the other four (*MAD1L1, ZNF823, STAT6, ADGRV1*) are all plausible SCZ candidate genes (Fig. 6G, Fig. S24E). *MAD1L1* is a known SCZ risk gene (*7*) and plays a role in neurogenesis (*70*). The cTWAS result for *MAD1L1* was largely driven by a single cPeak controlled by a caQTL of nmglut cells at 6 h (Fig. 6G). *STAT6* is important for neuroinflammation, learning and memory (*71, 72*), while *ADGRV1* is a risk gene for several brain disorders (*73*).

### SCZ risk genes and cholesterol metabolism genes were differentially activated in SCZ patient-specific neurons

Leveraging our sizable SCZ iPSC cohort (*n*=28), we examined whether the SCZ-relevant gene sets highlighted above (SCZ GWAS risk genes, cTWAS genes, or cholesterol metabolic genes in C6 cluster) showed differential neuronal activation in SCZ cases. We carried out single-cell DEG analysis in neurons between sex/age-matched SCZ cases and controls (Fig. S25A, Table S1). We identified 753 to 1,753 SCZ-associated DEGs (*FDR* < 0.05) in each neural subtype, of which 59-61% were activity-dependent (Fig. S25B-D, Table S31). GO enrichment analysis showed that activity-dependent DEGs were strongly enriched in GO related to axon guidance, axonogenesis, and synaptic transmission (Fig. S25B-D). Cholesterol biosynthetic process is among the most enriched GO terms in npglut (Fig. S25D), which is consistent with our observed enrichment of cholesterol genes in the SCZ-relevant C6 module (Fig. 2B-C). Indeed, 5 of 6 cholesterol genes in the C6 module exhibited higher expression in npgut and/or nmglut of SCZ, representing a 3.8-fold enrichment (Fig. 6H-I). CaQTL-based cTWAS SCZ genes (Fig. 6E) and the prioritized SCZ GWAS genes (Table S5) were also enriched (2.1 and 3.4-fold, respectively) for SCZ-associated DEGs (Fig. 6H, Fig. S25E-F). In contrast, SCZ or ASD risk genes from rare variant analysis (Table S5) did not show enrichment of SCZ-associated DEGs. Moreover, many genes, including those cholesterol-related ones, only showed SCZ-associated differential expression upon neuronal activation (Fig. 6I, Fig. S25E-I). The observed GO enrichments (Fig. S25B-D), including the top-ranking SCZ-associated cholesterol synthesis genes (Fig. 6I), were also supported by pseudobulk-based DEG analysis (Fig. S26A-D), highlighting the importance of context-specific regulation of SCZ risk genes and potential lipid/cholesterol dysregulation in SCZ.

## Discussion

Leveraging sn-multiomics of a large human iPSC cohort, we characterized the genetic control of individual variations of neuronal response to KCl stimulation. We found both shared and cell-type-specific TFs worked together, possibly through regulatory cascades, to drive cell-type-specific neuronal responses to stimuli. We identified thousands of eGenes and cPeaks with QTLs (including ASoC variants) in unstimulated or KCl-stimulated neurons, and found stimulation substantially increased the number of eGenes and cPeaks. Compared to the QTLs from unstimulated neurons, stimulation-specific QTLs shared less with postmortem brain QTLs and enabled discovery of more risk genes for NPD. While acknowledging the difficulties in precisely disentangling biological context-specificity from stimulation-induced detectability, we showed iPSC models robustly revealed stimulation-specific QTLs otherwise inaccessible or obscured in human brains.

Our large cohort enabled us to characterize genetic variation of activity-dependent gene expression and chromatin activity relevant to NPD. We found that eQTLs upon stimulation showed stronger enrichments of NPD heritability than baseline conditions, and most candidate genes were identified under the stimulation contexts. Compared to the eQTL results, caQTL-based cTWAS identified more putatively causal cPeaks upon stimulation. Although cTWAS effectively controls false discovery rate (*59*), our results should still be interpreted as statistical prioritization rather than definitive causal inference. Functional validation would be necessary to establish causality for the prioritized NPD genes. Nonetheless, these results suggest that many NPD risk variants/genes may only manifest functional effects upon neuronal stimulation, highlighting the importance of mapping context-specific genetic variants.

There are two possible explanations for the larger proportions of NPD heritability explained by caQTLs than by eQTLs. First, gene expression regulation is considerably more complex than individual regulatory elements, involving multiple enhancers that affect transcription and other RNA-stability-regulating elements. Thus, the effect of a genetic variant on expression is likely smaller than on chromatin accessibility, making it harder to identify eQTLs than caQTLs. Second, as we observed, epigenomic and transcriptomic changes are not always synchronized, such that regulatory sequences of an ERG may remain open even after gene expression has diminished. This makes it possible to identify caQTL effects in the absence of eQTL effects. Other explanations may include differences in QTL effect size, proximity to causal variants, or regulatory buffering (*74, 75*). Nevertheless, caQTL mapping, particularly in stimulated cellular contexts, substantially complements eQTL mapping in unraveling the functional risk variants (*56, 76*).

Multiple analyses support a role of activity-dependent lipid/cholesterol metabolism in NPD. The ERG cluster C6, which showed GWAS enrichment for SCZ, was also enriched for cholesterol metabolic genes. 5 of 6 cholesterol genes in C6 were differentially expressed in SCZ neurons only upon stimulation. Moreover, SREBF1 and SREBF2, TFs with targets enriched for ASD genes, both regulate lipid/cholesterol synthesis (*52, 53*). We also observed enrichment of the GO “lipid droplets formation” among targets of RORB, an ASD risk TF. Finally, *CPT1C*, a SCZ cTWAS gene identified in stimulated neurons, regulates neuronal lipid metabolism to maintain synaptic endurance (*64*). These results align well with the existing evidence that points to possible influences of lipids dysregulation on central aspects such as neurodevelopment, synaptic function, and immune mediation in NPD and neurodegenerative disorders (*40, 77–80*). Together with previously reported increases of cholesterol in SCZ and some other NPDs (*40, 78–80*), our observed expression increase of cholesterol synthesis genes in SCZ neurons suggests that cholesterol-targeting drugs may help improve clinical symptoms of NPD (*40*).

Despite a large iPSC cohort, our QTL mapping may still be underpowered. Furthermore, although KCl-stimulated genes in our iPSC models were enriched for in vivo activity-dependent genes (*21, 37*), our model may only partially recapitulate the in vivo neural activation. Moreover, although sampling time during the day was well-controlled, some activity-dependent genes like *FOS* also exhibit brain rhythmicity (*81*), which may confound our findings. Finally, although our chosen time points of KCl stimulation are commonly used for modeling neuron activation (*21, 29, 37*), additional sampling time points may provide a more comprehensive temporal assessment of activity-dependent gene regulation. Nevertheless, our work provides mechanistic insights on neuron subtype-specific activity-dependent gene regulation, substantially expanding the repertoire of context-specific causal variants/genes for NPD and other brain traits.

## Materials and Methods

### Human iPSC lines and cell culture

We initially started with 107 iPSC lines (European ancestry) of which 100 lines were successfully differentiated into both excitatory and inhibitory neurons (Table S1). Of the 100 iPSC lines used for data production, 58 with their IDs starting with “CD” were reprogrammed at Rutgers University Cell and DNA Repository (RUCDR)-NIMH Stem Cell Center using the cryopreserved lymphocytes (CPLs) of donors of Molecular Genetics of Schizophrenia (MGS) cohort, and have been used in previous studies (*65, 66, 80, 82, 83*). The rest were purchased from the California Institute of Regenerative Medicine (CIRM). 28 iPSC lines are from SCZ cases and 72 are from healthy controls. The donors’ average age is 52 (+/-17) years old, and 56 of them are males. For TF-Knockout by CRISPR-based DNA base editing, two donor iPSC lines, CW20107 from CIRM and KOLF2.2J from Jackson Lab (JAX), were used as part of the SSPsyGene consortium (*83*). All iPSC lines were verified for their identify based on the matching of snRNA/ATAC-seq-inferred genotypes with their known genotypes. Cells were maintained in feeder-free mTeSR Plus (Stemcell # 100-0276) and passaged using ReLeSR (Stemcell # 100-0483) every 4-6 days following the vendor’s instructions. The Endeavor Health Institutional Review Board (IRB) approved this study.

### Lentivirus preparation

Lentiviral particles were prepared using low-passage 293T cells maintained in DMEM medium supplemented with 10% FBS. 24 h prior to transfection, 95% confluent 293T cells were dissociated using Accutase and replated at 1:3 ratio to achieve 70-80% confluence the next day. On the day of transfection, lentiviral plasmids including FUW-rtTA (Addgene # 20342), TetO-Ascl1-puro (Addgene # 97329), Dlx2-hygro (Addgene # 97330), pTet-O-Ngn2-puro (Addgene # 50247) were co-transfected with pMDLg/pRRE (Addgene #12251), pMD2.G (Addgene #12259) and pRSV-Rev (Addgene #12253) at 1:1:1:1 molar ratio using FuGENE HD (Promega). 18 h post transfection, DMEM with 10% FBS medium was replaced with fresh mTeSR Plus medium. 48 h post transfection, supernatant containing viral particles was collected and cell debris were removed by centrifugation at 500 x g for 5 min. Supernatant was aliquoted into low-binding tubes and stored at -80°C.

### Neuronal differentiation and co-culture

We differentiated iPSC lines into both excitatory and inhibitory neurons and co-cultured with rat glial cells. For excitatory neuronal differentiation, we used the method for deriving Ngn2-induced glutamatergic neurons (*30*) with minor modifications. On DIV (days in vitro) 0, 60-80% confluent iPSCs were dissociated into single cells using Accutase and replated into Matrigel-coated 6-well plate at 5 x10^5^ cells per well in mTeSR Plus medium with 5 mM ROCK inhibitor, together with appropriate amount of rtTA virus and Ngn2-puro virus. On DIV1, medium was refreshed with mTeSR Plus containing 5 mM ROCK inhibitor and 2 mg/ml doxycycline. On DIV2 to DIV4, cells were treated with neural culture medium (Neuralbasal medium with 1x B27Plus and 1x Glutamax) supplemented with 2 mg/ml puromycin, 2 mg/ml doxycycline to remove non-transduced cells. On DIV5 to DIV6, puromycin was withdrawn and cells were treated with neural culture medium with 2 mg/ml doxycycline. On DIV7, cells were ready for replating.

For generating excitatory neurons, we used the method for deriving Ascl1/Dlx2-GABAergic neurons (*31*) with minor modifications. On DIV0, 60-80% confluent iPSCs were dissociated into single cells using Accutase and replated into Matrigel-coated 6-well plate at 7 x 10^5^ cells per well in mTeSR Plus medium with 5 mM ROCK inhibitor together with rtTA virus, Ascl1-puro virus and Dlx2-hygro virus. On DIV1, medium was refreshed with mTeSR Plus containing 5 mM ROCK inhibitor and 2 mg/ml doxycycline. From DIV2 to DIV4, cells were treated with neural culture medium supplemented with 2 mg/ml puromycin, 150 mg/ml hygromycin and 2 mg/ml doxycycline to remove non-transduced cells. On DIV5 to DIV6, cells were treated with neural culture medium with 2 mg/ml doxycycline and 2 mM AraC. Cells were ready for replating on DIV7.

To co-culture the excitatory and inhibitory neurons, on DIV7, separately cultured Ngn2-glutamatergic neurons, Ascl1/Dlx2-GABAergic neurons, and primary rat cortical astrocytes (Thermofisher; N7745100) were dissociated using Accutase at 37°C for 15-20 min. Ngn2-glutamatergic neurons, Ascl1/Dlx2-GABAergic neurons and astrocytes were replated at 5:5:1 ratio onto Matrigel pre-coated 12-well plate in neural culture medium supplemented with 2 mg/ml doxycycline, 10 ng/ml BDNF, 10 ng/ml GDNF, 10 ng/ml NT-3, and 1% FBS. For most co-cultures, we pooled the neural cells individually differentiated from 3-5 donor lines with equal proportion. On DIV8, we refilled cells with more medium. From DIV9 to DIV33, the medium was refreshed every three days with half volume changes. Doxycycline was withdrawn at DIV14 and 1 µM AraC was added in the medium from DIV8 to DIV17 to ensure neuron purity. DIV33 neurons were used for KCl stimulation.

### KCl stimulation of neuronal cultures

To model neuronal activation, we followed a previously used protocol to treat the cells with KCl (*28*). Briefly, one day before stimulation, old medium was aspirated and DIV33 neuron co-culture were silenced overnight in neural culture medium supplemented with 1 mM TTX and 100 mM DL-AP5. The next day, we added 0.45 volume (that is, 0.45 ml for 1 ml original culture medium) of warmed depolarization solution (10 mM HEPES, 170 mM KCl, 1 mM MgCl_2_, 2 mM CaCl_2_) to initiate KCl stimulation. We prepared three conditions with different treatment durations (0 h, 1 h, and 6 h). After stimulation, cells were dissociated for processing for 10x Genomics sequencing library preparation.

### Single-nuclei multiomics sequencing library preparation

After stimulation, cells were briefly washed with 1x PBS and dissociated in Accutase at 37°C for 40 min with gentle shaking. After Accutase incubation, we pipetted cells multiple times and filtered suspension twice with 40 mm tip filter (Sigma # BAH136800040-50EA) to obtain single cells. The washed cells were subjected to fresh nuclei isolation following 10x Genomics’ protocols (CG000365 and CG000338). For each library, around 15,000 freshly isolated nuclei were loaded for GEM capture targeting 10,000 nuclei recovery. The pre-amplified DNAs (for snATAC-seq) and cDNAs (for snRNA-seq) were shipped to the University of Minnesota Genomics Center (UMGC) for sn-multiomics sequencing library construction following 10x Genomics’ standard protocol.

### Immunocytochemistry

For Immunocytochemistry, iPSCs/neurons were fixed with 4% PFA in PBS at room temperature for 15 min. After three brief washes in PBS, cells were permeabilized with 0.5% Triton X-100 in PBS for 15 min at room temperature and further blocked with 3% BSA and 0.1% Triton X-100 in PBS at room temperature for 1 h or 4°C overnight. After blocking, samples were incubated with primary antibodies diluted with blocking buffer at room temperature for 1 h or 4°C overnight. After 3 PBS washes, samples were incubated with secondary antibodies diluted in blocking buffer at room temperature for 1 h followed by 3 more PBS washes. Then samples were incubated in PBS containing 1 mg/ml DAPI (4’, 6-diamidino-2-phenylindole) at room temperature for 10 min. After DAPI staining, samples were washed once with PBS and mounted on glass slides. The images were taken by a Nikon ECLIPSE C2 confocal microscope.

### BDNF OCR peak deletion by CRISPR/Cas9 editing

For BDNF OCR peak deletion, two gRNAs flanking the targeted region were cloned into pSpCas9(BB)-2A-Puro (PX459) V2.0 (Addgene # 62988). Two iPSC lines were used for editing. For editing, 24 h prior to transfection, 60-80% confluent iPSCs were dissociated into single cells using Accutase and replated into Matrigel-coated 60 mm dish at 4.5 x 10^5^ cells per well in mTeSR Plus medium with 5 mM ROCK inhibitor. On the day of transfection, 3 mg of plasmid DNA carrying gRNA1 and 3 mg of plasmid DNA carrying gRNA2 were introduced into iPSCs using LipofectamineSTEM (Thermofisher) following vendor’s instruction at 1:2 DNA:reagent ratio. 24-48 h post transfection, cells were selected with 0.5 mg/ml puromycin; 48-72 h post transfection, cells were selected with 0.25 mg/ml puromycin. Afterwards, antibiotics were withdrawn, and cells were maintained in regular mTeSR Plus until colonies reached appropriate size for picking. After colony picking, genomic DNAs (gDNAs) from collected cell pellets was isolated using QuickExtract DNA Extraction Solution for PCR amplification. Amplified DNAs were loaded on 1% agarose gel for electrophoresis to examine fragment size. gDNAs of the colonies with confirmed peak deletion were used for Sanger sequencing confirmation. The confirmed colonies (n = 2-3) with the expected peak deletion were sub-cloned and expanded for cryopreservation. Please refer to Table S32 for gRNA and primer sequences.

### Reporter gene assay for validating ASoC SNP

Reporter gene assay was performed as described (*65*). In brief, Plenti-PGK-V5-LUC-Neo plasmid (Addgene # 21471) was digested with XhoI and SalI to remove the PGK promoter 5’ upstream of Firefly Luciferase (LUC) gene. Afterwards we inserted a single strand oligo containing PmeI cutting site and 32-bp mini promoter through Gibson Assembly to create a new vector pLenti-PmeI-minP-LUC-PGK-Neo. 151-bp single strand oligo spanning a ASoC SNP site for each allele (75-bp 5’ upstream and 75-bp 3’ downstream) was introduced into linearized vector through Gibson assembly. Then the Lenti-virus particles were prepared as described above. To infect cells, we differentiated iPSC (lines CD19 and CD43) into Ngn2 Glut and Ascl1/Dlx2 GABA neurons separately as described above. At DIV7, GABA neurons were infected with lenti-virus containing the reporter gene and the cloned oligos for each allele. At DIV8, Ngn2 Glut, Ascl1/Dlx2 GABA neurons and primary rat astrocytes were mixed at 2:1:1 ratio and replated onto Matrigel-coated 12-well plate. At DIV30, we performed KCl stimulation for 6 h as described above. RNAs were isolated for qPCR quantification of the LUC reporter level of the two alleles, normalized by Neomycin expression (encoded by the Lentivirus).

### RNA isolation and qPCR

Total RNA was isolated using RNeasy Plus Kits (QIAGEN) following vendor instructions. For qPCR, RNAs were reverse transcribed using High-Capacity cDNA Reverse Transcription Kit (Thermofisher) following vendor instruction. cDNAs were diluted with nuclease-free water at 1:10 ratio and qPCR reactions were prepared using Taqman Universal PCR Master Mix (Thermofisher). Reactions were loaded on Roche LightCycler 480 system in a 384-well white plate. The 2^−ΔΔCt^ method was used for RNA expression quantification, with GAPDH as endogenous control. Please refer to Table S32 for qPCR assay information.

### TF knockout (KO) for functionally validating ASD gene network

To functionally validate the predicted GRN of three ASD-associated TFs (TCF4, MEF2C, RORB), we used CRISPR-based cytosine base editing system to conduct TF KO by introducing premature stop codons (iSTOP) for each TF in two donor iPSC lines, CW20107 and KOLF2.2J as part of the SSPsyGene consortium (*83*). After confirming the TF KO by Western blot, we differentiated the engineered iPSC lines into NGN2-Glut and GABA neurons, co-cultured with astrocytes, and conducted KCl stimulation as described above. Cell cultures were then dissociated and subjected to 10x Chromium Next GEM Single Cell 3ʹ scRNA-seq library preparation following 10x Genomics’ standard protocol.

### Multiomics sequencing and data quality control (QC)

10x Genomics Chromium single cell Multiome sequencing libraries were sequenced at UMGC on NovaSeq S4 platform targeting pair-end (2 x 150bp) 50K reads per nuclei for ATAC library and 25K reads per nuclei for gene expression library. Briefly, after raw data collection, Illumina’s BCL2FASTQ software was used to demultiplex and assemble the fastq files corresponding to reads (R1/R2) and indices (i5/i7) for Cell Ranger ARC. The fastq files were subsequently processed by 10x Genomics Cell Ranger ARC (v2.0.2) and aligned twice to both the human GRCh38.p14 genome and a contingent of human GRCh38.p14 and mouse GRCm38 provided by 10x Genomics for efficient identification and removal of rodent astrocytes. For snATAC-seq, the per-library Transcription Start Sites (TSS) enrichment score is usually 7-9, and the Fraction of high-quality Reads overlapping Peaks (FRiP) is usually 40-80%. We excluded cells with >15% of the reads mapped to mitochondrial genes, cells with > 8,000 or < 400 number of features, and cells with > 40,000 or <500 UMI counts (filtered by 400 < nCount < 8,000 using Seurat).

### Multiomic data analyses

#### Barcode level identification of cell line identity

For each library that contained cells derived from 1-5 iPSC lines, barcode-level identification was performed separately for snRNA-seq (GEX) and snATAC-seq data with genotyping information of all donors. Briefly, the BAM files generated from GEX and ATAC assays (gex_possorted_bam.bam and atac_possorted_bam.bam) were processed by demuxlet (*84*) with known genotype information provided as .vcf files. Barcodes (cells) identified as singlets with P1 likelihood (p1LLK) < 1 x 10^-8^ were collected, and only barcodes positively identified in both GEX and ATAC assays were retained for downstream analysis.

#### snRNA-seq data analyses

snRNA-seq (GEX) data were processed by extracting gene expression data from the .h5 files generated by Cell Ranger ARC. Only barcodes (cells) confidently assigned to individual iPSC lines were retained. With Seurat 5.1.0 (*85*), To assign cell type identity, Leiden clustering was performed at the library level based on the first 30 PCs (or appropriate as determined by the elbow plot) and subsequently comparing cell clusters and their marker gene expressions (*GAD1*, *GAD2*, *SLC17A6*, *SLC17A7*, *NEFM*). The library-level gene expression matrices were subsequently collated in Seurat 5.1.0 (*85*) as one large object with cell line-specific metadata assigned. Harmony (*86*) was used to integrate the library-level gene matrices and removed sequencing batch-derived effects. Two integrated Seurat objects were generated, one for sequencing batch 24 that including libraries of 18 lines across three time points and the other for all 84 sequencing libraries of all 100 lines.

#### snATAC-seq data analyses

For snATAC-seq data, the aggregation function of Cell Ranger ARC was used to merge and generate new fragment files for MACS2-based peak calling with the CallPeaks() function in Signac (*87*) with default settings. We made two merged fragment files, one containing libraries from sequencing batch 024 (18 lines) and the other from all batches for 100 cell lines. From each fragment file, we iterated through the combinations of cell types (npglut, nmglut, GABA) and stimulation stages (0 h, 1 h, 6 h), which generated nine peak sets (cell type x time). We further generated a union peak set from sequencing batch 024 by setting CallPeaks (combine.peaks = TRUE) for peak analysis across different cell types. Finally, peaks that fell within the ENCODE blacklisted regions were removed. We also removed peaks that fell within chromosomes X and Y and the mitochondrial genome.

### Approximation of developmental stages for iPSC-derived neurons

We used the reference single-cell dataset of human embryonic and postmortem brains (*34*) to evaluate our iPSC-derived neurons and their comparable developmental stages. Briefly, the excitatory neuron (ExNeu) and interneuron (IN) populations were extracted using the identity assigned in the original publications. The excised data were then processed as documented in the original publication with Harmony-assisted data normalization, scaling, and dimension reduction using the first 30 PCs to generate the anchor gene set, UMAP dimensions, and unimodal UMAP coordinates for projection. The three main cell types identified (npglut, nmglut, GABA) from snRNA-seq were used as queries. Each cell type was firstly randomly subset to 10,000 cells to reduce computational demands. Subsequently, each cell subset was separately projected by the MapQuery() function to its corresponding cell map (npglut/nmglut to ExNeu and GABA to IN) to show their approximate developmental stages in the human brain.

### PCA analysis

For PCA analysis, we generated pseudo-bulk count matrices from merged snRNA-seq data using the AggregatedExpression() function from Seurat, and each sample represented one cell line in one of three main cell types and its stimulation stage. To further eliminate the interference from sequencing batch-specific effects, we ran regression using the ComBat-seq() function from the R package sva (*88*) using sequencing batch information as the factor. The sva-adjusted pseudo-bulk count matrices were log-transformed and estimated for their observation-level weights using the voom() function from the R package limma (*89*). Finally, we calculated the principal components (PCs) with the prcomp() function and plotted the samples using their first two PCs using the fviz_pca_ind() function from the R package factoextra.

### Pseudo-bulk RNA-seq differential gene expression analysis

SnRNA-seq data of the 18 lines from sequencing batch 24 were used. Gene differential expression (DE) analysis was performed using the R package limma. Briefly, the ComBat-seq-adjusted pseudo-bulk count matrices generated from PCA analysis were used as the initial input. Low-expression genes whose Count Per Million (CPM) value was less than < 1 in half of lines in any of the 3 time points were removed. The matrices were log-transformed and observation-level weights were calculated by voom(). Subsequently, we made the design matrix of known covariates (time, batch, age, sex, SCZ-affection status, and the percentage of corresponding cell types). We fitted the linear model using the lmFit() function of limma. After checking the mean-variance trend (SA plot) for gene distributions, the DE values (log_2_FC, *p*-value, and FDR) were calculated using the topTable() function with corresponding coefficients (1 h vs 0 h, 6 h vs 0 h, 6 h vs. 1 h) for all three cell types.

### MAGMA gene set enrichment analysis

We performed MAGMA analysis using MAGMA version 1.08b (*90*) to evaluate the enrichment for the GWAS risk of several psychiatric disorders (SCZ, Neuroticism, ASD, BP, MDD) (*3*, *4, 6, 7, 69, 91, 92*), as well as the GWAS set of Crohn’s disease that served as a control set (*93*) (Table S33). The method was adapted from our previous publications with modifications (*66*). Specifically, we compiled the MAGMA-required gene annotation data files based on the more recent GRCh38/hg38 genome to get the gene-SNP annotation file. With the gene-SNP annotation file, we then performed gene-level analysis on SNP *p*-values using the reference SNP data of 1,000 Genomes European panel (g1000_eur_hg38, --bfile) and the pre-computed SNP *p*-values from each disorder’s GWAS data set. The sample size (ncol=) was derived from either GWAS data frames or specified according to the affiliated README data. Subsequently, the result files (--gene-results) from the gene-level analysis were read in for competitive gene-set analysis (--set-annot), where we used default setting (correct=all) to control for gene sizes in the number of SNPs and the gene density (a measure of within-gene L.). The gene-set analysis produced the output files (.gsa.out) with competitive gene-set analysis results that contained the effect size (b) and the statistical significance of the enrichment of each gene set (upregulated/downregulated) for each disorder’s GWAS data set.

### Gene ontology enrichment analysis

GO term analysis was performed for the DEG sets specific to each cell type and expression pattern. Briefly, the list of DEGs was used as the SynGo (*94*) input and all the expressed genes in the corresponding cell type were used as the background gene list. A similar approach was used to calculate the enrichment of different gene expression patterns against a multitude of known NPD gene sets. For cTWAS gene sets, the enrichment analysis was conducted using the Enrichr package (*95–97*) with the default background gene set, and Benjamini-Hochberg method was used for multiple testing correction.

### Differential peak accessibility analysis

SnATAC-seq data of the 18 lines from sequencing batch 24 were used. DA peak accessibility was performed using aggregated pseudo-bulk counts based on the three cell-type-specific peak intervals (nmglut, npglut, GABA). Briefly, the AggregatedExpression() function was used to generate cell line x peaks matrices using the cell-type-specific peak intervals (approximately 300K peaks per cell type). The ComBat_seq() function from sva was subsequently applied to correct group bias for each count matrix. We removed all low-count peaks (CPM < 1 in at least half of the cell lines), and approximately 200K of the original 300K peaks survived filtration. The matrices were log-transformed and observation-level weights were calculated by voom(). Subsequently, we applied design matrices of known covariates (time, batch, age, sex, SCZ-affection status, and the percentage of corresponding cell types) as we did in DEG analysis. We fitted the linear model using the lmFit() function of limma. The DA values (log_2_FC, *p*-value, and FDR) of peaks for each cell type and contrast were calculated using the topTable() function with corresponding coefficients (1 h vs 0 h, 6 h vs 0 h).

### Stratified linkage disequilibrium score regression (sLDSC) for GWAS enrichment analysis

sLDSC (*98*) analysis was performed by using the hg38 version of European genotype data (SNPs) from 1,000 Genomes Phase 3 and v2.2 baseline linkage disequilibrium/weights. Briefly, linkage disequilibrium score estimations were pre-calculated from the hg38 version of the 1,000 Genomes EUR file set (w_hm3_no_hla.snplist), window size 1 cM (ld-wind-cm 1). We used the summary statistics of major psychiatric disorders and non-psychiatric diseases for partition heritability, with several data sets lifted over from hg19 to hg38 when necessary. Disease-specific regressions were performed independently using hm3 SNP weights against each disease for cell-type-specific analysis.

### SnATAC-Seq profiles and peak calling for caQTL and gene regulatory network (GRN) analyses

To integrate the snATAC-seq data, we utilized ArchR. Quality control measures were applied by filtering cells with a transcription start site (TSS) enrichment score of at least 4 and fragment counts of at least 1,000. After quality control and integration of expression data, we retained a total of 552,653 cells. To call peaks from the snATAC-Seq data, pseudo-bulk replicates were generated for each cell type and for each time point. To ensure equal representation of all cell types, we followed the default setting of ArchR and sampled a maximum of 500 cells and 25 million fragments from each pseudo-bulk replicate. With the pseudo-bulk replicates, 501-bp fixed-width peaks were called using MACS2 (*99*) and integrated in ArchR. Peaks were required to be reproducible in at least two samples, and sex chromosomes were excluded in the analysis. To ensure high-quality peak detection, we limited the number of peaks to a maximum of 250,000 for each cell type and time point context. Consensus peaks were obtained through an iterative overlap peak merging approach in ArchR. This method allowed for robust and reproducible peak identification across different samples and conditions.

### Single-cell embeddings

To reduce computational cost, we down-sampled 18 cell lines (115,855 cells in total) from Batch 024 for all analyses related to gene regulation, including dimensionality reduction, chromVAR analysis, pseudotime inference, peak-gene mapping, and gene regulatory network construction. Using ArchR, we performed dimensionality reduction on snRNA-Seq and snATAC-Seq data separately, utilizing the iterative latent semantic indexing approach. To create a more robust two-dimensional representation using UMAP, we combined the embeddings of expression and peak accessibility, subsequently correcting for batch effects using Harmony.

### Inference of pseudotime

Pseudotime trajectories were estimated for the GABA, nmglut, and npglut cell types separately to model the transitions between neuron states from unstimulated neurons to neurons stimulated for 1 h and 6 h. Based on the UMAP constructed in the previous section, cell-type-specific trajectories were inferred using the addTrajectory() function in ArchR. We fine-tuned the sparsity of the spline fit and quality filters for the trajectory of each cell type to avoid overfitting of smoothing spline. Cells were divided into 100 bins based on pseudotime, and the average activities were calculated by pooling cells with similar pseudotime values.

### Co-activation analysis to link OCRs to genes

The 18 lines from sequencing batch 24 were used. To link OCRs to potential target genes, we assessed the correlation of gene expression and normalized chromatin accessibility for all ORCs within 500 kb of genes. All differentially expressed genes were included in the analysis. For each cell type, peaks present in at least 5% of the cells were considered. Gene expression was regressed on peak accessibility using a negative binomial mixed regression implemented using lme4 R package with the following formula:

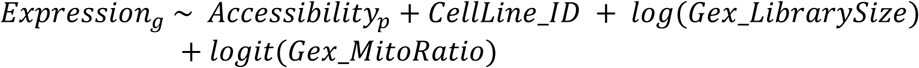

where:

- *Expression*_*g*_ represents the raw counts of gene expression for gene *g*,
- *Accessibility*_*p*_ denotes the normalized chromatin accessibility of peak *p*,
- *CellLine_ID* is a categorical variable representing cell line identity, included as a random intercept to account for donor-specific effects,
- *Gex_LibrarySize* refers to the single-cell level library size from the snRNA-Seq data, captured by the number of reads per cell,
- *Gex_MitoRatio* captures the single-cell level percentage of mitochondrial reads from the snRNA-Seq experiment, serving as a measure control for cell quality.

To control for multiple testing, the Benjamini–Hochberg procedure was applied to calculate the false discovery rate (FDR). OCR-gene pairs were considered significant and biologically relevant if they had an FDR ≤ 0.05 and a positive regression coefficient.

### Activity-by-Contact (ABC) analysis

We employed the Activity-by-Contact (ABC) model (*45, 100*) to predict enhancer-gene regulatory relationships. For our analysis, we utilized snATAC-Seq data from all 18 lines in sequencing batch 24. Additionally, we incorporated time-specific bulk Micro-C data derived from one cell line (CD-11), averaged across all cell types, which allowed us to capture the dynamic nature of chromatin interactions during different neuronal stages. Our analysis aimed to predict enhancer-gene pairs across nine distinct contexts, defined by different combinations of time and cell type. To achieve this, we called peaks for each specific context and subsequently predicted ABC scores under these conditions. An enhancer-gene pair was classified as significant if its ABC score was greater than or equal to 0.021, a threshold recommended by the software to achieve 70% recall.

### Gene cluster or module analysis

The 18 lines from sequencing batch 24 were used. To cluster genes based on their expression trajectory in pseudotime, we first selected 5,221 differentially expressed and highly variable genes, using criteria of *FDR* ≤ 0.05 and abs (Log_2_FC) > 1 in at least one differential expression test. We then normalized expression for each gene separately, so that genes with similar trajectory patterns would be clustered together, irrespective of their absolute expression levels. We then performed K-means clustering to group the genes into 15 distinct modules. Briefly, similarity in expression patterns was defined based on the scaled and normalized expression profiles of genes across 100 pseudotime points for each of the three cell types, resulting in a total of 300 data points per gene. K-means clustering was applied to group genes with similar temporal expression trajectories. The optimal number of clusters (modules) was set to 15, which provided sufficient resolution to capture distinct transcriptional dynamics while minimizing redundancy among modules with overlapping expression patterns. Gene Ontology enrichment analysis was conducted using the Enrichr package (*95–97*), and GWAS enrichment analysis was performed in MAGMA. We inferred chromatin accessibility trajectories for the 15 gene modules by mapping all peaks to the genes within each module, as described in the previous section.

### Defining candidate TFs regulating early and late responses

The 18 lines from sequencing batch 24 were used. We used chromVAR to estimate single-cell level enrichment of TF activity. With motif annotations curated from Cis-BP, we estimated chromVAR motif deviations implemented in the ArchR toolkit. We also had single-cell expression from scRNA-seq for all TFs. For each TF, we assessed its role in early and late responses by testing differential motif activity and expression between conditions, one cell type at a time. Specifically, a TF was considered a candidate TF for early response if it showed both higher motif activity and increased expression at 1 h compared to 0 h. Similarly, a TF was considered a candidate TF for late response if it showed higher motif activity and elevated expression at 6 h compared to both 1 h and 0 h. Differential expression analysis was performed using the limma-voom analysis, and differential motif analysis was performed using one-tail Wilcoxon signed-rank test using single-cell level motif deviations estimated by chromVAR.

### Gene regulatory network (GRN) analysis

The 18-line data from sequencing batch 24 were used. To construct the gene regulatory network, we followed these steps: (1) Identify Potential TF Regulators: We identified a set of potential TF regulators by selecting TFs that were differentially expressed in at least one condition. We also ensured that a TF’s motif activity (estimated by chromVAR) and its expression activity were positively correlated. Specifically, we filtered TFs based on the correlation between their expression trajectories and motif activity trajectories, retaining only those with a Spearman’s correlation > 0.3. (2) Define candidate TFs of a gene: we linked Open Chromatin Regions (OCRs) to a gene using co-activation analysis, with *FDR* < 0.1; and then examined the presence of the TF motif (*P*-value cutoff of 5 x 10^-5^) in these OCRs linked to the gene, using motifmatchr in ArchR. (3) Correlate TF Motif Activity with Target Gene Expression: For motifs present in at least one peak associated with a gene, we correlated the motif activity of TFs with the expression of the target genes. We retained the TF-gene pairs that exhibited a Spearman’s correlation > 0.5 between the motif activity of the TF and the expression of the target gene.

To infer TFs that may play roles in ASD (Fig. 2I), we used Fisher’s exact test to evaluate the enrichment of ASD risk genes in each TF’s predicted targets. The background genes of this test are all genes (expressed in at least 5% of cells) included in GRN reconstruction. FDR was used to correct for multiple testing.

### eQTL mapping

To associate the genotypic variation and gene expression variation within each context (combination of cell types and time points), we used the scRNA read count matrix of 36,601 features and 548,800 cells from 100 cell lines and three cell types (GABA, nmglut, and npglut). We first generated pseudo-bulk matrices by summing up the read counts by each feature for cells within a certain time point, a certain cell type, and a certain cell line. We removed samples with less than 1 million total reads. We also removed genes that were not protein-coding, in mitochondria, with less than 2,000 reads across all cells, or without HGNC symbol. After filtering, we retained the pseudo-bulk read count matrix of 14,818 genes by 824 samples (out of 900). Then, we performed trimmed mean of M values (TMM) normalization and inverse Normal Transformation. The resulting gene expression matrix served as input for MatrixeQTL, along with covariates of cell type composition, sex, age, SCZ affection status, 5 genotype PCs, and 15 gene expression PCs. Genotypes for iPSC donors were downloaded from dbGaP (phs000021.v3.p1 and phs000021.v3.p2 for MGS lines, and phs002032.v1.p1 for CIRM lines). For each cell line source, genotype data were processed on the TOPMed Imputation Server using a standardized pipeline that included liftover to GRCh38, quality control, phasing, and genotype imputation. Genotypes were phased with Eagle v2.4 and subsequently imputed with Minimac4 against the TOPMed all-population reference panel (GRCh38). Post-imputation variants were filtered on any missing values, imputation quality *R*^2^ < 0.3 (default setting), and MAF < 5%. Then we used the intersection of imputed SNPs from both sources (*n* = 5,966,820 SNPs). Additional 3,377 SNPs were then removed due to HWE filter (*P* < 1 x 10^-6^). We used *cis*-variants 250 kb up- and downstream of the gene body. This *cis* window size and the number of gene expression PCs were chosen to optimize the number of eGenes with at least one significant eQTL discovered. We adopted the default procedure to define significant eQTL in MatrixeQTL (*101*). In brief, we computed the *t* statistic of the genetic effect and derived the nominal *P* value for each gene-SNP pair. FDR was used to correct for multiple testing across all gene-SNP pairs. To assess if this procedure adequately controls for multiple testing, we shuffled the genotype across donors and applied MatrixeQTL procedure for each context in the shuffled data. We found less than two eGenes from the analysis, confirming that the procedure effectively controlled for false discovery rates.

In addition to the eQTL mapping in each context, we also performed eQTL mapping by aggregating all contexts. We removed the 5 cell lines from batch 22 and summed up the read counts per cell line per contexts. TPM normalization was applied to the summed reads and then we further summed up all contexts within the same cell line. We proceeded to eQTL mapping with the resulting 95 samples (cell lines) by applying TMM, INT, and MatrixeQTL. This is the same procedure as described above, except that the number of expression PCs was 21 instead of 15. eQTL results from this approach are called “pseudo-bulk eQTL” in subsequent sections.

### Dynamic eQTL testing

We tested the genetic effect heterogeneity across 3 times points, one cell type at a time. We used the following procedure to choose the candidate set of gene-SNP pairs to be tested for each cell type. We considered only eGenes found in at least one time point, at a relaxed threshold FDR < 0.2. For each of these eGenes, we started with its eQTLs with the smallest nominal *P* values per condition. Then we took a union of these top eQTLs along with the pseudo-bulk eQTLs. There are at most four eQTLs for each eGene in a cell type. We then performed LD pruning with *r*^2^ < 0.1 on these eQTLs within the same eGene.

The pseudo-bulk gene expression used for testing dynamic eQTLs were jointly normalized within each cell type. Specifically, we applied inverse Normal transformation across all the pseudo-bulk samples within the same cell type. PCA was applied to the normalized gene expression of each cell type. The first 15 gene expression PCs were used as covariates in the test below. Our interaction effect model compares the genetic effect in one stimulation context with the resting context in the same cell type. In total we have six different sets of comparison (GABA 0 h vs. GABA 1 h, GABA 0 h vs. GABA 6 h, nmglut 0 h vs. nmglut 1 h, nmglut 0 h vs. nmglut 6 h, npglut 0 h vs. npglut 1 h, npglut 0 h vs. npglut 6 h). For each comparison, we selected the gene-SNP pairs with *FDR* < 0.2 from eQTL mapping, and run the interaction effect model as following:

Gene expression ∼ Age + Sex + Disease + cell type proportion + Genotype + Time Point + Genotype x Time Point + 5 Genotype PCs + 15 Gene expression PCs

*P* values were derived from the Student’s *t*-test statistic of the interaction term of “Genotype x time point”. We used ACAT (*102*) to aggregate the *P* values of SNPs from the same gene. We then adjusted multiple testing using Benjamini-Hochberg method.

To compare dynamic eQTLs vs. static eQTLs in terms of cell type sharing and comparison with GTEx, we classified an eGene by whether it is also a dynamic eGene from the interaction testing. All other eGenes were considered static eGenes. To test if dynamic eQTLs tend to be associated with certain epigenomic features, compared to other (static) eQTLs, we directly used the summary statistics of the interaction tests. Specifically, we tested if the dynamic eQTLs are enriched in upregulated OCRs using the tool TORUS (*65, 103*). TORUS assesses if dynamic eQTLs tend to be in up-regulated peaks in the matching cell types, compared to SNPs unlikely to be dynamic eQTLs. Importantly, since only eQTLs are used in this test, the variants in the background are all eQTLs, with the majority of them being static eQTLs.

#### Comparison of neuronal eQTLs with published eQTL datasets

We estimated the proportion of shared eQTLs between two datasets using π1 analysis. For effect size correlation analysis, we used the Rb estimator (*104*). This estimator is distinct from the Spearman or Pearson correlation approach because the latter does not account for estimation errors in the eQTL effects and thus underestimates the correlation of true eQTL effects (*104*).

### TORUS enrichment analyses of eQTLs and ASoC variants

We applied a Bayesian hierarchical model (TORUS) to perform SNP-based enrichment analysis, testing the enrichment of certain features with eQTL and ASoC variants (*65, 103*). Briefly, TORUS associates the prior probability that a variant is causal to gene expression with the annotations of the variant with a logistic regression model. TORUS then uses this prior in a simple fine-mapping model (assuming a single causal variant per genetic locus) of eQTL data. TORUS uses Maximum Likelihood to estimate the parameters of the logistic regression. To assess the enrichment of ASoC in eQTLs, we applied the ASoC in matched condition as the neuronal activity eQTLs. ASoC SNPs were treated as binary annotations: 1 being within the significant ASoC and 0 otherwise. We used SNPs that did not pass the ASoC testing (FDR>5%) for comparison. For testing disease GWAS enrichment, TORUS assumes that every variant is a risk variant or not, represented by a binary indicator variable (1 or 0). The prior probability of the indicator of a SNP being 1 depends on its annotations. Here, we tested ASoC SNPs categorised by their cell type x time against a selection of diseases/traits GWAS databases. All the annotations are encoded as binary (1 if an SNP is found in the corresponding GWAS database, 0 otherwise). We performed univariate analysis in TORUS to assess the enrichment of each ASoC cell type x time combinations. The GWAS datasets used for enrichment/TORUS analysis were from multiple sources, including NPDs, neuro-related traits, and control disorders/traits, as listed in Table S33.

### Analysis of genetic effect sharing between neuronal activity eQTLs, GTEx eQTL, and ASoC

We used Storey’s *n*1 (Pi1) analysis to analyze the sharing between our eQTLs and other summary statistics. This is a common approach to investigate sharing of eQTLs under different cell types. For each context in our eQTLs and each tissue in GTEx brain eQTLs, we obtained all eQTLs (FDR < 0.05) in a context and estimated the proportion of non-null tests (*π*1) based on the binomial *p* values of these gene-SNP pairs in a GTEx brain tissue. A similar approach was taken when we computed *π*1 for our dynamic eQTLs in GTEx brain tissues.

We also used Storey’s *π*1 analysis to investigate the sharing between our eQTLs and ASoCs. For each ASoC SNP with FDR < 0.05, we located its nearest gene and obtained the nominal *P* values of the corresponding eQTLs in the matched context if possible. Then, we estimated non-null proportions of these *P* values.

### caQTL Mapping

We calculated pseudo-bulk counts from ATAC-seq for GABA neurons of 94 iPSC lines and Glut neurons of 95 iPSC lines (after QC) under each time point. The peak insertion counts were initially normalized for library size using the trimmed mean of M values (TMM) method. Subsequently, we applied a rank-based inverse normal transformation (INT) to standardize the accessibility of each peak to a standard normal distribution, utilizing the RNOmni package (*105*). These counts served as the input for caQTL analysis. We employed TensorQTL (*106*) for caQTL mapping. To adjust for batch effects and confounding factors, we included three genotype principal components and between four to nine chromatin accessibility PCs as covariates in our analysis:

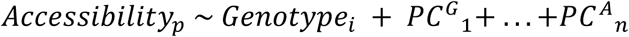

where *Accessibility*_*p*_ denotes the normalized chromatin accessibility of peak *p*, *Genotype*_*i*_ genotype of a *cis*-SNP *i*, *PC*^*G*^_*n*_ are genotype principle components, and *PC*^*A*^_*n*_ are the chromatin accessibility principle components,

For each peak, we conducted 10,000 permutations and computed beta-approximated *P*-values of the top QTLs. To account for multiple testing, we calculated *Q*-values using the qvalue package. Peaks with a *Q*-value ≤ 0.05 were considered significant and designated as cPeaks (caQTL peaks). This threshold controlled the *FDR* and ensured the reliability of our findings.

To investigate time-dependent QTL effects for each cell type, we included the union of cPeaks from all time points. This inclusion ensured that all cPeaks from different time points were considered. For each cPeak, we identified and tested the SNP from the strongest QTLs across all time points. This SNP served as the representative variant for assessing QTL effects. We first log-normalized the counts using the NormalizeData() function in Seurat, then applied rank-based INT to each peak. To test time-dependent effects, we modified the caQTLs test above, including Time, and Genotype-Time interaction term. We included 3 genotype PCs, 5 chromatin accessibility PCs, and their interaction effects with time as covariates and ran linear regression as:

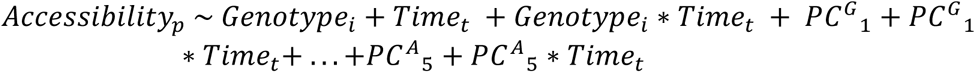

where *Time*_*t*_ is a categorical variable of the duration of KCl stimulation.

We employed an ANOVA test to compare the full model (including the time-genotype interaction effects) with a reduced model (excluding the time-dependent effect). This allowed us to assess the significance of the time-dependent effect on QTLs. FDR correction was applied to adjust for multiple comparisons, ensuring the robustness of our results. To validate the test statistics and avoid inflation, we performed 1,000 permutations. This permutation approach verified that the observed test statistics were not artificially inflated, thereby enhancing the reliability of our time-dependent QTL effect analysis.

### ASoC mapping

All 95 lines (after QC) were used to maximize the statistical power. We applied an improved, two-step ASoC mapping with WASP to calibrate for mapping bias and to gain more accurate results. BAM files generated from Cell Ranger ARC (atac_possorted_bam.bam) were firstly split and re-merged by cell line using barcodes from initial demultiplexing. Each line-specific BAM file included aligned reads of all cell types and three stimulation stages. GATK (4.2.6.0) HaplotypeCaller was applied to each BAM file (per cell line), which identified and extracted all heterozygotic SNP sites and homozygotic SNP sites carrying alternative alleles. The extracted SNP coordination was subsequently used to generate the positional reference of WASP. Next, the original BAM files from Cell Ranger ARC were re-split at the cell line x cell type x stimulation stage (time) levels to generate raw BAM files for WASP calibration. For a set of BAM files from the same line, WASP calibration was performed at the individual BAM files using the line-specific SNP heterozygosity information to re-align reads carrying alternative alleles with bwa and remove any bad read pairs. A second round of GATK HaplotypeCaller was used to identify heterozygous SNP sites and their counts in each calibrated BAM file to generate corresponding VCF files as recommended by the GATK Best Practice (software.broadinstitute.org/Gatk/best-practices/) with VariantRecalibrator (-an DP -an Q.D. -an F.S. -an SOR -an M.Q. -an ReadPosRankSum -mode SNP -tranche 100.0 -tranche 99.5 -tranche 95.0 -tranche 90.0) and reference databases including HapMap v3.3 (priority = 15), 1000G_omni v2.5 (priority = 12), Broad Institute 1000G high confidence SNP list phase 1 (priority = 10), Mills 1000G golden standard INDEL list (priority = 12), and dbSNP v154 (priority = 2). Heterozygous sites with tranche level >95.0% were extracted. And only SNPs with corresponding rs# records found in dbSNP v154 were retained. The individual VCF files were subsequently merged at the cell type x stimulation stage level to increase the statistical power (each merged output included 96-100 lines as cell type varied). After merging, Biallelic SNP sites (GT: 0/1) with minimum read depth count (DP) ≥ 30 and minimum reference or alternative allele count ≥ 2 were retained. The SNPs were further filtered by total read depth of 10 per sample called as heterozygous. The binomial *p*-values (non-hyperbolic) were calculated using the binom.test(x, n, P = 0.5, alternative = “two.sided”, conf.level = 0.95) from the R package, and Benjamini & Hochberg correction was applied to all qualified SNPs as the correcting factor of R function p.adjust(x, method = "fdr"). We set the threshold of ASoC SNP at FDR value = 0.05.

### Homer enrichment analysis of TF-binding motifs for ASoC SNPs

We used HOMER (*107*) to assess the enrichment of TF binding motifs in sequences flanking ASoC SNPs (±25 bp) adapted from our previous investigations (*65*). We used the JASPAR database (2022 release) with all 522 human TF motifs. For each cell type x time, we used the ASoC SNPs (FDR < 0.05) as input intervals. The parameters used were findMotifsGenome.pl <input.bed> hg38 -cpg -mknown jasper2018.known <output>. HOMER provided the genetic background for the enrichment test, and the enrichment significance was derived from HOMER.

### Brain eQTL enrichment analysis for ASoC SNPs

We performed eQTL enrichment analysis to investigate the differences in the association between ASoC and non-ASoC SNPs in GTEx cortex-eQTLs (*55*) and PsychENCODE eQTLs (*56*) databases with adaptation (*65*). For ASoC/non-ASoC SNPs in each set of cell type x time combination, the SNPs were assigned to genes (eQTL targets) if the distance of the SNP and its corresponding gene TSS was less than 500 kb, and the SNP was associated with the expression of the corresponding gene at FDR < 0.05.

### Micro-C analysis of ASoC SNPs and their targets

The Micro-C experiment was conducted in co-cultured Ngn2-Glut and GABA neurons of two donor lines (CD11, CD12) at 0 h, 1 h, and 6 h post-stimulation. Dovetail™ Micro-C (Pan-promoter Capture) kit was used process the samples. To minimize background interference from cell-line discrepancies, we used the samples derived from a single line (CD11) from comparing different time points. The Micro-C data was performed by an external vendor and analysed using ChiCAGO (*108*) at 500 bp resolution. To assign the micro-C targets of each SNP, we firstly associated each SNP (0 h, 1 h, 6 h) with its nearest interacting micro-C fragment (from 0 h, 1 h, 6 h samples) if their distance was less than 2.5 kb (upstream/downstream). If such an association could be established, we extracted the other end of the interacting micro-C fragment and checked if it overlapped with the promoter region (-2 kb / +1 kb of TSS). If the other fragment fell into the promoter region, we added the original SNP to the pool of ASoC SNPs with their corresponding identity.

### Promoter and enhancer enrichment analysis for ASoC SNPs

We defined the interactions of ASoC SNPs and human enhancers/promoters according to our previously developed algorithms with adaptation (*65*). The analysis was similar to that implemented in GREAT and was described previously (*68, 109, 110*) for testing the folds of peak enrichment within the annotated genomic regions and epigenetically annotated regions. To adapt the original algorithm for SNP-based testing, we developed an updated version of the algorithm, and the enrichments were calculated using the formula below:

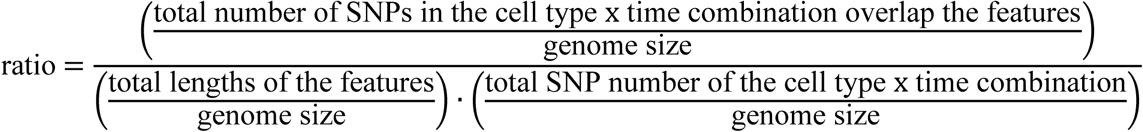

For ASoC SNPs that were cell-type-specific or shared by three cell types, we analyzed enrichment considering SNP intervals as the unit similar to OCR peaks. The enrichment *p*-values were estimated based on the binomial test as implemented in GREAT and as previously described (*109, 110*).

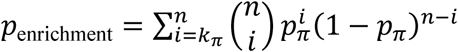

from which we calculated the p-value using the binom.test(x, n, p, alternative=“greater”) function in R, where

*x* = numbers of SNPs that fell within the designated epigenetically annotated features;
*n* = total length of the SNP intervals (in bp);
*p* = total length of the designated epigenetically annotated features (in bp) / genome size (3.2 x 10^9^ bp)

The Gene-based annotation of the genome was derived from GENCODE v35 as part of the built-in database of the updated HOMER package (*107*). The list of human forebrain/non-forebrain enhancers was from the VISTA enhancer browser (*111*). Human-gained enhancers were acquired from GSE63648 using data from either the frontal or occipital cortex at 12 post conception weeks (PCW) and marked by H3K27ac or H3K4Me2 (*112*). The definitions of chromatin state were assembled using an imputed 25-state model derived from individual #E081 of fetal brain tissue by the Roadmap Epigenomics Project (*110, 113*). For estimating the enrichment of ASoC SNPs (Fig. 2A), the categories of epigenomic features used for promoter and enhancer annotations were as used in (*110*).

### Integrative analysis of NPD GWAS with neuronal activity eQTLs or caQTLs

Causal TWAS (cTWAS) (*59*) was used to analyze the heritability of NPD mediated by the molecular traits from our QTLs and ASoCs, and to discover putatively causal genes (eQTL-based cTWAS) and cPeaks (caQTL-based cTWAS) for NPD phenotypes. cTWAS is a generalization of the standard TWAS. In standard TWAS, one first trains the prediction models of gene expression traits from eQTL data, and then correlates the predicted expression traits, one at a time, with the phenotype in a GWAS cohort. However, due to confounding by nearby variants and other expression traits, TWAS has high false positive rates (*59*). In cTWAS, we control the confounding by fitting a joint regression model of the phenotype against all gene expression traits and all variants in the same region. Let *G*_*m*_ be the genotype of the m-th variant, and *X*_*j*_ be the expression of the *j*-th gene. Assuming the prediction models of gene expression are given, we denote *X̃*_*j*_ as the genetically predicted expression of *X*_*j*_. Our model is:

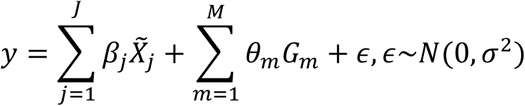

where *β*_j_ and *θ*_m_ are the effect sizes of gene expression j and the variant m, respectively. cTWAS assumes a spike-and-slab prior for the effects of gene expression and variants. The parameters of these distributions are estimated through an expectation-maximization (EM) algorithm, using all the data from the genome. Once these parameters are estimated, cTWAS solves the regression model above with SuSiE (*114*), a fine-mapping method. The results would be Posterior Inclusion Probabilities (PIPs) for all the expression traits and variants. While the model is formulated using individual level data, the method can be used with summary statistics. In that case, a reference LD matrix matching the GWAS samples is used. As a Bayesian method, cTWAS does not need to correct for multiple testing as traditionally done by frequentist methods. Intuitively, the PIPs have already included the prior information: the low prior probability of a molecular trait or variant having a causal effect effectively controls for multiple testing.

The cTWAS model above has only gene expression in a single context. But it is easy to extend the model to allow multiple groups of QTL in the same model, with exactly the same algorithm. For example, we could have eQTLs from multiple tissues or cellular contexts; or one eQTL group and one caQTL group, in the model. We note that different groups generally have different prior parameters, most importantly, the proportion of molecular traits having effects on the phenotype in each group. These parameters are estimated by cTWAS. This allows us to favor certain groups of molecular traits (those with higher prior) in the fine-mapping analysis. We used this version of cTWAS in our study. This integration can combine evidence across multiple groups to improve power. For example, if we have eQTLs across several contexts in the model, then we can obtain the PIP of a gene in each of the contexts. These PIPs can be combined to obtain a single summary of how likely a gene is causal, denoted as “gene PIP”. In our study, the PIP of a gene is simply the sum of PIPs of all expression traits of this gene across all contexts. This gene PIP approximates the probability that the gene is causal in at least one context. Meanwhile, we can also quantify the importance of each context, defined as the ratio of PIP of that context over the total PIP across all contexts. This allows us to assign causal context for a putative causal gene.

When we used cTWAS to integrate eQTLs with NPD traits, we first extract top eQTLs per gene per context and use these eQTLs as the genetic prediction models (denoted as “weights”) of the gene expression traits. cTWAS then includes eQTL data across all 9 contexts in analysis. We note that even though we analyzed all 9 contexts together, a particular gene may occur only in a subset of 9 contexts, as it may not have eQTLs in all contexts. However, one problem is that our eQTL study has incomplete power, thus for a particular gene, we may miss some relevant contexts. To address this issue, we used a relaxed threshold. Suppose a gene has eQTLs in one context (C1), but no eQTL in the second context (C2). If the top eQTLs in C1 has *p* value < 0.1 in C2, then this eQTL would be included as the weight in C2. This allows us to include as many contexts as possible for a single gene.

To run cTWAS, we used the default setting with the prior variance shared by types of QTL, meaning QTLs across all 9 contexts would have the same variance, and 10% down-sampled SNPs used for estimating parameters. We allowed at most 5 causal signals per locus. We used the LD reference computed from 10,000 White British samples from UK Biobank (*59*). In characterizing the high confidence genes of cTWAS, we identified the contexts that support the findings. If the total PIP from the stimulated states (1 h and 6 h) was greater than the PIP from 0 h by at least 0.5, we denoted the gene as “dynamic”; and “static” otherwise.

We used the same procedure when applying cTWAS to caQTLs and ASoC variants. Here the units for analysis are not gene expression traits, but chromatin accessibility of OCRs (that is, cPeaks). We included all the OCRs with either caQTLs or ASoC SNPs. For each OCR, we used either the top caQTLs or the associated ASoC SNP as the weights. In the cases where a peak is associated with both caQTLs and ASoC, we chose the variant with smaller nominal *P* values. In addition, we used *P* < 0.05 to determine what contexts to include in cTWAS; for example, for a SNP that is a caQTL or ASoC in one context (under the FDR cutoff), if the SNP has *P* < 0.05 for the same peak in another context, then the second context would also be included in cTWAS.

### Single-cell DEG analysis in SCZ and TF-KO neurons

We performed DEG analysis between neurons from SCZ donors and those from controls to identify genes that exhibited SCZ case-control differential neuronal activation. Briefly, we first subset the snRNA-seq data of the 28 SCZ lines and the sex/age-matched 28 controls (17 males and 11 females in SCZ cases with an average age of 48; 16 males and 12 females in controls with an average age of 50). The subset Seurat object was first split by individual x cell type and processed by standard Seurat SCTransform for data transformation. Subsequently, all transformed libraries were reintegrated by Seurat IntegrateLayers using “HarmonyIntegration” method to construct the master Seurat object for MAST analysis using the zlm hurdle model. We used covariates including cell co-culture batch, sex, age, and cell type fraction in MAST test. The Seurat wrapper FindMarkers was used to find DE genes between SCZ case and control groups. Only genes expressed in at least 5% of cells in either comparison group were used. We recalculated FDR for the result of each context (cell type/ timepoint). FDR < 0.05 was used as cut-off for DEGs used for the gene set enrichment analyses.

For scRNA-seq of TF-KO neurons, we followed similar data processing and QC procedure as descried above. For DEG analysis of each TF-KO, we subset 630 cells (the minimum number) for each cell type/time per cell line and used MAST in Seurat as described above with cell line as a covariate to control for random effect.

### Statistical analyses

For genetic analyses (eQTL and caQTL mapping and GWAS enrichment analyses), different statistical methods were used and specified in the method section above and in different figure legends. For other assays, unless otherwise specified, Student’s *t*-test (between two groups) or the Kruskal Wallis test with Dunn’s multiple comparisons and *P*-value adjustment (more than two groups) was used to determining significance between groups. Samples were assumed to be unpaired and have non-parametric distribution unless otherwise specified. Data were analyzed using R 4.3.2 and GraphPad Prism 10. Results were considered as significant if *P* < 0.05 (*: *P* < 0.05; **: *P* < 0.01; ***: *P* < 0.001; ****: *P* < 0.0001). All data are reported as mean ± SEM.

## Supporting information

Revised Supplementary Materials and Methods

Revised Supplementary Tables

## Acknowledgments

We thank Rutgers University Cell and DNA Repository (RUCDR; 2U24MH068457) for producing iPSC lines of Molecular Genetics of Schizophrenia (MGS) cohort. We also thank Dr. Matthew Stephens (University of Chicago) for providing feedback on eQTL analyses.

## Author contributions

L.L. performed eQTL and cTWAS analyses. S.Z. led the sn-Multiomics data QC, DEG, and DA peak analyses, and ASoC analysis. Z.W. performed caQTL, transcriptome clustering, OCR-gene linkage, and GRN analyses. H.Z. carried out neuronal differentiation and stimulation as well as sequencing library preparation. C.L. performed the multiomics data QC, DEG, and DA peak analyses. C.T. and E. O. analyzed scRNA-seq data for TF knockout experiment and helped with other data analyses. A.C.D. helped with data QC, DEG (SCZ case-control), and DA peak analyses. D.S., A.B., and A.M. generated and characterized the iPSC lines with TF knockout. X.S. and X.Z. helped with NPD risk gene enrichment in gene clusters and with SCZ case-control comparison. A.K., B.J., and W.W. helped with iPSC culture and CRISPR/Cas9 editing. Z.P.P. and A.R.S. helped with experimental design and data interpretation. L.L., S.Z., Z.W., H.Z., X.H., and J.D. wrote the manuscript with input from other authors. X.H. and J.D conceived, designed and supervised the lab work and data analyses.

## Data and materials availability

This study did not generate new unique reagents. The NIMH repository iPSC lines used in this study, including the ones derived from CRISPR/Cas9 editing, will be distributed from Coriell Institute for Medical Research (CIMR, www.coriell.org) under a material transfer agreement (MTA) with NIMH Repository and Genomic Resource (NRGR) (www.nimhgenetics.org). The corresponding author is available for providing guidance on obtaining these iPSC lines. The snRNA/ATAC-seq raw data and processed count matrices are accessible at Gene Expression Omnibus (GEO) under accession codes PRJNA194194 and GSE286488. The scRNA-seq raw data and the processed data for TF-KO are accessible at GEO under accession number GSE312445. Genotypes for iPSC donors can be obtained through controlled access at dbGaP (phs000021.v3.p1 and phs000021.v3.p2 for MGS lines and phs002032.v1.p1 for CIRM lines). All code used in the analyses is available on Zenodo (*115*).

## Supplementary Materials

Supplementary text Figs. S1 to S26 Tables S1 to S33 References (116–125)

