## Supplementary material for "Single-cell multiomics of neuron activation reveals context-specific genetics of brain disorders": Revised Supplementary Materials and Methods

**The PDF file includes:**

Supplementary Text

Figs. S1 to S26

Tables S1 to S33

Reference (116-124)

Fig. S20. Strong correlation of the effect size between our neuron activity caQTL and other caQTLs. 46
Fig. S21. caQTL (including ASoC) mapping. 48
Fig. S22. Functional validation of ASoC SNPs by comparing to brain eQTL, reporter gene assay, and Micro-C. 49
Fig. S23. Enrichment of ASoC SNPs in different regulatory sequence elements. 52
Fig. S24. Integrative analysis of caQTL and GWAS of NPD phenotypes. 53
Fig. S25. Differential expressed genes (DEGs) in neurons from SCZ cases. 56
Fig. S26. Consistent DEG results from using pseudobulk or single-cell data. 57
***List of Supplemental Tables*** 58
***References and Notes (only appeared in Supplementary Materials)*** 60

### Supplementary Text

#### Sn-Multiomics data quality control (QC) and preliminary analysis

The data were from a total of 84 single nucleus (sn)-Multiomics (snRNA-seq and snATAC_seq) sequencing libraries, each containing cells from a single cell co-culture batch (often with cells from multiple induced pluripotent stem cell (iPSC) lines) at each time point. These libraries were sequenced in 6 different batches. The sequencing reads of 1,053,422 nuclei (Tables S1-2) were aligned to a hybrid human/rat genome using 10x Genomics Cell Ranger ARC (v2.0.2) and analyzed in Seurat 5.1.0 (*85*). We performed data QC for each sequencing batch and after merging all batches (Fig. S1A). For each sequencing batch, we demultiplexed (*84*) different donor lines ($n$=2-5) in each sequencing library, and confirmed the clear separation of clusters of iGlut (*SLC17A6*+) and iGABA (*GAD1*+/*GAD2*+) on Uniform Manifold Approximation and Projection (UMAP) (Fig. S1B-G). For snATAC-seq data, we transferred the cell identity labels from snRNA-seq data on UMAP of gene-activity-score (gact) (Fig. S1H-I) and verified the robust transcription start site (TSS) read enrichment (>5) (Fig. S2A-D, Table S2). After merging all the post-batch-QC data, we obtained 651,012 neurons with an average of 7,510 unique molecular identifiers (UMI) per nucleus that are comparable across the 100 lines (only 95 lines were used for downstream ATAC-seq data analysis due to the substandard TSS enrichment metric of 5 lines in sequencing batch 22) (Table S3, Fig. S4A-D). We defined three major subtypes of neurons: GABA ($n$=251,501), NEFM+ Glut (npglut; with stronger *NEFM* expression) ($n$=159,735), and NEFM- Glut (nmglut; with weaker *NEFM* expression) ($n$=137,564) (Fig. 1E-H, Fig. S4E-G, Table S1), with reproducible cell type clustering patterns across samples (Fig. S3A-E).

#### Transcriptomic and epigenomic landscape of cell-type-specific neuronal activation

With sn-Multiomics data of co-cultured human Glut/GABA neurons and mouse glia, we first identified differentially expressed genes (DEGs) upon stimulation by KCl (1 h vs. 0 h, and 6 h vs. 0 h) in each neuron subtype. We used pseudobulk RNA-seq data from the 18 donor lines (sequencing batch 24; Table S1) in our limma DEG analysis to minimize any possible batch effect. We performed variation partition analysis to evaluate relative contribution of each variable to the expression variation (Fig. S7A). Although our main DEG result was based on the 18 lines, we made sure that the main cell line characteristics and sequencing QC were similar between the 18 lines and the full cohort of 100 lines (except for more snATAC-seq read pairs per cell in the full datasets) (Table S1-3). Moreover, we confirmed that the log2FC and the list of DEGs upon stimulation were highly similar between the 18 lines and the full cohort (Fig. S7B,C; Table S4). After Combat-seq correction for co-culture batch, the snRNA-seq samples of the 18 lines showed clear separation by time points in principal component analysis (PCA) plots (Fig. S7D). We used log2 fold change (FC) > 0.25 or < -0.25 and *FDR* < 0.05 as a cutoff for DEG. The upregulated genes showed much larger magnitude of expression changes than the downregulated genes, with the well-known ERGs such as *FOS* and *FOSB* exhibiting the largest FC (Fig. S7E, Table S4). Comparing to a similar study using iPSC-derived Glut and GABA neurons separately cultured and assayed by bulk RNA-seq (*29*) (Fig. S8), we found that for the upregulated genes at 45 min of KCl stimulation (*29*), 79% of them were also upregulated with $\log_2$FC > 0.5 at 1 h in our npglut (60.2-fold enrichment, Fisher's exact test $P = 2.4 \times 10^{-52}$) and GABA (24.5-fold enrichment, Fisher's exact test $P = 2.3 \times 10^{-67}$), while about 81-83% were upregulated ($\log_2$FC > 0.25) in our npglut

(24.5-fold enrichment, Fisher's exact test $P$ = 1.2 × $10^{-33}$) and GABA (19.7-fold enrichment, Fisher's exact test $P$ = 1.8 × $10^{-46}$). For late response genes (LRGs), about 52-62% of those upregulated at 4 h in Sanchez-Priego et al. study (*29*) were also upregulated ($log_2FC>0.5$) at 6 h in our npglut (7.2-fold enrichment, Fisher's exact test $P$ = 2.3 × $10^{-175}$) and GABA (9.9-fold enrichment, Fisher's exact test $P$ = 4.9 × $10^{-248}$), while about 69-71% were upregulated with $log_2FC>0.25$ in our npglut (6.1-fold enrichment, Fisher's exact test $P$ = 6.6 × $10^{-162}$) and GABA (8.1-fold enrichment, Fisher's exact test $P$ = 3.0 × $10^{-212}$). For genes downregulated at 4 h (neurons at 1 h had too few downregulated genes) of KCl stimulation (*29*), 45~74% of them were also downregulated (with $log_2FC<-0.5$ or -0.25) at 6 h in our npglut or GABA (4.3 to 4.9-fold enrichments; Fisher's exact test $P$ = 1.7 × $10^{-106}$ to 3.6 × $10^{-133}$). The larger number of DEGs in our study may be attributed to the larger sample size and single-cell resolution.

We also compared our results to that from studying mouse models of neuron activation (*21, 37*). In mouse dentate granule neurons upon *in vivo* electroconvulsive stimulation (*21*), 3,006 genes were upregulated and 2,115 were downregulated at 1 h after stimulation, while 3,087 were upregulated and 2,845 were downregulated at 4 h after stimulation ($log_2FC > 0.25$ or < -0.25, q-value <0.05) (*21*). The numbers of in vivo DEGs are comparable to that in our stimulated Glut neurons. More than 70% of the DEGs in both datasets had same directional expression changes (mostly upregulated) and showed strong correlation (Pearson's $R^2$ =0.272) of $log_2FC$ (Fig. S8B). For the 2,471 shared LRGs, over 64% of them had same directional expression changes, albeit a weaker but significant Pearson's correlation of $log_2FC$ ($R^2$=0.0597) (Fig. S8C). In another mouse study of sensory-experience-activated neuronal expression (*37*), of the 8,313 DEGs in at least one cell type, 419 upregulated and 192 downregulated genes (> 2-fold) were defined as ERGs ($n$=362) and LRGs ($n$=249). 59.2% of the Ex ERGs (29/49; 15.6-fold enrichment, Fisher's exact test $P$ =6.1 × $10^{-20}$) and 29.7% of the Ex LRGs (11/37; 4.4-fold enrichment, Fisher's exact test $P$ = 8.7 × $10^{-6}$) were shared with our npglut or nmglut DEGs (> 2-fold) at 1 h or 6 h (Fig. S8D-E). At a $log_2FC$ cutoff of 0.25 (a cut-off that allows inclusion of more NPD risk genes), our npglut or nmglut DEGs also showed similar enrichments (3.5 to 6.6-fold) for mouse sensory-experience-induced ERGs and LRGs, with a notable increase of overlapping mouse sensory-experience-induced Ex ERGs (55.1%) and LRGs (54.1%) (Fig. S8D,E). For our iGABA neurons, DEGs with >2-fold or $log_2FC>0.25$ expression increase at 1 h or 6 h also showed significant overlap with mouse sensory-experience-induced Inh ERGs (34.8% or 16/46; 7.2-fold enrichment, Fisher's exact test $P$ = 4.0 × $10^{-8}$; FC > 2-fold) or LRGs (27.5% or 11/40; 5.1-fold enrichment, Fisher's exact test $P$ = 6.4 × $10^{-5}$; FC > 2-fold) (Fig. S8D,E). At an individual gene level, we found that some sensory-experience-induced canonical immediate-early genes like *Nr4a1, Nr4a2, Nr4a3, Fos, Fosl2*, and *Egr1* known to regulate the late phases of gene expression (*37*) were also shared in our ERGs. We also confirmed that LRGs tend to be more cell type-specific in both datasets; for example, *Crh*, an Inh neuron LRG in mice (*37*) that encodes the stress hormone, corticotropin-releasing hormone (*37*), was increased (> 2-fold) by KCl stimulation only in GABA at 6 h.

To examine the biological relevance of these activity-dependent DEGs, we performed gene set enrichment analyses. Analysis of synaptic gene ontologies (SynGO) (*116*) showed that only the upregulated genes showed significant enrichment for synaptic genes (Fig. S9A). Our MAGMA (*117*) analysis found that GWAS risk of NPD or traits showed higher enrichment in upregulated genes than downregulated or unchanged genes, with strongest enrichment for SCZ (Fig. S10A). We also found a lack of enrichment for autism spectrum disorders (ASD), which may be due to the relatively small number of ASD GWAS risk loci. We thus performed an enrichment analysis using a set of 102 ASD risk genes from rare variant analysis (*36*), together with a set of SCZ rare variant-based risk genes (*35*), and other sets of GWAS risk genes for SCZ (*7*), bipolar disorder (BP) (*118*), major depressive disorder (MDD) (*4*), and post-traumatic stress disorder (PTSD) (*119*) (Table S5). We found strong enrichment for ASD risk genes among the upregulated genes across cell types, and to a lesser extent for SCZ rare variant-based risk genes (Fig. S9B). Interestingly, the majority of SCZ rare risk genes were upregulated by stimulation, rather than downregulated (Fig. 2B, Fig. S9C). We also confirmed the pseudobulk-based expression changes of some selected SCZ and ASD genes in single neurons (Fig. S9D) and validated the downregulation of *SNAP91* (a SCZ GWAS risk gene) and *IMMP2L* (a GWAS risk gene for SCZ and MDD) by qPCR in independent cell cultures (Fig. S9E).

We next analyzed the snATAC-seq (Fig. 1F, Fig. S1H) to characterize the landscape of activity-dependent open chromatin region (OCR) peaks upon stimulation. For the same 18 donor lines used for DEG analysis, we used limma (*89*) to analyze a set of merged peaks (170K for GABA, 196K for nmglut, and 207K for npglut) to identify differentially accessible (DA) peaks upon neuronal stimulation in each cell type. Peak variation partition analysis showed a similar pattern to expression variation partition, with cell type and time point contributing the most (Fig. S10D). No obvious associated *P*-value inflation was found for DA peak analysis (Fig. S10E). We found that the proportion of DA peaks with FC > 2-fold (~11.8% of OCRs) in KCl-stimulated npglut at 6 h were comparable to that in mouse dentate granule neurons (11.5% of total OCRs, FC>2) upon *in vivo* electroconvulsive stimulation (*21*). We also confirmed that chromatin accessibility of well-established ERGs (e.g., *ARC*, *FOS*, and *NPAS4*) and LRGs (e.g., *BDNF*) (Fig. S10F) were similarly increased in both datasets (*21*). Similar to DEGs (Fig. S7E), the upregulated peaks often showed a larger magnitude of changes (Fig. S10F). As expected, *FOS* showed the largest increase of peak accessibility at 1 h, highlighting its driving role as an ERG in inducing activity-dependent gene expression (Fig. S10F).

**Activity-dependent expression of BDNF is regulated by cell-type-specific OCR**

*BDNF* is a SCZ risk gene (*7*) that encodes a neurotrophin important for neuronal differentiation and synaptic plasticity (*120, 121*). Despite being a well-established LRG, its cell-type-specific regulation is unknown. We found that *BDNF* exhibited late response in both Glut and GABA cells (Fig. 1I-J), but the DA peaks upon stimulation were different between cell types (Fig. S11B, Fig. S12A, and Table S6). The DA peak showing the strongest increase of accessibility in npglut and nmglut was about 49 kb upstream (putative enhancer) of the TSS of *BDNF* (Fig. S11B, Fig. S12A), but with much weaker accessibility in nmglut. However, the DA peak showing the largest increase of peak accessibility in GABA was near the 3'-UTR of *BDNF*. These DA peaks showed peak-gene promoter linkage at 1 h and/or 6 h of stimulation and correlated with *BDNF* expression in their respective cell types (Fig. S11B, Fig. S12A), suggesting their possible enhancer function.

We next validated whether the DA peak 49 kb upstream of the TSS of *BDNF* indeed regulated the activity dependent *BDNF* expression in iGlut. We noted that this DA peak encompassed the binding site of early response AP-1 TFs (FOSB, JUNB, JUND, FOSL1, FOSL2) (Fig. S12B), suggesting an enhancer role of the OCR peak in regulating *BDNF* expression. We thus CRISPR/Cas9-engineered two iPSC lines (CD07 and CD15) by deleting the OCR peak (~600bp)

(Fig. S12C-E). We then differentiated the two isogenic pairs of CRISPR/Cas9-edited iPSC lines into iGlut and carried out KCl stimulation. We found that while *BDNF* expression was similar between the unedited and deleted lines at 1 h, its expression was significantly lower at 6 h of stimulation in the OCR-deleted lines (Fig. S12E). This supports the role of this OCR in regulating the activity-dependent expression of *BDNF* in iGlut. These results highlight that similar transcriptional responses across cell types may have distinct epigenomic regulatory mechanisms.

**Chromatin priming during neuron activation**

We noted frequent chromatin "priming" during neuron stimulation by KCl. Multiple ERGs like *EGR1* and LRGs including *DUSP4, ATP1B1*, *CREM, BDNF, PCSK1,* and *SCG2* (Fig. S6B,C) showed high gene activity scores at 0 h or 1 h, but gene expression was activated later. To mechanistically characterize the observed chromatin "priming", i.e., chromatin was partially active ("primed") before stimulation, we used a published histone mark dataset from iPSC-derived neurons in both stimulated and unstimulated states (*29*). This dataset, however, includes only H3K27ac, but not H3K4me1. We thus used H3K27ac to annotate a set of OCRs that respond to neuron activation and examined the chromatin states of these OCRs before and after stimulation in GABA cells. Specifically, we defined a set of "response cis-regulatory-elements (CREs)" as the OCRs showing differential accessibility upon stimulation ($FDR \leq 0.05$ and $\log_2 FC > 1$) and also having H3K27ac mark in the stimulated state. To better detect priming, we further limited our analysis to CREs whose nearby genes had low expression at 0 h (using the expression of *FOS*, an early response gene, as the threshold). In total, we had 2,776 CREs. We then assessed the chromatin states (open chromatin and H3K27ac) of these response CREs at 0 h (before stimulation). We found heterogeneous patterns of CRE activation (Fig. S14A): (1) Some CREs (33%) were in the closed chromatin state before stimulation, and were activated "*de novo*" by the neuron activation signal; (2) Some CREs (28%) were in the "primed" state: chromatin was partially accessible but had no H3K27ac mark, and became fully activated by neuron activation (see the example below in Fig. R19B,C); (3) Some CREs (39%) already had enhancer activities before stimulation, marked by H3K27ac.

Next, we examined the chromatin states of a few loci before and after stimulation. As an example, in GABA cells (Fig. S14B), we examined the regulatory sequences near an ERG, *cysteine and serine rich nuclear protein 1 (CSRNP1)*, a gene essential for cephalic NPC proliferation and survival in zebrafish (*122*). We confirmed that *CSRNP1* showed very low expression at 0 h (below the level of *FOS*) and much higher expression at 1 h and 6 h of stimulation. The chromatin of the OCR sequence (Chr3:39152481-39153281) near the TSS was partially open (note the high read count, called as a peak by ArchR) at 0 h but became fully active at 1 h. To corroborate the observed chromatin priming at the *CSRNP1* locus, we used seq2PRINT to map TFBS (*123*) in this region. To ensure comparable coverage and library size across cell types and stimulation time points, cells were down-sampled so that each context had similar coverage. We first trained the seq2PRINT model using pseudobulk aggregations of all contexts and all cells. Chromosomes were partitioned into training, validation, and test sets, and the model was fit using a five-fold scheme to ensure evaluation across different chromosomes. Next, the seq2PRINT model was fine-tuned for each context using Low-Rank Adaptation (LoRA). Finally, we calculated TF-binding scores for all significant candidate regulatory elements identified by the co-activation-based OCR–gene linkage analysis, using the average scores across all folds from the LoRA-fine-tuned models. We found weak binding sites at 0 h, however, two strong binding sites of the AP-1 family motif emerged at 1 h. This analysis suggests that the partially open (i.e., primed) chromatin at 0 h likely enabled the access by AP-1 family (FOS/JUNB), which then fully activated the CRE at 1 h.

**Integrative multiomic analysis identifies regulatory program of neuronal activation**

While some TFs, e.g., FOS and JUNB, are well known regulators of early response, the regulatory programs controlling cell-type-specific late response are far less clear. We thus leveraged our multiomic data to identify putative TF regulators. We limited the analysis to TFs that showed differential expression and motif enrichment in at least one condition. We used chromVAR (*124*) to assess a single-cell level motif enrichment score in each of the 9 contexts (3 cell types, 3 time points). Combining motif scores with gene expression, we classified the role of each TF in early or late response. For a given cell type, we considered a TF to be a candidate regulator of early or late response when it showed higher motif enrichment and gene expression at 1 h (vs. 0 h), or at 6 h (vs. 1 h), respectively (Table S13).

We identified 145 candidate TF regulators of early response, nearly half of which are shared by all cell types (Fig. S14C). Some shared TFs with the largest motif changes from 0 h to 1 h are well-known early response TFs, for example FOS, JUNB, EGR1, and NPAS4, while some others have less established roles in early response like BACH2, JDP2, MAFK, and SMARCC1 (Fig. S14D). Compared to early response TF regulators, late response TFs are much more cell-type-specific (Fig. S14E). Only 6 TFs were shared across cell types, including some important for neurodevelopment, such as MEF2C and ESR1 (Fig. S14D, F). This highlighted the distinct regulatory programs controlling late responses in Glut and GABA neurons.

To understand the cell-type-specific responses, we focused on TFs whose roles are limited to only Glut or GABA cells. For Glut cells, we observed a relatively large number of candidate TFs regulating late response. The TFs showing strongest motif enrichment at 6 h are regulators of early response in all cell types, such as the AP1 family TFs (FOSL, JUND) and a member of SWI/SNF family (SMARCC1) (Fig. S14D). While the motifs of these TFs are also enriched in GABA cells, their motif enrichment is much stronger at 6 h in Glut cells, and their expression is also upregulated in Glut cells (Fig. S14D). These results thus suggested that a subset of early response TF regulators continue to drive late response, primarily in Glut cells.

For GABA cells, we found 21 TF candidates for late response (Table S13). Most of the TFs showed modest changes of motif activities between 1 h and 6 h, with a few exceptions (NR2F2, NR2C1, ESRRA) (Fig. S14D). We hypothesized that some TFs without large motif changes from 1 h to 6 h may still play a role in neuronal response. We thus assessed motif enrichment of TFs at 6 h in GABA cells. The TFs showing highest motif enrichment include several GABA-specific TFs, ID3, EVX1, DLX5, and TCF4, a risk gene and a master regulator in SCZ (*49*). These TFs showed high expression and strong motif enrichment even before stimulation (Fig. S14D-F). We also found other TFs likely regulating early response specifically in GABA cells, such as EMX2 and NR4A2 (Fig. S14D-F). In contrast, there were much fewer Glut-specific early response TFs (Fig. S14C). These results thus suggest that distinct TF activities before and during the early response to the stimulation primed the epigenome of GABA cells, leading to varied late responses in GABA cells.

Altogether, our results highlighted a shared regulatory program controlling early responses of all cell types, and distinct programs controlling late responses in Glut and in GABA cells. Some of the key TFs in these programs are NPD risk genes (e.g., *MEF2C*, *TCF4*), highlighting the importance of these regulatory programs to the development of NPD.

### Supplemental Figures

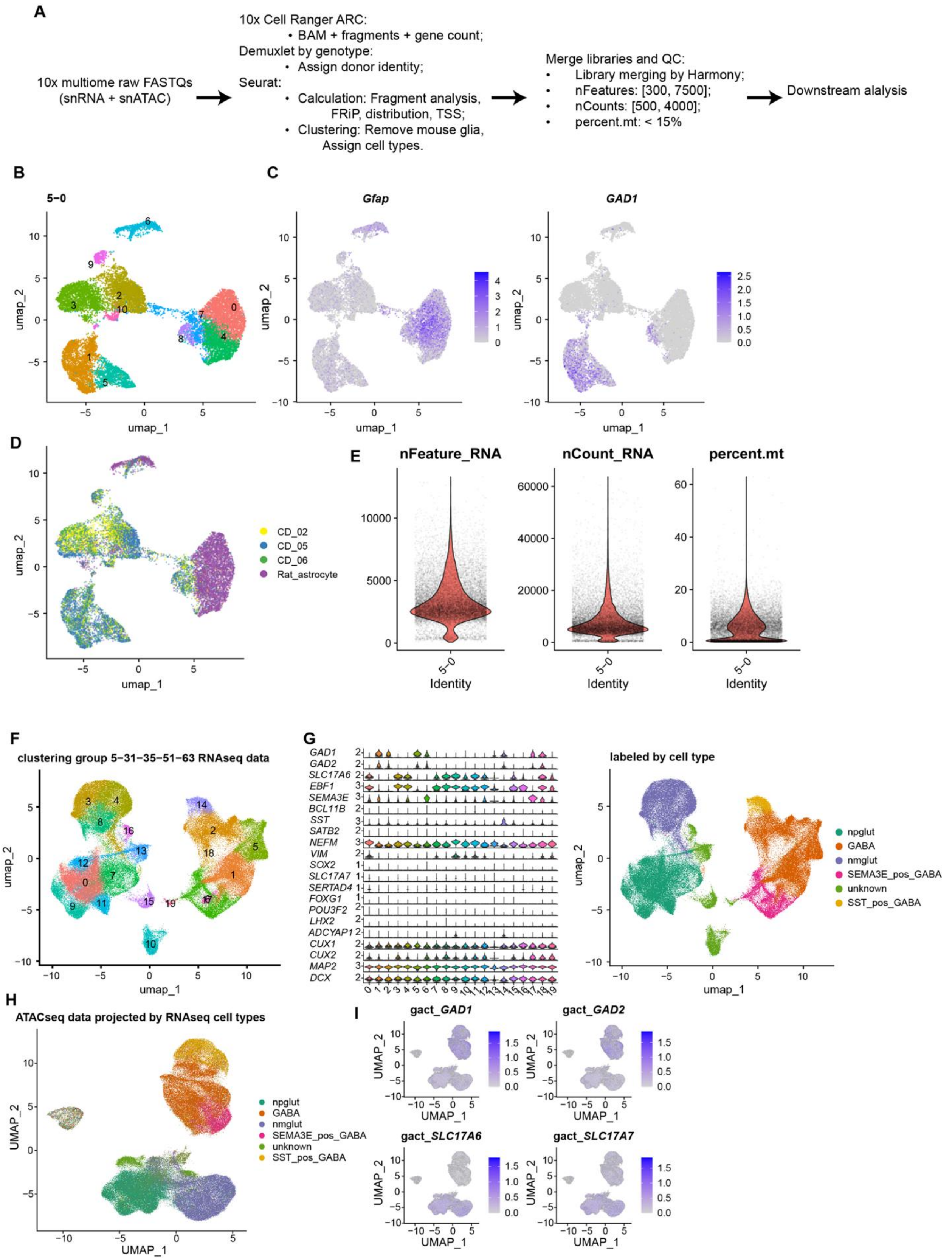

**Fig. S1. Multiomics data processing and quality control (QC) by batch.**

(A) The flowchart depicts raw data processing from FASTQ format to QC-data for downstream analysis. (B) snRNA-seq UMAP shows a representative sequencing library, library 5-0 at 0 h, for cell clustering. (C) Feature plots of the same library show the expression of Rat *Gfap* (astrocyte marker) and human *GAD1* (GABAergic neuron marker). (D) UMAP of the same library shows individual identity of each cell (i.e., demultiplexed three iPSC lines and rat astrocytes). (E) Violin plots of the same library show the distribution of RNA features per cell (nFeature_RNA), UMI counts per cell (nCount_RNA), and the percentage of mitochondrial transcripts per cell (percent.mt). (F) UMAP projection of aggregated snRNA-seq results of sequencing batch 024, including 15 libraries from 18 lines and all three time points (0 h, 1 h, and 6 h). Rat astrocytes have been removed before library aggregation. (G) The violin plot (left) shows the expression of key genes in each cluster of (F) and the UMAP projection (right) shows the assigned major cell type identities. Npglut, iGlut with higher NEFM expression; nmglut, iGlut with lower NEFM expression; GABA, iGABA neurons stained positives for GAD1 and GAD2; unknown, cells cannot be assigned to either iGlut or iGABA. (H) UMAP of aggregated snATAC-seq data from the same sequencing batch (024) with transferred label from snRNA-seq. (I) Gene activity (OCR peak reads) plots (1 kb TSS+gene body) of human GAD1, GAD2 (GABAergic neuron markers) and human SLC17A6, SLC17A7 (glutamatergic neuron markers) of the same ATAC-seq UMAP projection in (H).

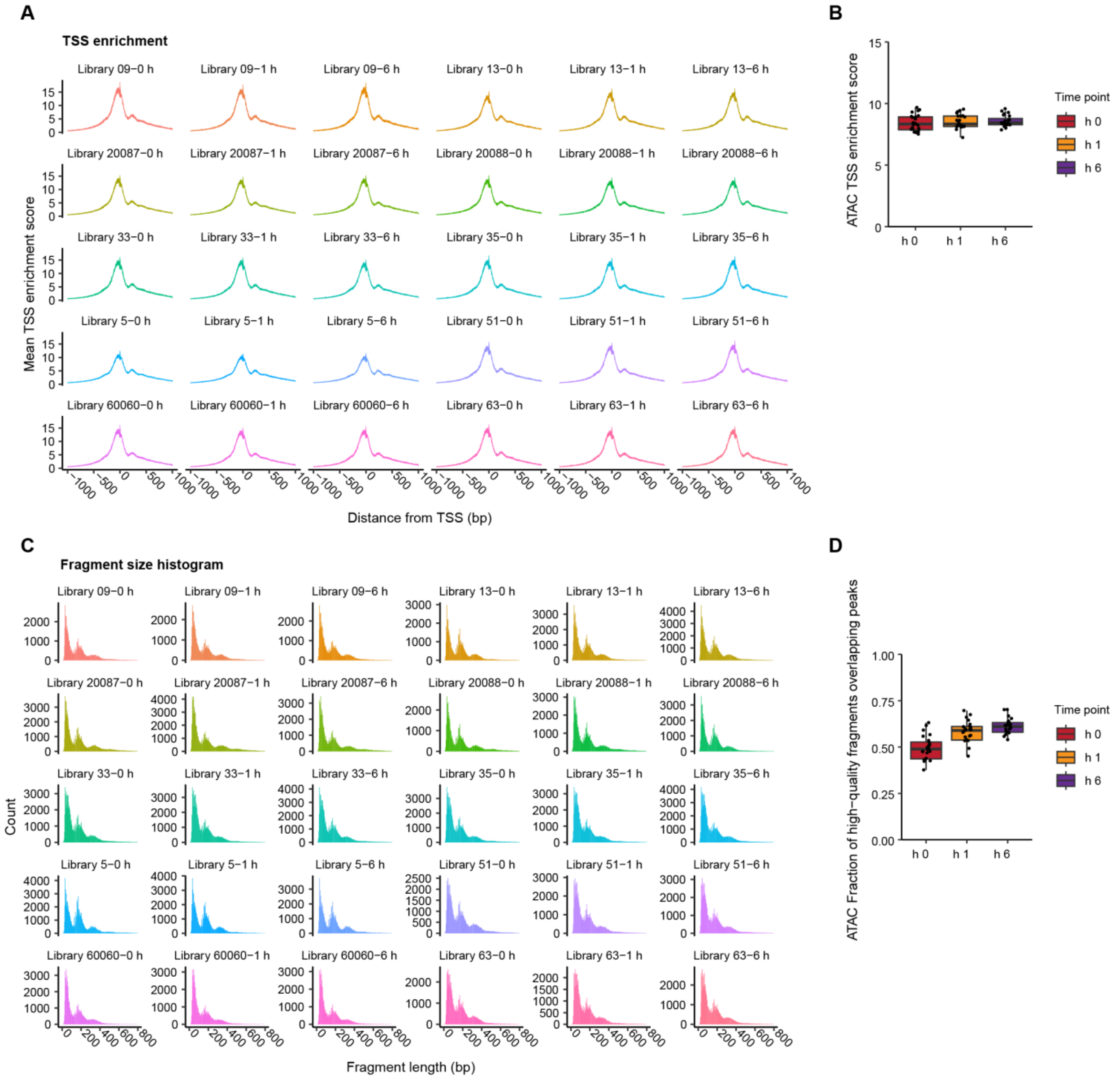

**Fig. S2. QC metrics of snATAC-seq.**

(A) TSS enrichment plots (1 kb TSS+gene body) for 30 representative sequencing libraries, related to Figure 1. (B) The box-whisker plot shows the summarized distribution of TSS enrichment values of all 84 sequencing libraries in the study. (C) Fragment size distribution patterns of the 30 representative sequencing libraries as in (A). Note the expected pattern of nucleosome periodical size in each graph. (D) The box-whisker plot shows the summarized proportions of ATAC-seq Fragment of Reads in Peaks (FRiP) for all 84 sequencing libraries in the study.

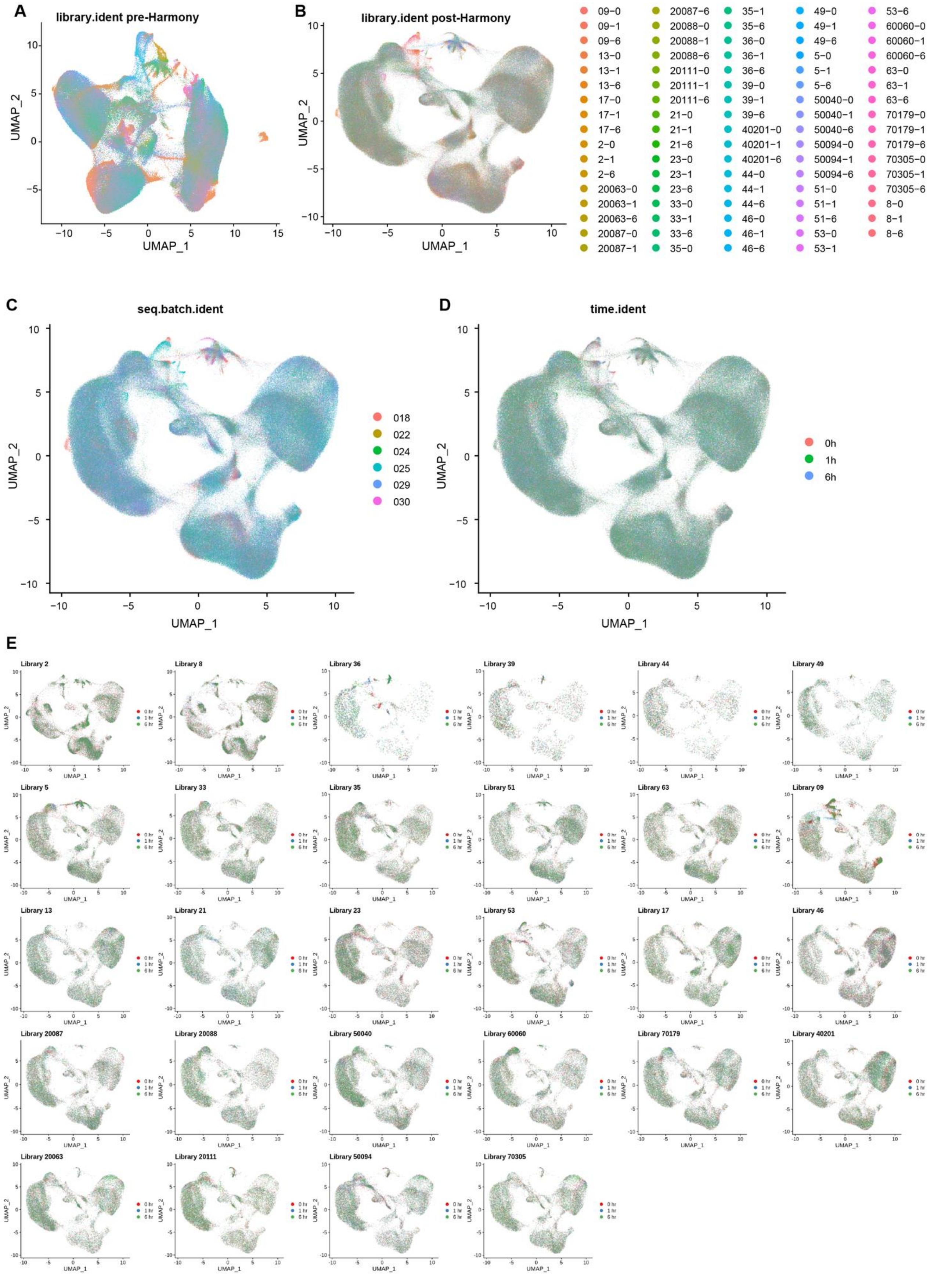
A
library.ident pre-Harmony
B
library.ident post-Harmony
C
seq.batch.ident
018
022
024
025
029
030
D
time.ident
0h
1h
6h
E
UMAP_1
UMAP_2

**Fig. S3. Integrative analyses of snRNA-seq for all 100 lines.**

(A) UMAP projection plot shows the merged raw snRNA-seq data from all 84 sequencing libraries, pre-Harmony normalization. (B) UMAP of the same merged data post-Harmony. (C) UMAP projection of post-Harmony snRNA-seq data stratified by different sequencing batches ($n$ = 6). (D) UMAP projection of post-Harmony snRNA-seq data stratified by different stimulation time points (0 h, 1 h, and 6 h). (E) UMAP of sequencing libraries from (D) show reproducibility of cellular composition and distribution of different contexts (cell types × time points). Note that each UMAP shows three sequencing libraries from the three time points.

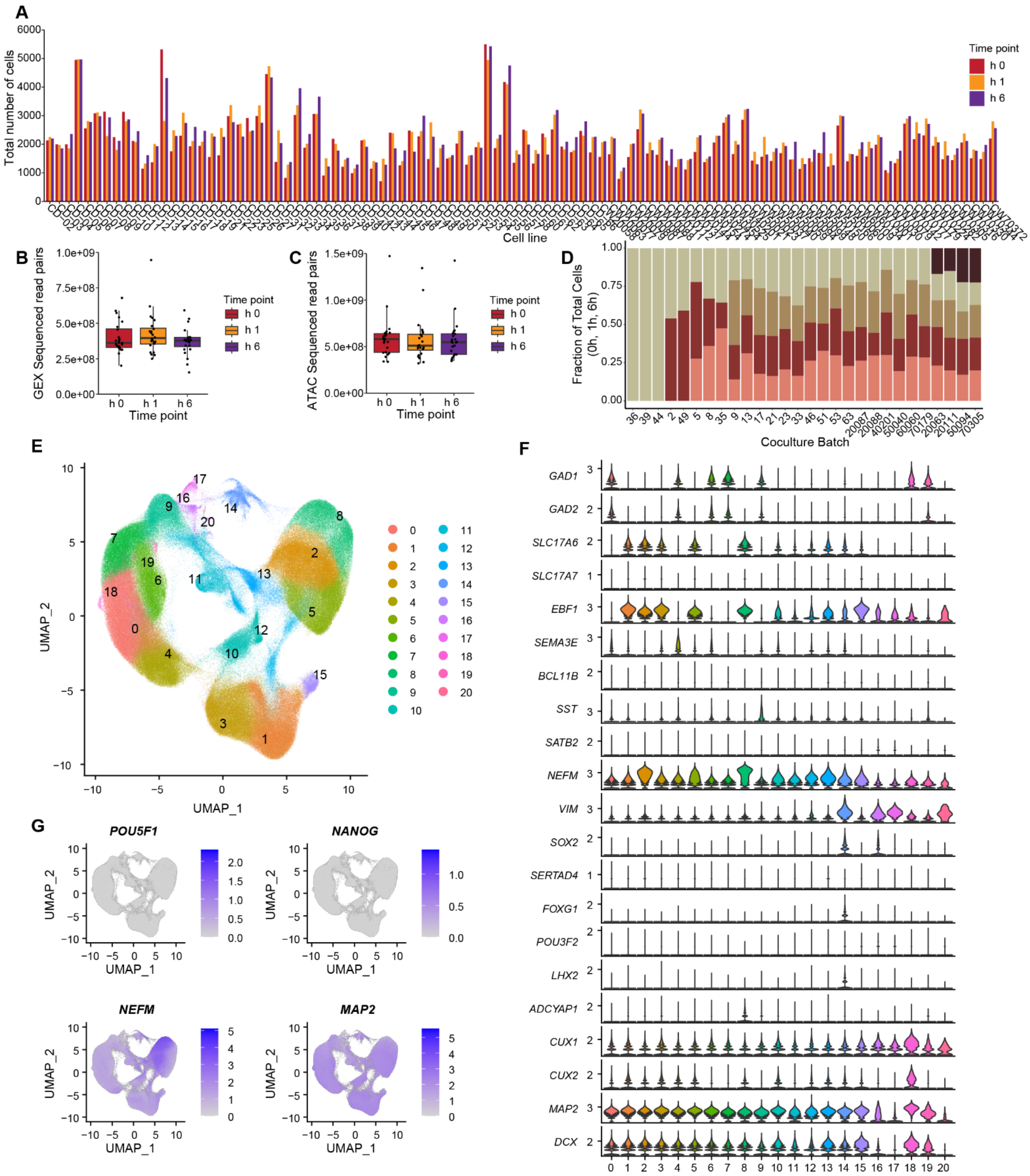
A
Total number of cells
Cell line
Time point
h 0
h 1
h 6
B
GEX Sequenced read pairs
Time point
C
ATAC Sequenced read pairs
Time point
D
Fraction of Total Cells
(0h, 1h, 6h)
Coculture Batch
E
UMAP_1
UMAP_2
F
GAD1
GAD2
SLC17A6
SLC17A7
EBF1
SEMA3E
BCL11B
SST
SATB2
NEFM
VIM
SOX2
SERTAD4
FOXG1
POU3F2
LHX2
ADCYAP1
CUX1
CUX2
MAP2
DCX
G
POU5F1
NANOG
NEFM
MAP2

**Fig. S4. Additional QC metrics on the merged snRNA-seq library.**

(A) Bar graph showing the number of demultiplexed cells of each of the 100 iPSC lines across three time points. (B) Box-whisker plot showing the distribution of mean snRNA-seq UMI counts/cell of all cell lines across three time points. (C) Box-whisker plot showing the distribution of mean snATAC-seq reads/per cell of all cell lines across three time points. (D) Proportion of neurons of each cell line in different pool/co-culture batches. Note that the number of cell lines in each pool are all as expected without any dropout. Each color in a bar represents a different cell line. (E) Leiden-based clustering results of the merged snRNA-seq library (resolution = 1). A total of 20 clusters were defined. (F) Violin plots show the expression of key cell type-specific marker genes across all clusters. (G) Feature plots show the absence of expression of pluripotency marker genes *POU5F1* and *NANOG*, and strong expression of neuron-specific marker genes *NEFM* and *MAP2* in the merged library.

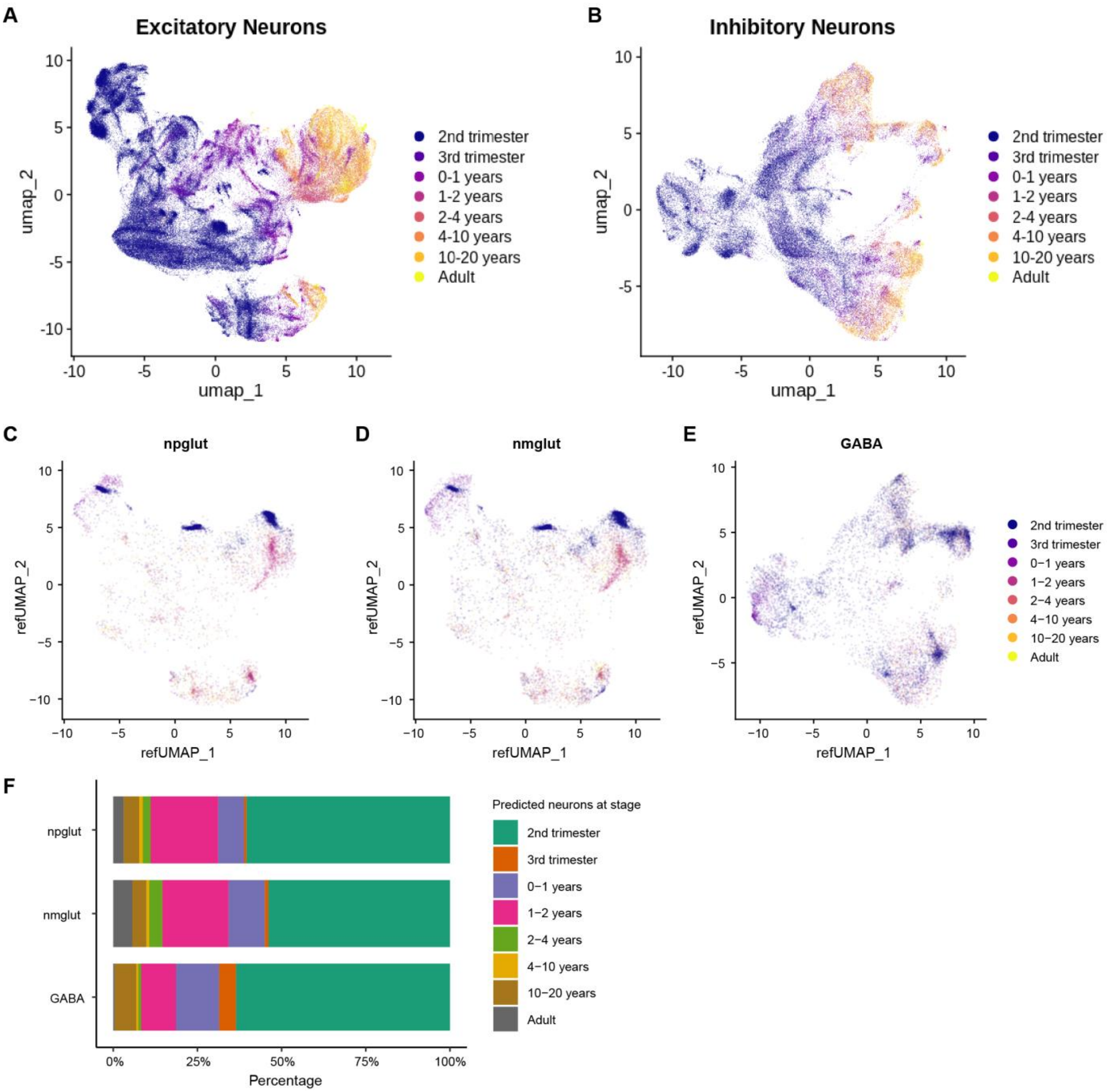

**Fig. S5. Projection of iPSC-derived iGlut and iGABA neurons to brain excitatory and inhibitory neurons of various developmental stages.**

(A) and (B) UMAP projection of all developing brain excitatory neurons and inhibitory neurons (*34*), respectively. (C) and (D) UMAP projection of subsampled 10,000 npglut and nmglut neurons (this study), respectively, using principal components (PCs) derived from A. (E) UMAP projection of subsampled 10,000 iGABA neurons (this study) using PCs derived from B. (F) Percentages of the neurons transcriptionally similar to those matched excitatory or inhibitory neurons at different brain developmental stages.

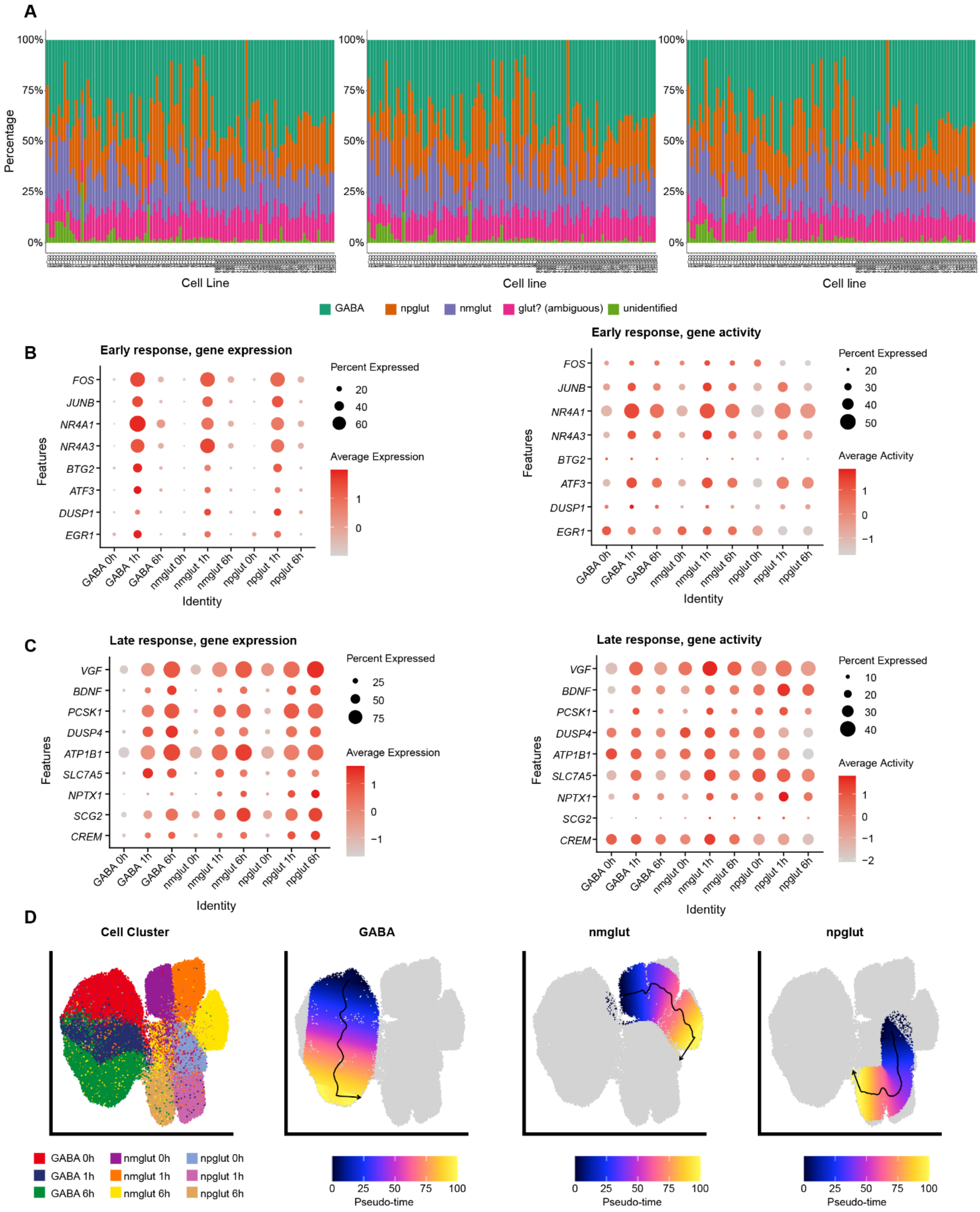
A
Percentage
100%
75%
50%
25%
0%
Cell Line
Cell line
GABA
npglut
nmglut
glut? (ambiguous)
unidentified
B
Early response, gene expression
Features
FOS
JUNB
NR4A1
NR4A3
BTG2
ATF3
DUSP1
EGR1
Percent Expressed
20
40
60
Average Expression
1
0
GABA 0h
GABA 1h
GABA 6h
nmglut 0h
nmglut 1h
nmglut 6h
npglut 0h
npglut 1h
npglut 6h
Identity
Early response, gene activity
Percent Expressed
20
30
40
50
Average Activity
1
0
−1
C
Late response, gene expression
VGF
BDNF
PCSK1
DUSP4
ATP1B1
SLC7A5
NPTX1
SCG2
CREM
Percent Expressed
25
50
75
Average Expression
1
0
−1
Late response, gene activity
Percent Expressed
10
20
30
40
Average Activity
1
0
−1
−2
D
Cell Cluster
GABA
nmglut
npglut
GABA 0h
GABA 1h
GABA 6h
nmglut 0h
nmglut 1h
nmglut 6h
npglut 0h
npglut 1h
npglut 6h
0
25
50
75
100
Pseudo-time

**Fig. S6. Cell compositions and known early or late response gene expression across three time points.**

(A) Reproducible cell compositions (major subtypes of neurons) of 100 cell lines across time points. (B) Gene expression and gene activity (at TSS and gene body) of some known early response genes (ERGs) in each context (cell type × timepoint). (C) Gene expression and gene activity (at TSS and gene body) of some known late response genes (LRGs) in each context (cell type × timepoint). (D) Pseudotime trajectories of neuron stimulation of each cell type (see Methods). The direction of an arrow in each UMAP indicates late response time points along 100 bins of pseudotime points.

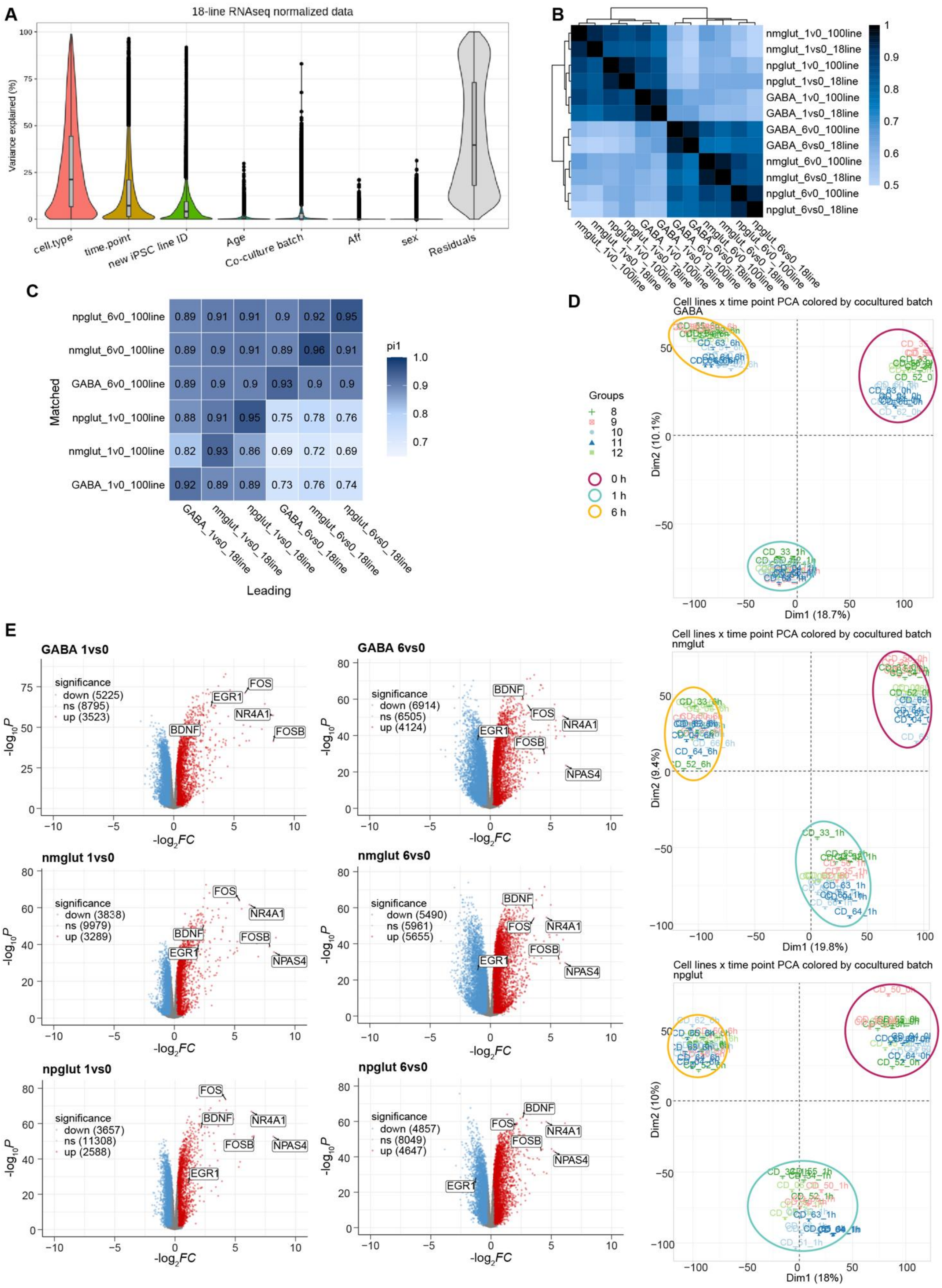
A
18-line RNAseq normalized data
Variance explained (%)
cell.type
time.point
new iPSC line ID
Age
Co-culture batch
Aff
sex
Residuals
B
nmglut_1v0_100line
nmglut_1vs0_18line
npglut_1v0_100line
npglut_1vs0_18line
GABA_1v0_100line
GABA_1vs0_18line
GABA_6v0_100line
GABA_6vs0_18line
nmglut_6v0_100line
nmglut_6vs0_18line
npglut_6v0_100line
npglut_6vs0_18line
C
pi1
Matched
Leading
GABA_1vs0_18line
nmglut_1vs0_18line
npglut_1vs0_18line
GABA_6vs0_18line
nmglut_6vs0_18line
npglut_6vs0_18line
npglut_6v0_100line 0.89 0.91 0.91 0.9 0.92 0.95
nmglut_6v0_100line 0.89 0.9 0.91 0.89 0.96 0.91
GABA_6v0_100line 0.89 0.9 0.9 0.93 0.9 0.9
npglut_1v0_100line 0.88 0.91 0.95 0.75 0.78 0.76
nmglut_1v0_100line 0.82 0.93 0.86 0.69 0.72 0.69
GABA_1v0_100line 0.92 0.89 0.89 0.73 0.76 0.74
D
Cell lines x time point PCA colored by cocultured batch
GABA
Groups
8
9
10
11
12
0 h
1 h
6 h
Dim1 (18.7%)
Dim2 (10.1%)
Cell lines x time point PCA colored by cocultured batch
nmglut
Dim1 (19.8%)
Dim2 (9.4%)
Cell lines x time point PCA colored by cocultured batch
npglut
Dim1 (18%)
Dim2 (10%)
E
GABA 1vs0
significance
down (5225)
ns (8795)
up (3523)
GABA 6vs0
significance
down (6914)
ns (6505)
up (4124)
nmglut 1vs0
significance
down (3838)
ns (9979)
up (3289)
nmglut 6vs0
significance
down (5490)
ns (5961)
up (5655)
npglut 1vs0
significance
down (3657)
ns (11308)
up (2588)
npglut 6vs0
significance
down (4857)
ns (8049)
up (4647)
FOS
EGR1
NR4A1
FOSB
BDNF
NPAS4
-log10P
-log2FC

**Fig. S7. Differentially expressed genes (DEGs) upon neuronal stimulation**.

(A) Expression variance partitioned to each cellular variable in 18-line snRNA-seq data used for main DEG analysis. Aff, schizophrenia affection status. (B) Pairwise Pearson's correlations of the $\log_2$FC of each expressed gene at each cellular context (cell type × time point) from DEG analyses in the 18 lines and in the full cohort of 100 lines. Note the strongest correlation (near 1) between 18-line and 100-line datasets for the same context. (C) Heatmap of the Pi1 analysis showing the proportion of DEGs shared between the 18 lines (leading list, *FDR* < 0.05, log2FC│>0.25 or <-0.25) and the 100 lines (matching list, *FDR* < 0.05 but no FC cut-off). (D) Principal component analysis (PCA) plots show clear separation of each sample by time point for each cell type (GABA, top; nmglut, middle; npglut, bottom). Pseudobulk RNA-seq expression values of all the expressed genes (CPM >1 in at least half of the samples) in 18 lines after correcting for co-culture batch (each batch has 2-4 iPSC lines) were used for PCA. (E) Volcano plots of DEGs at 1 h and 6 h of stimulation (vs. 0 h) in each cell type. Red, upregulated; blue, downregulated; $\log_2$FC > 0.25 or < -0.25, *FDR* < 0.05. In all analyses, pseudobulk RNA-seq data were used. Highlighted genes are some known ERGs and LRGs.

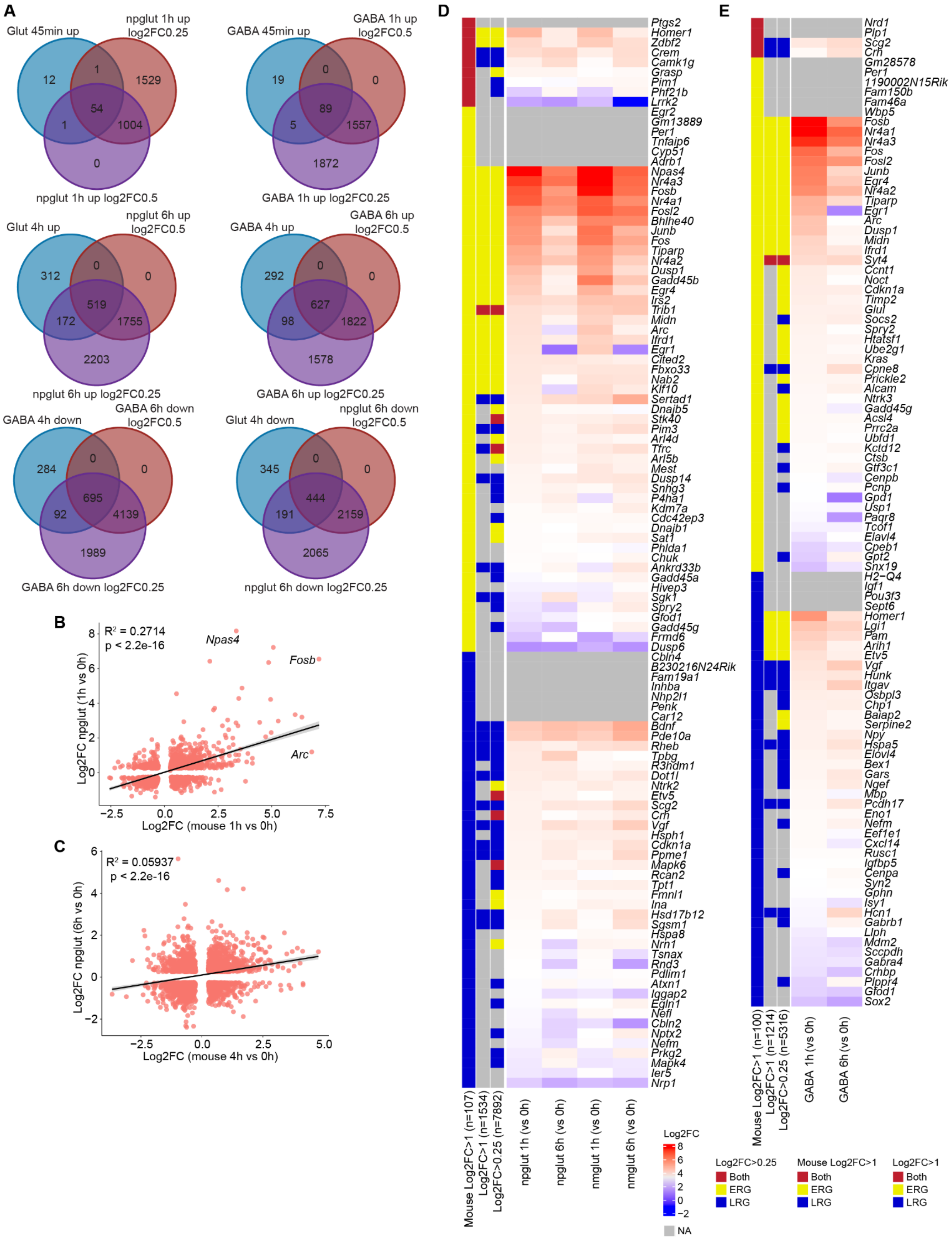
A
Glut 45min up
npglut 1h up log2FC0.25
npglut 1h up log2FC0.5
12
1
1529
54
1
1004
0
GABA 45min up
GABA 1h up log2FC0.5
GABA 1h up log2FC0.25
19
0
0
89
5
1557
1872
Glut 4h up
npglut 6h up log2FC0.5
npglut 6h up log2FC0.25
312
0
0
519
172
1755
2203
GABA 4h up
GABA 6h up log2FC0.5
GABA 6h up log2FC0.25
292
0
0
627
98
1822
1578
GABA 4h down
GABA 6h down log2FC0.5
GABA 6h down log2FC0.25
284
0
0
695
92
4139
1989
Glut 4h down
npglut 6h down log2FC0.5
npglut 6h down log2FC0.25
345
0
0
444
191
2159
2065
B
R² = 0.2714
p < 2.2e-16
Npas4
Fosb
Arc
Log2FC npglut (1h vs 0h)
Log2FC (mouse 1h vs 0h)
C
R² = 0.05937
p < 2.2e-16
Log2FC npglut (6h vs 0h)
Log2FC (mouse 4h vs 0h)
D
Ptgs2
Homer1
Zdbf2
Crem
Camk1g
Grasp
Pim1
Phf21b
Lrrk2
Egr2
Gm13889
Per1
Tnfaip6
Cyp51
Adrb1
Npas4
Nr4a3
Fosb
Nr4a1
Fosl2
Bhlhe40
Junb
Fos
Tiparp
Nr4a2
Dusp1
Gadd45b
Egr4
Irs2
Trib1
Midn
Arc
Ifrd1
Egr1
Cited2
Fbxo33
Nab2
Klf10
Sertad1
Dnajb5
Stk40
Pim3
Arl4d
Tfrc
Arl5b
Mest
Dusp14
Snhg3
P4ha1
Kdm7a
Cdc42ep3
Dnajb1
Sat1
Phlda1
Chuk
Ankrd33b
Gadd45a
Hivep3
Sgk1
Spry2
Gfod1
Gadd45g
Frmd6
Dusp6
Cbln4
B230216N24Rik
Fam19a1
Inhba
Nhp2l1
Penk
Car12
Bdnf
Pde10a
Rheb
Tpbg
R3hdm1
Dot1l
Ntrk2
Etv5
Scg2
Crh
Vgf
Hsph1
Cdkn1a
Ppme1
Mapk6
Rcan2
Tpt1
Fmnl1
Ina
Hsd17b12
Sgsm1
Hspa8
Nrn1
Tsnax
Rnd3
Pdlim1
Atxn1
Iqgap2
Egln1
Nefl
Cbln2
Nptx2
Nefm
Prkg2
Mapk4
Ier5
Nrp1
Mouse Log2FC>1 (n=107)
Log2FC>1 (n=1534)
Log2FC>0.25 (n=7892)
npglut 1h (vs 0h)
npglut 6h (vs 0h)
nmglut 1h (vs 0h)
nmglut 6h (vs 0h)
Log2FC
8
6
4
2
0
−2
NA
E
Nrd1
Plp1
Scg2
Crh
Gm28578
Per1
1190002N15Rik
Fam150b
Fam46a
Wbp5
Fosb
Nr4a1
Nr4a3
Fos
Fosl2
Junb
Egr4
Nr4a2
Tiparp
Egr1
Arc
Dusp1
Midn
Ifrd1
Syt4
Ccnt1
Noct
Cdkn1a
Timp2
Glul
Socs2
Spry2
Htatsf1
Ube2g1
Kras
Cpne8
Prickle2
Alcam
Ntrk3
Gadd45g
Acsl4
Prrc2a
Ubfd1
Kctd12
Ctsb
Gtf3c1
Cenpb
Pcnp
Gpd1
Usp1
Paqr8
Tcof1
Elavl4
Cpeb1
Gpt2
Snx19
H2−Q4
Igf1
Pou3f3
Sept6
Homer1
Lgi1
Pam
Arih1
Etv5
Vgf
Hunk
Itgav
Osbpl3
Chp1
Baiap2
Serpine2
Npy
Hspa5
Elovl4
Bex1
Gars
Ngef
Mbp
Pcdh17
Eno1
Nefm
Eef1e1
Cxcl14
Rusc1
Igfbp5
Cenpa
Syn2
Gphn
Isy1
Hcn1
Gabrb1
Liph
Mdm2
Sccpdh
Gabra4
Crhbp
Plppr4
Gfod1
Sox2
Mouse Log2FC>1 (n=100)
Log2FC>1 (n=1214)
Log2FC>0.25 (n=5316)
GABA 1h (vs 0h)
GABA 6h (vs 0h)
Log2FC>0.25
Both
ERG
LRG
Mouse Log2FC>1
Both
ERG
LRG
Log2FC>1
Both
ERG
LRG

**Fig. S8. Comparisons with other in vivo or in vitro activity-dependent expression changes.**

(A) The overlap between ERGs (45 min) and LRGs (4 h) of an independent in vitro study (*29*) and our KCl-stimulated DEGs ($log_2FC$ 0.25 or 0.5 as a cutoff) in each cell type. (B) and (C) Pearson's correlation of expression changes between KCl-stimulated npglut and mouse excitatory neurons upon in vivo electroconvulsive stimulation for ERGs and LRGs (*21, 37*), respectively. (D) and (E) Comparisons of in vivo ERGs and LRGs to that in our KCl-stimulated Glut and GABA neurons, respectively. Mouse ERGs and LRGs were defined by the authors using $log_2FC > 1$ as a cutoff at 1 h and 4 h, respectively in at least one neuronal subtype. Two types of FC cut-offs ($log_2FC > 0.25$ or $log_2FC > 1$) were used for DEGs (*FDR*<0.05) to define ERGs and LRGs in our study. ERGs, FC_1 h > FC_6 h; LRGs, FC_1 h < FC_6 h; Both, inconsistent between neuron subtypes or no difference between 1 h and 6 h. NA, downregulated, not DEG, or not expressed in our study.

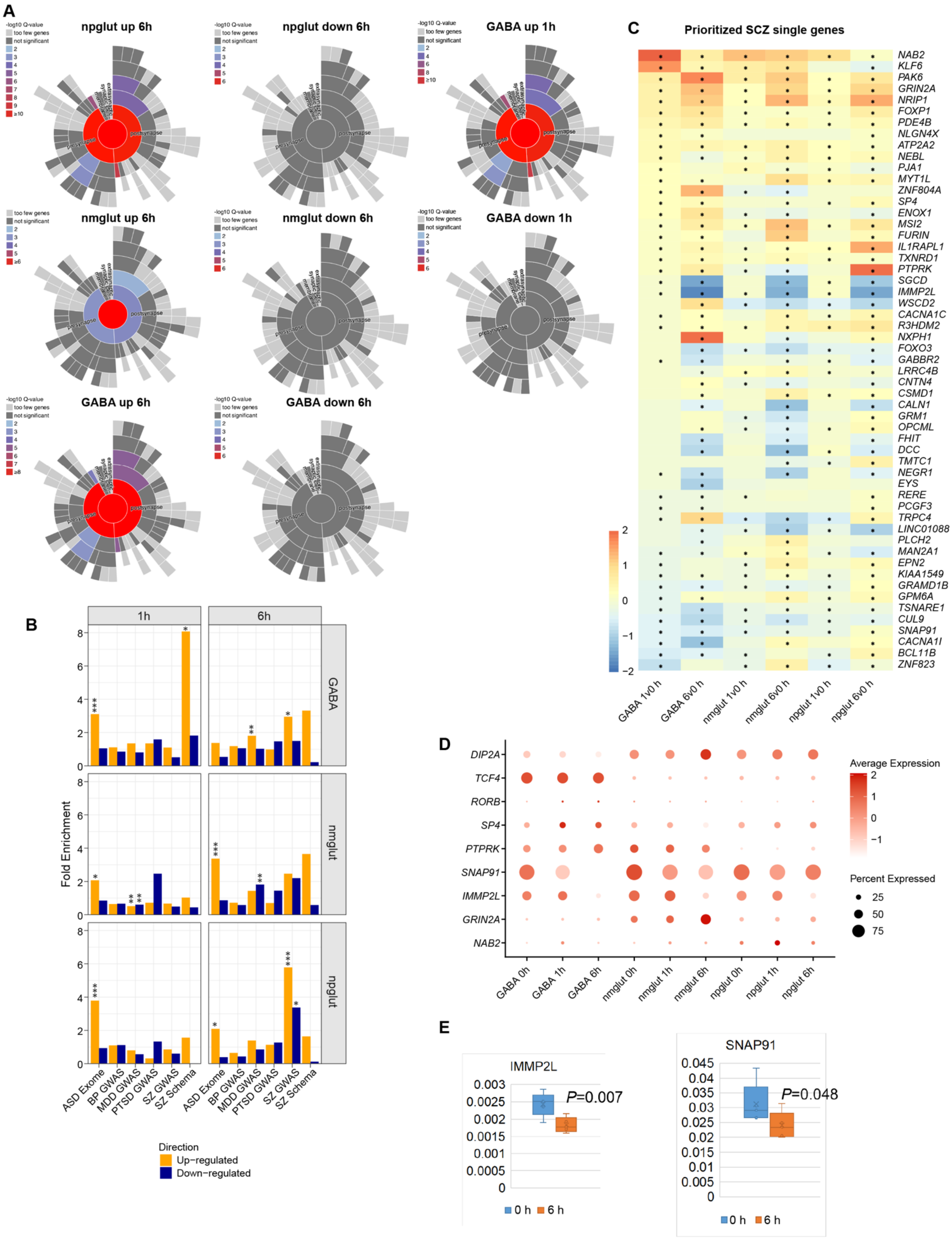
A
npglut up 6h
npglut down 6h
GABA up 1h
nmglut up 6h
nmglut down 6h
GABA down 1h
GABA up 6h
GABA down 6h
-log10 Q-value
too few genes
not significant
presynapse
postsynapse
B
1h
6h
GABA
nmglut
npglut
Fold Enrichment
ASD Exome
BP GWAS
MDD GWAS
PTSD GWAS
SZ GWAS
SZ Schema
Direction
Up−regulated
Down−regulated
C
Prioritized SCZ single genes
NAB2
KLF6
PAK6
GRIN2A
NRIP1
FOXP1
PDE4B
NLGN4X
ATP2A2
NEBL
PJA1
MYT1L
ZNF804A
SP4
ENOX1
MSI2
FURIN
IL1RAPL1
TXNRD1
PTPRK
SGCD
IMMP2L
WSCD2
CACNA1C
R3HDM2
NXPH1
FOXO3
GABBR2
LRRC4B
CNTN4
CSMD1
CALN1
GRM1
OPCML
FHIT
DCC
TMTC1
NEGR1
EYS
RERE
PCGF3
TRPC4
LINC01088
PLCH2
MAN2A1
EPN2
KIAA1549
GRAMD1B
GPM6A
TSNARE1
CUL9
SNAP91
CACNA1I
BCL11B
ZNF823
GABA 1v0 h
GABA 6v0 h
nmglut 1v0 h
nmglut 6v0 h
npglut 1v0 h
npglut 6v0 h
D
DIP2A
TCF4
RORB
SP4
PTPRK
SNAP91
IMMP2L
GRIN2A
NAB2
GABA 0h
GABA 1h
GABA 6h
nmglut 0h
nmglut 1h
nmglut 6h
npglut 0h
npglut 1h
npglut 6h
Average Expression
Percent Expressed
25
50
75
E
IMMP2L
P=0.007
SNAP91
P=0.048
0 h
6 h

**Fig. S9. Biological relevance of neuron activity-dependent DEGs.**

(A) Radial charts show the enrichment of synaptic GO terms in DEGs (from SynGO analysis) for up- or downregulated genes upon stimulation (1 h or 6 h) in each cell type. Note only stimulation-upregulated genes show enrichment of synaptic GO terms. (B) Bar plots show the enrichment of NPD GWAS genes or schizophrenia (SCZ) genes with disease-associated ultra-rare protein truncating variants (from SCHEMA study) and ASD risk genes from exome sequencing study. Up- or downregulated genes were separately analyzed for disease gene set enrichment using Fisher's exact test (compared to genes unaltered by stimulation). *: $P < 0.05$, **: $P < 0.01$, ***: $P < 0.001$. (C) $Log_2FC$ of SCZ GWAS risk genes (prioritized single credible risk gene in PGC3 SZ GWAS study) upon stimulation in each cell type. * Indicates a DEG with *FDR* <0.05. (D) Dot plot shows single neuron activity-dependent expression of some selected NPD risk genes. (E) qPCR confirmation of the mRNA expression changes of two stimulation downregulated SCZ risk genes (*IMMP2L* and *SNAP91*) in co-cultured neurons at different stages post-stimulation (using the same cDNAs for snRNA-seq; n = 3 libraries). Expression was normalized mRNA level of *GAPDH*.

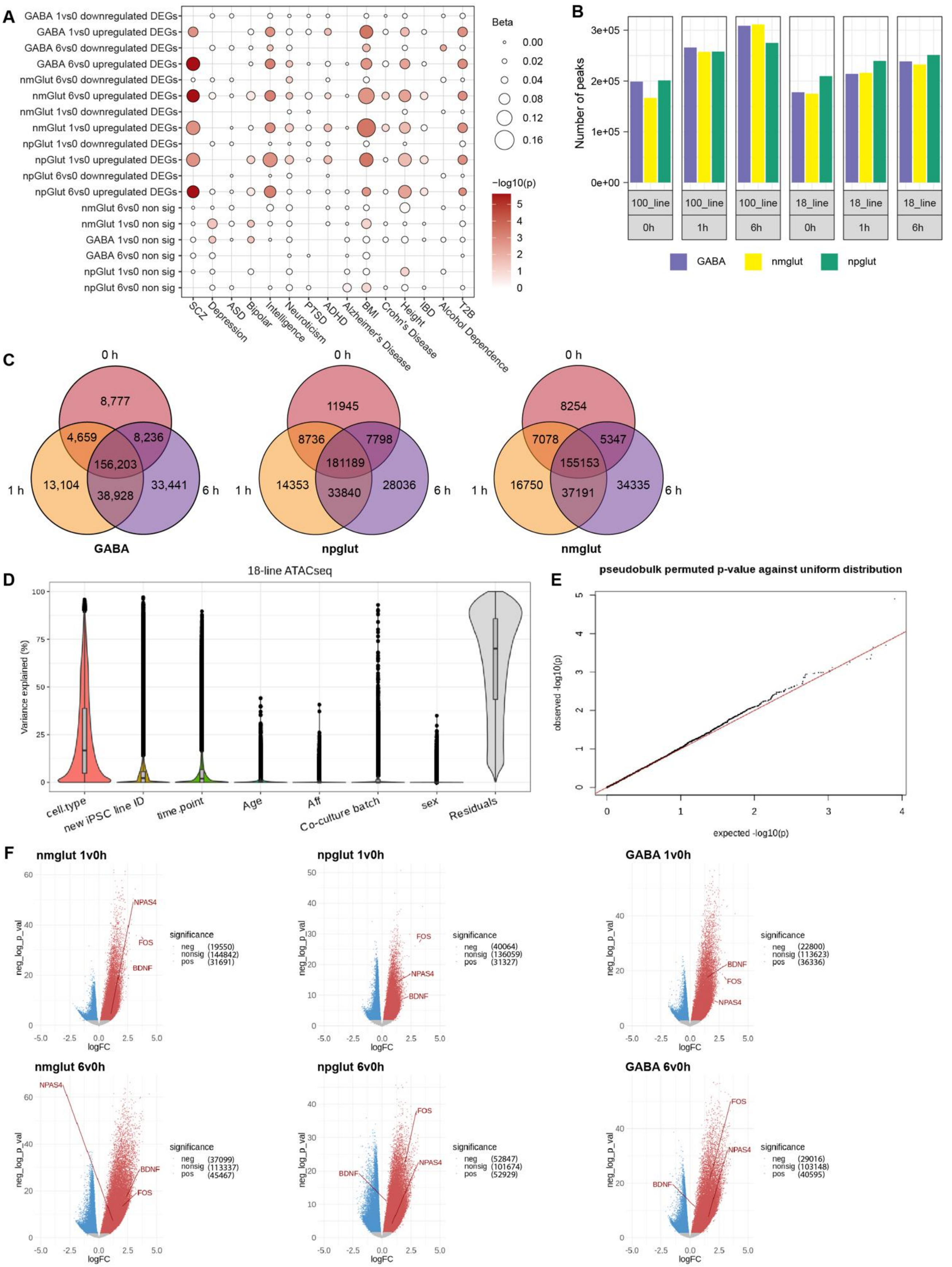
A
B
C
D
E
F
Beta
-log10(p)
Number of peaks
GABA
nmglut
npglut
18-line ATACseq
Variance explained (%)
pseudobulk permuted p-value against uniform distribution
observed -log10(p)
expected -log10(p)
nmglut 1v0h
npglut 1v0h
GABA 1v0h
nmglut 6v0h
npglut 6v0h
GABA 6v0h
neg_log_p_val
logFC
significance

**Fig. S10. GWAS enrichment of differentially expressed genes and the mapping of differentially accessible (DA) OCR peak.**

(A) GWAS enrichment of differentially expressed genes (DEGs; 1 h vs. 0 h and 6 h vs. 0 h) sets. Several non-NPD traits were included as control in MAGMA analysis. IBD, inflammatory bowel disease; T2D, type 2 diabetes. (B) Peak counts of major neuron subtypes at different post-stimulation time points using the 18-line or the 100-line dataset. Note the comparable number of peaks called from the two datasets. (C) Venn diagram of OCR peaks in iGABA, npglut, and nmglut at three time points. (D) Violin plot shows variances partitioned to each cellular variable in the 18-line snATAC-seq dataset. Aff, schizophrenia affection status. (E) Q-Q plot shows the distribution of expected (x, assuming uniform distribution) against the permutated values ($-\log_{10}P$) of DA peaks. Note the minimal *p*-value inflation in our DA peak analysis. (F) Volcano plots of DA peaks in each cell type upon stimulation. Red dots, upregulated peaks; blue dots, downregulated peaks; $FDR < 0.05$ as cut-off.

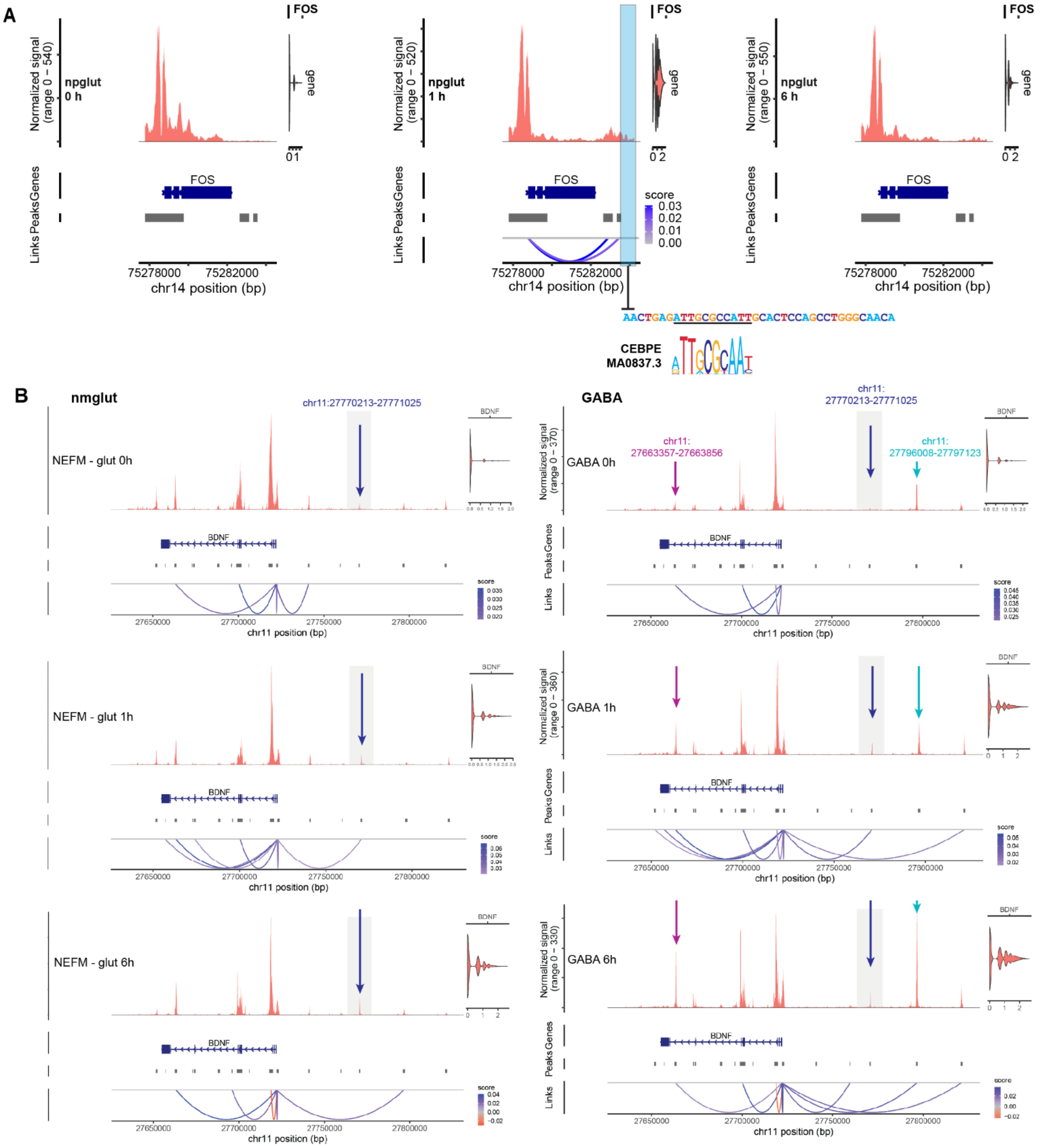

**Fig. S11. Activity-dependent OCR peaks at *FOS* and *BDNF* loci.**

(A) Putative OCR peaks that mediates FOS expression in early response. From left to right, panels showing OCR peaks, gene expression, and peak-gene linkage at 0 h, 1 h, and 6 h, respectively. Note the specific peak-gene linkage was only observed at 1 h post-stimulation and the presence of CEBPE-binding motif in the 3' OCR of FOS. Only data from npglut is shown. (B) The DA peak (dark blue arrow) showed weak accessibility in nmglut (left panel) and in GABA (right panel), but with stronger peak accessibility in npglut (see Fig. S12). Light blue arrow points to a GABA-specific DA peak. Note the same specific peak-gene linkage as in npglut (see Fig. S12) only presents at 1 h in both nmglut and GABA.

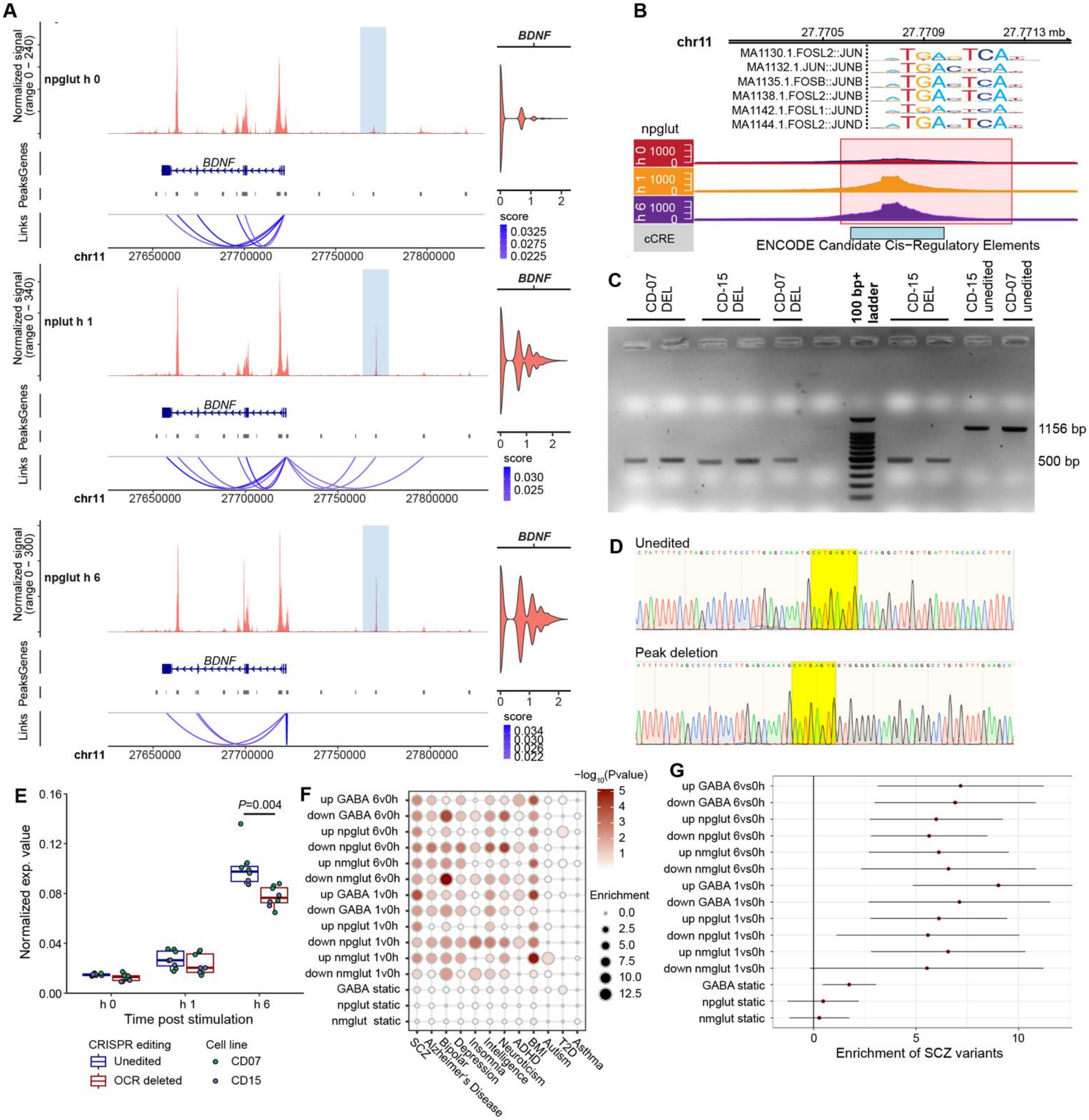

**Fig. S12. CRISPR/Cas9 deletion a stimulation-specific *BDNF* OCR peak and overall DA peak enrichment for NPD GWAS risk.**

(A) Peak-gene linkage plot of *BDNF* in npglut. In each of the three panels, the three tracks from the top to bottom are ATAC-seq signal, gene annotation, and peak-to-gene links. The highlighted is the most-upregulated peak and its gene-peak linkage was observed only at 1 h. (B) The highlighted peak in (A) overlaps with ENCODE CRE and has AP1 TF-binding sites. (C) The boxed OCR region in (B) was deleted by CRISPR/Cas9 editing and confirmed by PCR. Note the ~650 bp reduction in fragment size on 1% agarose gel in CRISPR-edited lines (vs. unedited lines).

Two donor lines (CD07, CD15) were used for editing. Each lane represents a different isogenic clone before or after editing. (D) Sanger sequencing confirmation of unedited and CRISPR-edited line carrying the *BDNF* peak deletion (homozygous). The PAM sequence is highlighted in yellow. (E) *BDNF* peak deletion affected *BDNF* mRNA expression (assayed by qPCR) in iGlut at 6 h post-stimulation. n = 3-5 independent cultures of 2 donor lines (CD07 and CD15). Unpaired two-side Student's *t*-test. (F) Stratified-LDSC analysis of GWAS SNP heritability enrichment in dynamic peak sets (up- or downregulated) for GWAS of several NPD and non-NPD control disorders. Shown are fold of enrichment (bubble size) and significance (color scheme, $-\log_{10}P$). Background SNPs are used as control for enrichment test. (G) SCZ heritability enrichment analysis from (F) for up- or downregulated, or static (unchanged) OCR peaks.

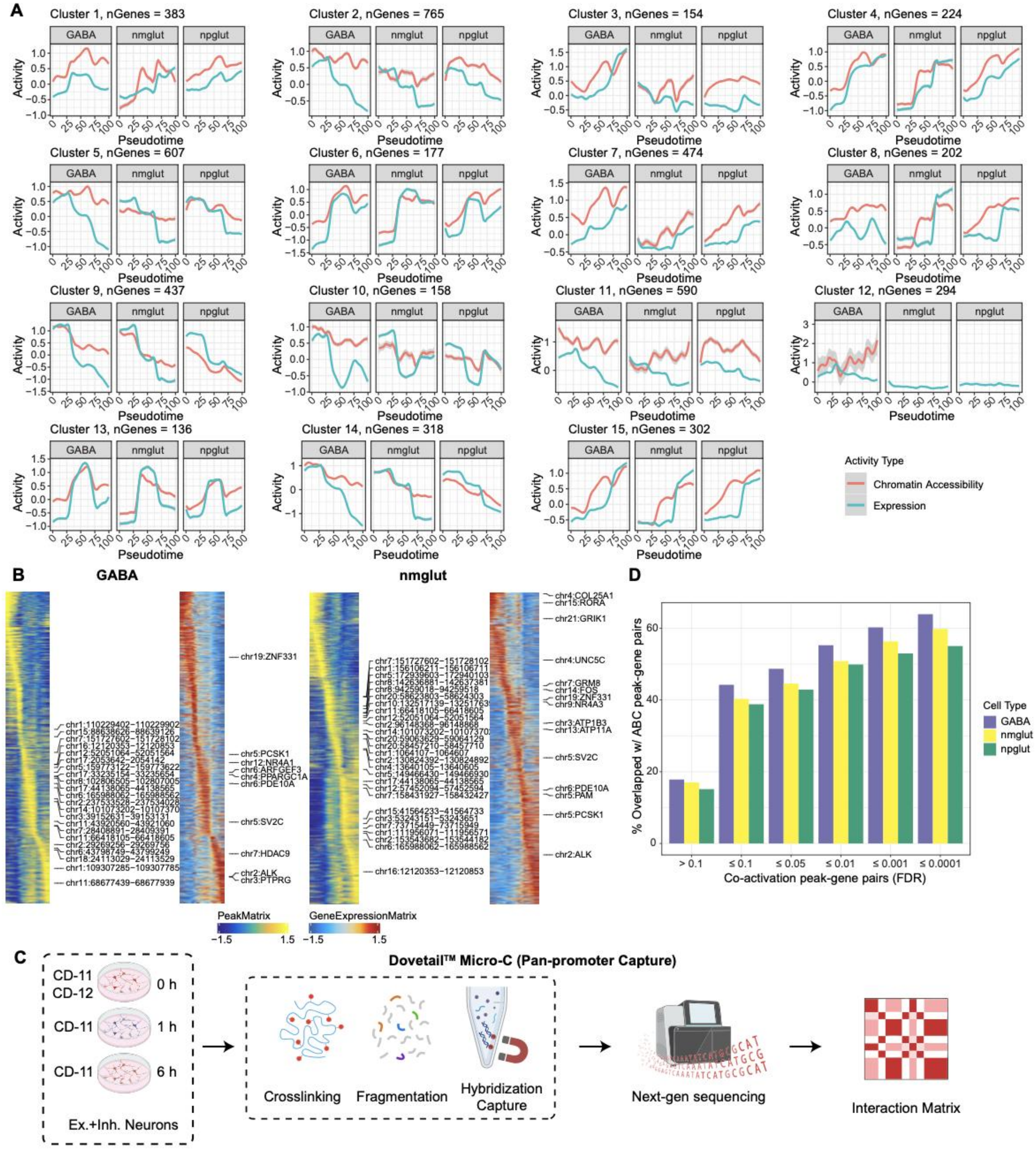

**Fig. S13. Neuron activity-dependent gene expression modules (clusters) and correlation between chromatin accessibility and gene expression for OCR-gene pairs.**

(A) Pseudotime trajectories of normalized chromatin accessibility (red) and gene expression (blue) for all expression clusters. Added LOESS smoothing for better visualization. (B) Heatmap of pseudotime activity of chromatin accessibility and gene expression for identified OCR-gene pairs in GABA and nmglut. For each cell type, the left panel depicts normalized chromatin accessibility of mapped peaks, while the right panel illustrates normalized gene expression. Highlighted are most variable peak-gene pairs. (C) Schematics of the Dovetail Micro-C (Pan promoter capture) experimental design. The co-cultured excitatory and inhibitory neurons before and after KCl stimulation (as in Figure 1A) were processed for Micro-C via three major steps (crosslinking, fragmentation, and hybridization capture), followed by sequencing and data analyses to identify chromatin interaction bins (using CHICAGO). Neuron co-cultures (both iGlut and GABA) of two iPSC lines (CD11, CD12) were used, of which CD11 line has data from all three time points (0 h, 1 h, and 6 h) of KCl stimulation. (D) Proportion of peak-gene pairs overlapped with ABC-defined peak-gene contacts at different *FDR* cutoffs.

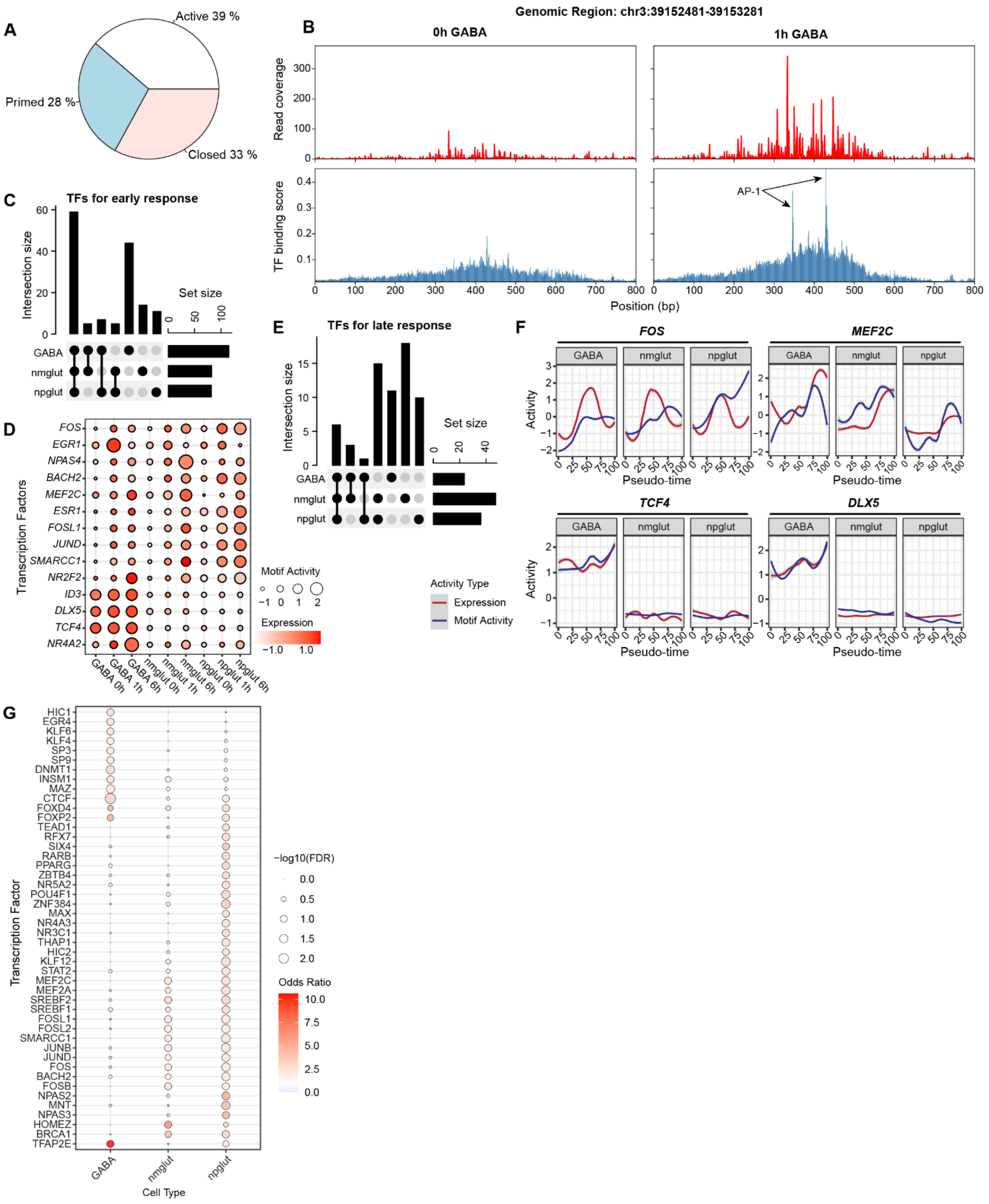
A
Active 39 %
Primed 28 %
Closed 33 %
B
Genomic Region: chr3:39152481-39153281
0h GABA
1h GABA
Read coverage
TF binding score
AP-1
Position (bp)
C
TFs for early response
Intersection size
Set size
GABA
nmglut
npglut
D
Transcription Factors
FOS
EGR1
NPAS4
BACH2
MEF2C
ESR1
FOSL1
JUND
SMARCC1
NR2F2
ID3
DLX5
TCF4
NR4A2
GABA 0h
GABA 1h
GABA 6h
nmglut 0h
nmglut 1h
nmglut 6h
npglut 0h
npglut 1h
npglut 6h
Motif Activity
Expression
E
TFs for late response
Intersection size
Set size
GABA
nmglut
npglut
Activity Type
Expression
Motif Activity
F
FOS
MEF2C
TCF4
DLX5
GABA
nmglut
npglut
Activity
Pseudo-time
G
Transcription Factor
HIC1
EGR4
KLF6
KLF4
SP3
SP9
DNMT1
INSM1
MAZ
CTCF
FOXD4
FOXP2
TEAD1
RFX7
SIX4
RARB
PPARG
ZBTB4
NR5A2
POU4F1
ZNF384
MAX
NR4A3
NR3C1
THAP1
HIC2
KLF12
STAT2
MEF2C
MEF2A
SREBF2
SREBF1
FOSL1
FOSL2
SMARCC1
JUNB
JUND
FOS
BACH2
FOSB
NPAS2
MNT
NPAS3
HOMEZ
BRCA1
TFAP2E
GABA
nmglut
npglut
Cell Type
−log10(FDR)
Odds Ratio

**Fig. S14. TF regulation of early and late neuronal response and ASD-related gene regulatory network (GRN).**

(A) and (B) Characterization of chromatin states (open chromatin and H3K27ac mark) for "responsive OCRs" ($FDR \leq 0.05$ and $\log_2 FC > 1$) upon KCI stimulation. (A) Breakdown of chromatin states at 0 h. Primed = partially accessible but no H3K27ac mark. (B) Chromatin state changes upon stimulation at *CSRNP1* locus in GABA cells. Red bars, number of ATAC-seq reads; blue bars, TF binding scores at each position, inferred from seq2PRINT. Note the strong binding sites of AP-1 family motif at 1 h. (C) Number of cell-type-specific and shared candidate TFs regulating early responses. (D) Motif activity (bubble size) and expression (color) of selected TFs across time points and cell types. Motif activities are in a Z-score scale. (E) Number of cell-type-specific and shared candidate TFs regulating late responses. (F) Pseudotime trajectories showing expression (red) and motif activity (blue) of FOS, MEF2C, TCF4, and DLX5. (G) TFs in the reconstructed GRNs that have targets enriched for ASD risk genes ($FDR \leq 0.1$, Fisher's exact test).

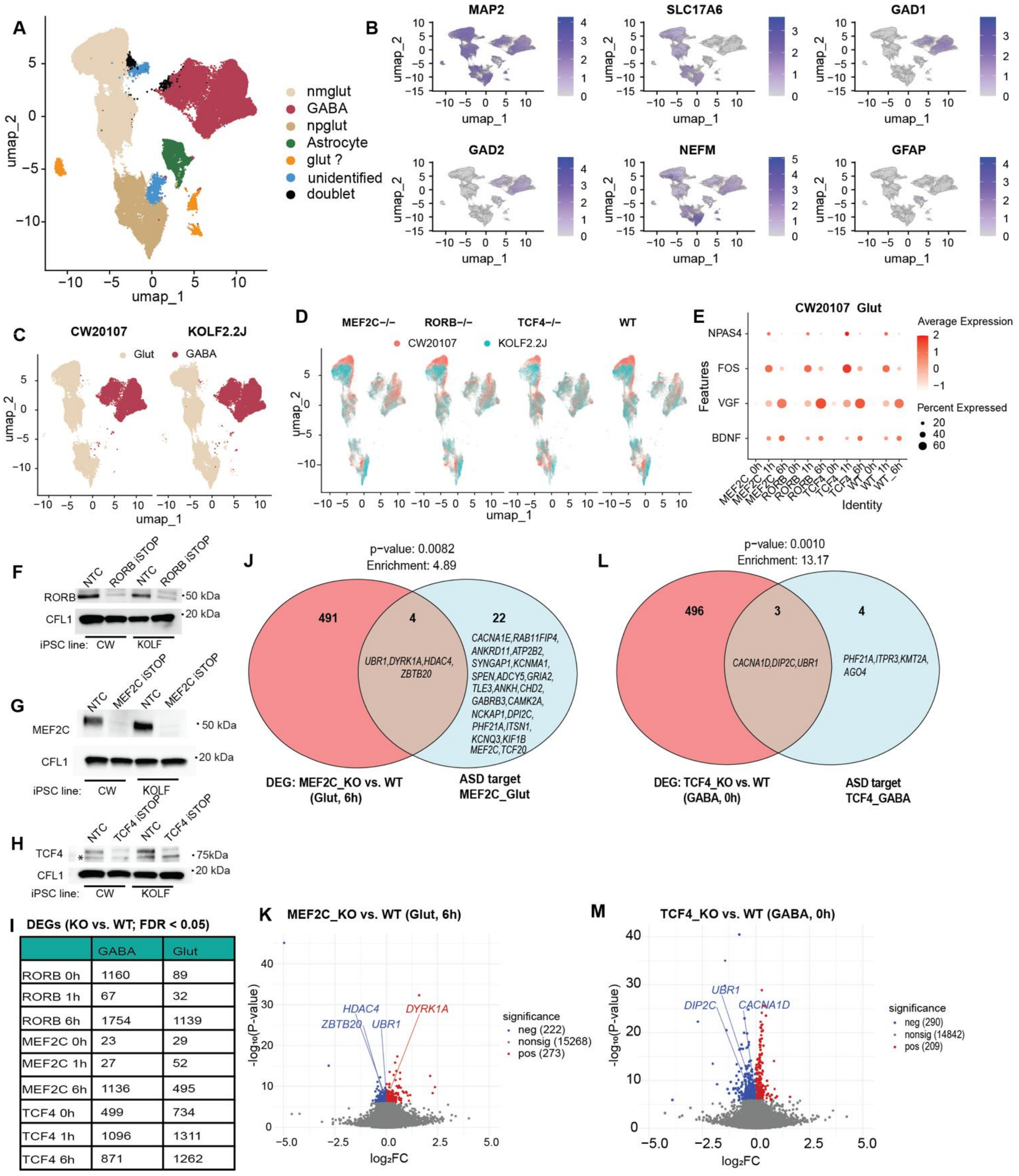

I DEGs (KO vs. WT; FDR < 0.05)

| | GABA | Glut |
|---|---|---|
| RORB 0h | 1160 | 89 |
| RORB 1h | 67 | 32 |
| RORB 6h | 1754 | 1139 |
| MEF2C 0h | 23 | 29 |
| MEF2C 1h | 27 | 52 |
| MEF2C 6h | 1136 | 495 |
| TCF4 0h | 499 | 734 |
| TCF4 1h | 1096 | 1311 |
| TCF4 6h | 871 | 1262 |

**Fig. S15. The ASD subnetwork validation by TF knockout (KO) in KCl-stimulated neural co-cultures.**

(A) Integrated scRNA-seq UMAP for iGlut and GABA neurons co-culture derived from WT and TF-KO lines (*MEF2C*-/-, *RORB-/-,* and *TCF4-/-*). TF-KO was carried out in iPSC lines by introducing premature stop codons (iSTOP) for each TF. Neuronal co-cultures and KCI stimulation were conducted in the same way as the main neuron stimulation experiment. Cell type assignment was followed the same procedures as our neuron-stimulation scRNA-seq in Figs. S1-4. (B) Feature plots confirming similar cell subtype-clustering (npglut, nmglut, and GABA) to our initial neuron stimulation experiment. (C) and (D) UMAPs showing reproducibility between the two donor lines (CW20107 and KOLF2.2J), and across WT and TF-KO lines (*MEF2C*-/-, *RORB-/-,* and *TCF4-/-*), respectively. (E) Confirmation of effective neuron activation upon KCl stimulation by examining expression dynamics of some ERGs and LRGs (only Glut from CW20107 line shown). (F) to (H) Western blot confirmation of TF-KO (LoF via iSTOP) for RORB (F), MEF2C (G) and TCF4 (H). NTC (non-transfected control, i.e., no editing) line in CRISPR editing. CFL1 was used as endogenous control for normalizing TF expression to confirm TF KO. * In (H) indicates a nonspecific protein band. (I) Number of DEGs identified in each context between TF-KO and WT lines. MAST was used for scRNA-seq DEG analysis with cell line modeled as random effects. (J) and (K) DEGs in TF-KO neurons (Glut at 6 h) (vs. WT) overlapping with the predicted ASD-associated target genes and the volcano plot showing the overlapping DEGs for MEF2C, respectively. (L) and (M) DEGs in TF-KO neurons (GABA at 0 h) (vs. WT) overlapping with the predicted ASD-associated target genes and the volcano plot showing the overlapping DEGs for TCF4, respectively. For (J) and (L), the overlap with the predicted ASD target genes of each TF was shown for the matched cell subtypes. Enrichment *P* values were derived from hypergeometric tests.

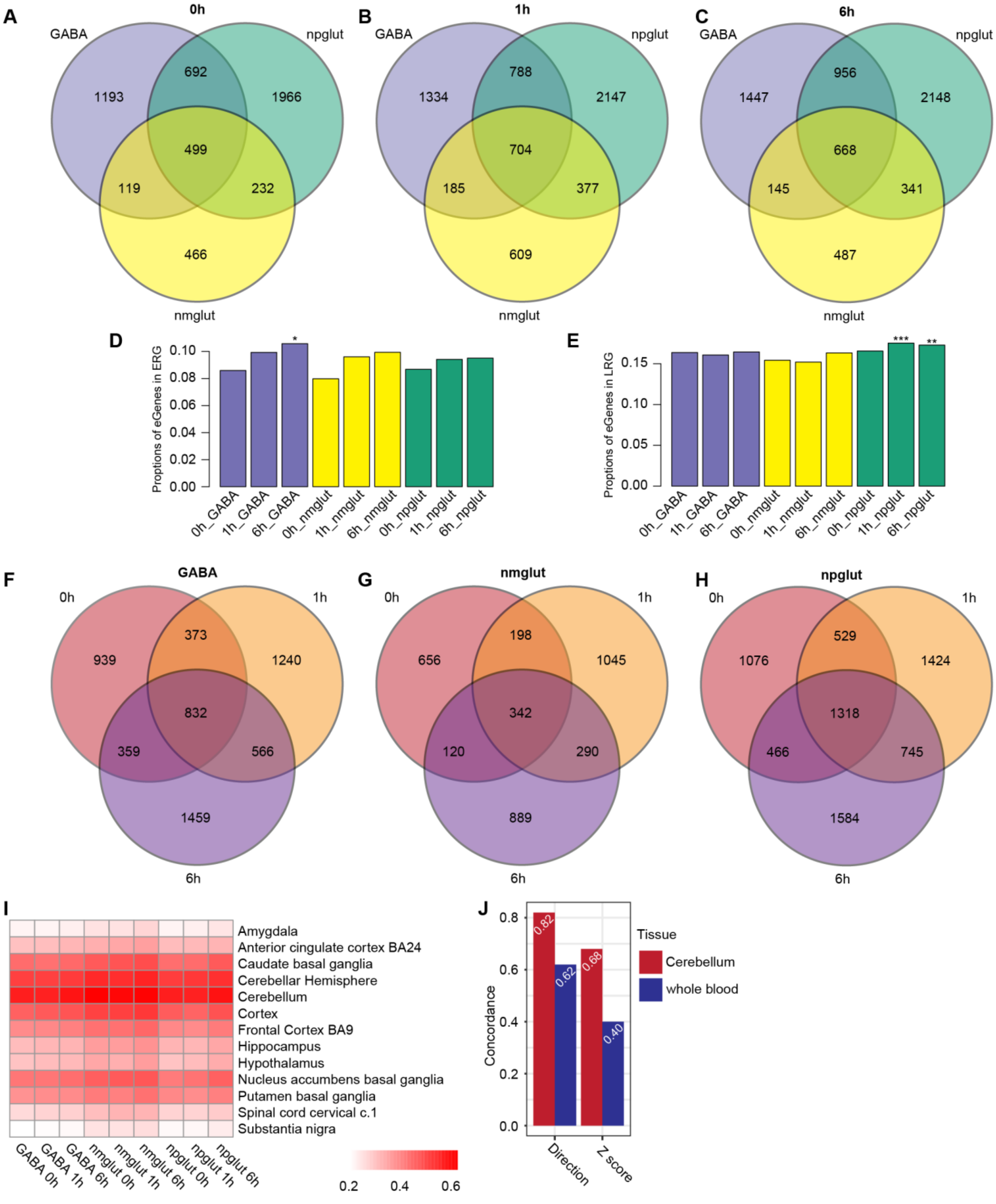

**Fig. S16. Neuron activation-dependent eQTL mapping and characterization.**

(A) to (C) Venn diagram of eGenes of the three cell types at 0 h, 1 h, and 6 h of KCl stimulation, respectively. (D) and (E) Proportions of eGenes that are also found to be in vivo ERGs and

LRGs (*21*), respectively. (F) to (H) Venn diagram of eGenes at the three time points for GABA, nmglut, and npglut, respectively. (I) Proportions of neuron activity eQTL shared with GTEx brains from Pi1 analysis. (J) Comparison of eQTL effects between 0 h nmglut and GTEx cerebellum or whole blood. The "Direction" bar, concordance of the signs of effects. The "Z-score" bar, effect size correlation.

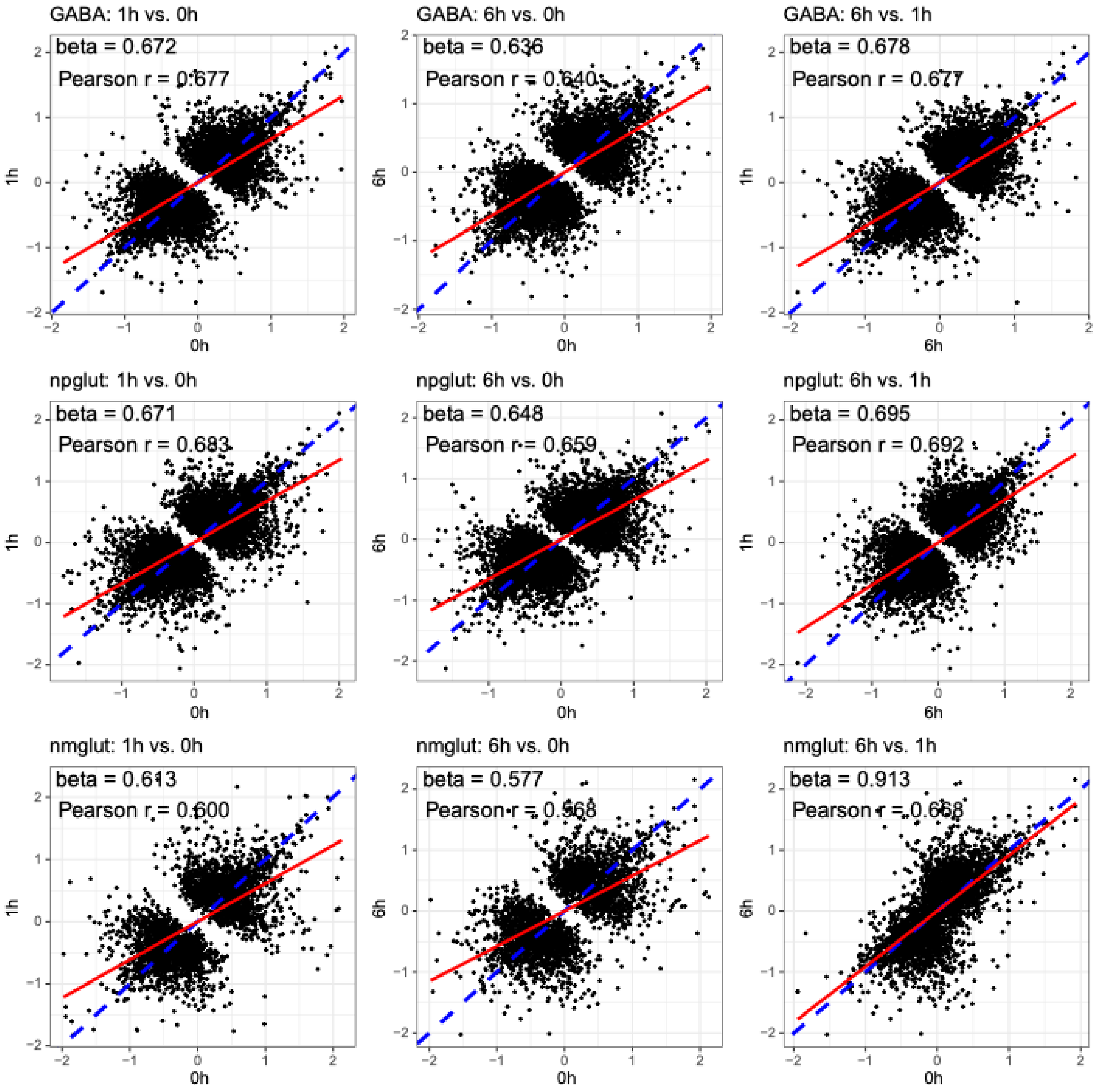

**Fig. S17. Regression plot of the effect size correlation between time points for each cell type.**

Blue dashed line is diagonal line with slope=1. Red line is fitted to the effect sizes. Regression β and Pearson's r are listed in each panel.

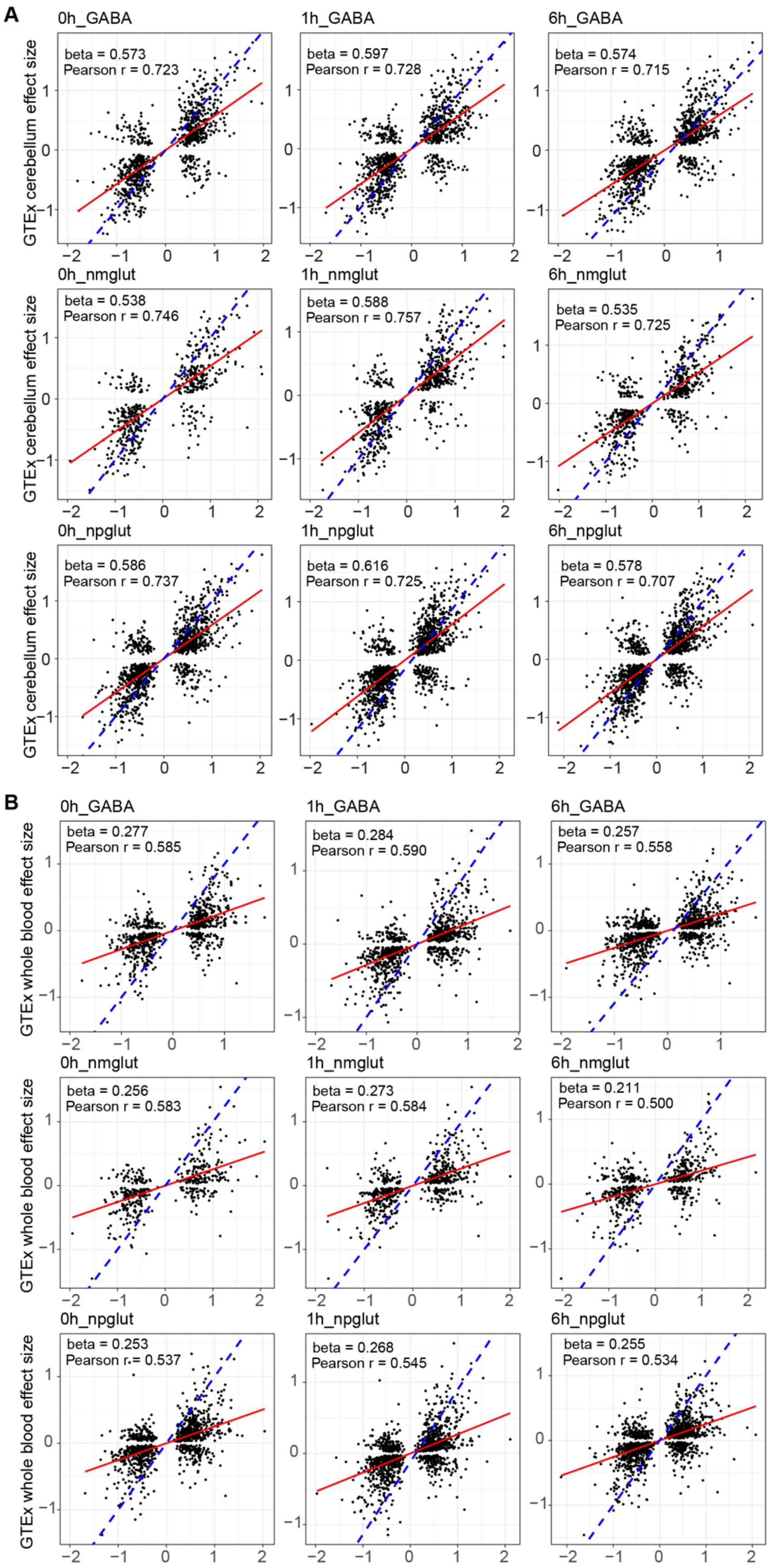
A
0h_GABA
beta = 0.573
Pearson r = 0.723
1h_GABA
beta = 0.597
Pearson r = 0.728
6h_GABA
beta = 0.574
Pearson r = 0.715
0h_nmglut
beta = 0.538
Pearson r = 0.746
1h_nmglut
beta = 0.588
Pearson r = 0.757
6h_nmglut
beta = 0.535
Pearson r = 0.725
0h_npglut
beta = 0.586
Pearson r = 0.737
1h_npglut
beta = 0.616
Pearson r = 0.725
6h_npglut
beta = 0.578
Pearson r = 0.707
GTEx cerebellum effect size
B
0h_GABA
beta = 0.277
Pearson r = 0.585
1h_GABA
beta = 0.284
Pearson r = 0.590
6h_GABA
beta = 0.257
Pearson r = 0.558
0h_nmglut
beta = 0.256
Pearson r = 0.583
1h_nmglut
beta = 0.273
Pearson r = 0.584
6h_nmglut
beta = 0.211
Pearson r = 0.500
0h_npglut
beta = 0.253
Pearson r = 0.537
1h_npglut
beta = 0.268
Pearson r = 0.545
6h_npglut
beta = 0.255
Pearson r = 0.534
GTEx whole blood effect size

**Fig. S18. Regression plots of the effect sizes correlation between our neuronal activity eQTL and GTEx eQTL.**

(A) Correlation with GTEx brain cerebellum eQTL, (B) Correlation with GTEx whole blood eQTL. Blue dashed line is diagonal line with slope=1. Red line is fitted to the effect sizes. Regression $\beta$ and Pearson's r are listed in each panel.

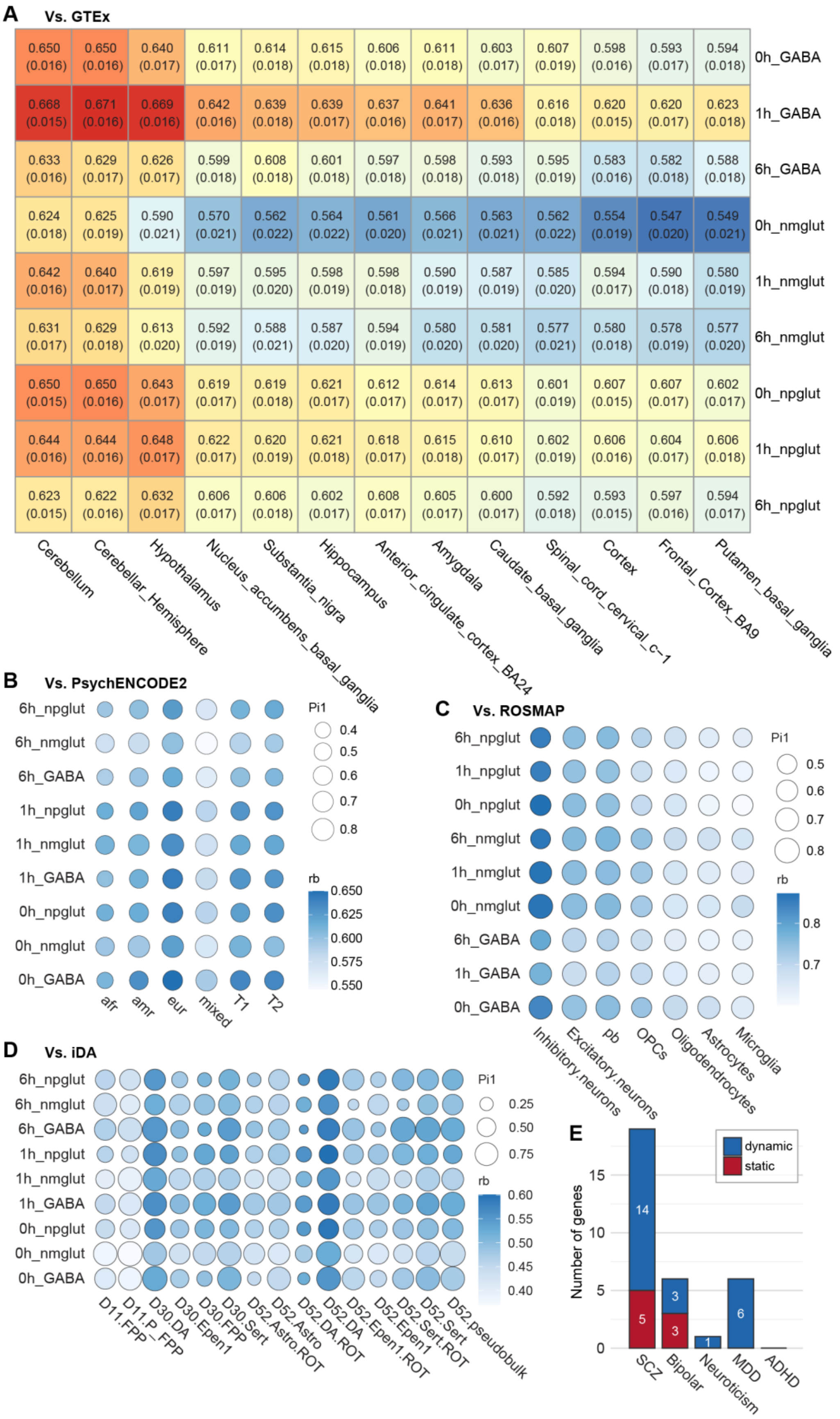
A
Vs. GTEx
0.650 (0.016) 0.650 (0.016) 0.640 (0.017) 0.611 (0.017) 0.614 (0.018) 0.615 (0.018) 0.606 (0.018) 0.611 (0.018) 0.603 (0.017) 0.607 (0.019) 0.598 (0.016) 0.593 (0.017) 0.594 (0.018) 0h_GABA
0.668 (0.015) 0.671 (0.016) 0.669 (0.016) 0.642 (0.016) 0.639 (0.018) 0.639 (0.017) 0.637 (0.016) 0.641 (0.017) 0.636 (0.016) 0.616 (0.018) 0.620 (0.015) 0.620 (0.017) 0.623 (0.018) 1h_GABA
0.633 (0.016) 0.629 (0.017) 0.626 (0.017) 0.599 (0.018) 0.608 (0.018) 0.601 (0.018) 0.597 (0.018) 0.598 (0.018) 0.593 (0.018) 0.595 (0.019) 0.583 (0.016) 0.582 (0.018) 0.588 (0.018) 6h_GABA
0.624 (0.018) 0.625 (0.019) 0.590 (0.021) 0.570 (0.021) 0.562 (0.022) 0.564 (0.022) 0.561 (0.020) 0.566 (0.021) 0.563 (0.021) 0.562 (0.022) 0.554 (0.019) 0.547 (0.020) 0.549 (0.021) 0h_nmglut
0.642 (0.016) 0.640 (0.017) 0.619 (0.019) 0.597 (0.019) 0.595 (0.020) 0.598 (0.019) 0.598 (0.018) 0.590 (0.019) 0.587 (0.019) 0.585 (0.020) 0.594 (0.017) 0.590 (0.018) 0.580 (0.019) 1h_nmglut
0.631 (0.017) 0.629 (0.018) 0.613 (0.020) 0.592 (0.019) 0.588 (0.021) 0.587 (0.020) 0.594 (0.019) 0.580 (0.020) 0.581 (0.020) 0.577 (0.021) 0.580 (0.018) 0.578 (0.019) 0.577 (0.020) 6h_nmglut
0.650 (0.015) 0.650 (0.016) 0.643 (0.017) 0.619 (0.017) 0.619 (0.018) 0.621 (0.017) 0.612 (0.017) 0.614 (0.017) 0.613 (0.017) 0.601 (0.019) 0.607 (0.015) 0.607 (0.017) 0.602 (0.017) 0h_npglut
0.644 (0.016) 0.644 (0.016) 0.648 (0.017) 0.622 (0.017) 0.620 (0.019) 0.621 (0.018) 0.618 (0.017) 0.615 (0.018) 0.610 (0.017) 0.602 (0.019) 0.606 (0.016) 0.604 (0.017) 0.606 (0.018) 1h_npglut
0.623 (0.015) 0.622 (0.016) 0.632 (0.017) 0.606 (0.017) 0.606 (0.018) 0.602 (0.017) 0.608 (0.017) 0.605 (0.017) 0.600 (0.017) 0.592 (0.018) 0.593 (0.015) 0.597 (0.016) 0.594 (0.017) 6h_npglut
Cerebellum
Cerebellar_Hemisphere
Hypothalamus
Nucleus_accumbens_basal_ganglia
Substantia_nigra
Hippocampus
Anterior_cingulate_cortex_BA24
Amygdala
Caudate_basal_ganglia
Spinal_cord_cervical_c-1
Cortex
Frontal_Cortex_BA9
Putamen_basal_ganglia
rb
0.66
0.64
0.62
0.6
0.58
0.56
B
Vs. PsychENCODE2
6h_npglut
6h_nmglut
6h_GABA
1h_npglut
1h_nmglut
1h_GABA
0h_npglut
0h_nmglut
0h_GABA
afr
amr
eur
mixed
T1
T2
Pi1
0.4
0.5
0.6
0.7
0.8
rb
0.650
0.625
0.600
0.575
0.550
C
Vs. ROSMAP
6h_npglut
1h_npglut
0h_npglut
6h_nmglut
1h_nmglut
0h_nmglut
6h_GABA
1h_GABA
0h_GABA
Inhibitory.neurons
Excitatory.neurons
pb
OPCs
Oligodendrocytes
Astrocytes
Microglia
Pi1
0.5
0.6
0.7
0.8
rb
0.8
0.7
D
Vs. iDA
6h_npglut
6h_nmglut
6h_GABA
1h_npglut
1h_nmglut
1h_GABA
0h_npglut
0h_nmglut
0h_GABA
D11.FPP
D11.P_FPP
D30.DA
D30.Epen1
D30.FPP
D30.Sert
D52.Astro
D52.Astro.ROT
D52.DA
D52.DA.ROT
D52.Epen1
D52.Epen1.ROT
D52.Sert
D52.Sert.ROT
D52.pseudobulk
Pi1
0.25
0.50
0.75
rb
0.60
0.55
0.50
0.45
0.40
E
dynamic
static
Number of genes
0
5
10
15
14
5
3
3
1
6
SCZ
Bipolar
Neuroticism
MDD
ADHD

**Fig. S19. Comparisons between our KCl-stimulated eQTL with other eQTL datasets.**

(A) eQTL effect size correlations, using the Rb estimator, for GTEx eQTL of different brain regions (*55*). (B) Comparison with PsychENCODE2 eQTL (*56*). Size and color of the bubbles represent estimated proportion of shared eQTL (Pi1), and the Rb estimated correlation, respectively. “afr”, “amr”, “eur”, and “mixed” represent ancestries of donors disregarding the prenatal periods. “T1” and “T2” represent trimesters of pregnancy when samples were taken. (C) Comparison with ROSMAP eQTL of different brain cell types (*57*). Pb, peripheral blood cells; OPC, oligodendrocyte precursor cells. (D) Comparison with scRNA-seq eQTL from iPSC-derived Dopaminergic (iDA) neurons (*58*). D11, D30, and D52 indicate the days of DA neuron differentiation. DA, dopaminergic neurons; FPP, floor plate progenitors; Epen1, ependymal-like 1; Sert, serotonergic-like neurons; ROT, rotenone treatment. (E) The number of risk genes of high-confidence (PIP > 0.8) identified in cTWAS for the five NPD phenotypes. “Dynamic” or “Static” according to how PIPs are partitioned across contexts (see Fig. 4D caption).

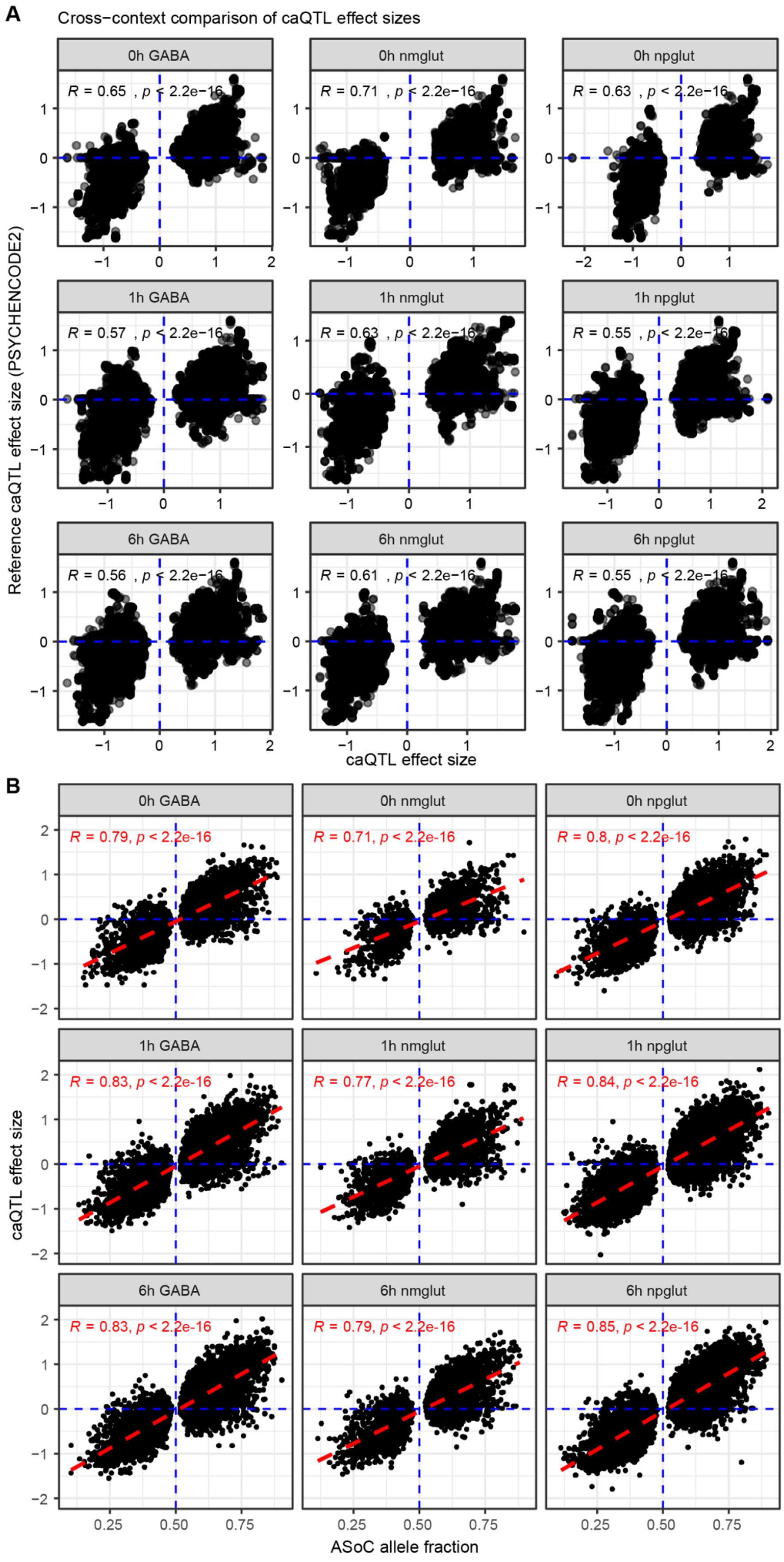
A
Cross−context comparison of caQTL effect sizes
0h GABA
0h nmglut
0h npglut
1h GABA
1h nmglut
1h npglut
6h GABA
6h nmglut
6h npglut
R = 0.65 , p < 2.2e−16
R = 0.71 , p < 2.2e−16
R = 0.63 , p < 2.2e−16
R = 0.57 , p < 2.2e−16
R = 0.63 , p < 2.2e−16
R = 0.55 , p < 2.2e−16
R = 0.56 , p < 2.2e−16
R = 0.61 , p < 2.2e−16
R = 0.55 , p < 2.2e−16
Reference caQTL effect size (PSYCHENCODE2)
caQTL effect size
B
R = 0.79, p < 2.2e-16
R = 0.71, p < 2.2e-16
R = 0.8, p < 2.2e-16
R = 0.83, p < 2.2e-16
R = 0.77, p < 2.2e-16
R = 0.84, p < 2.2e-16
R = 0.83, p < 2.2e-16
R = 0.79, p < 2.2e-16
R = 0.85, p < 2.2e-16
caQTL effect size
ASoC allele fraction

**Fig. S20. Strong correlation of the effect size between our neuron activity caQTL and other caQTLs.**

(A) Cross-context comparison of neuron activity caQTL effect sizes with reference PsychENCODE2 brain caQTLs. (B) Cross-context comparison of neuron activity caQTL effect sizes with our ASoC (*FDR* < 0.05) allelic fractions. Pearson's R and P values are listed in each panel.

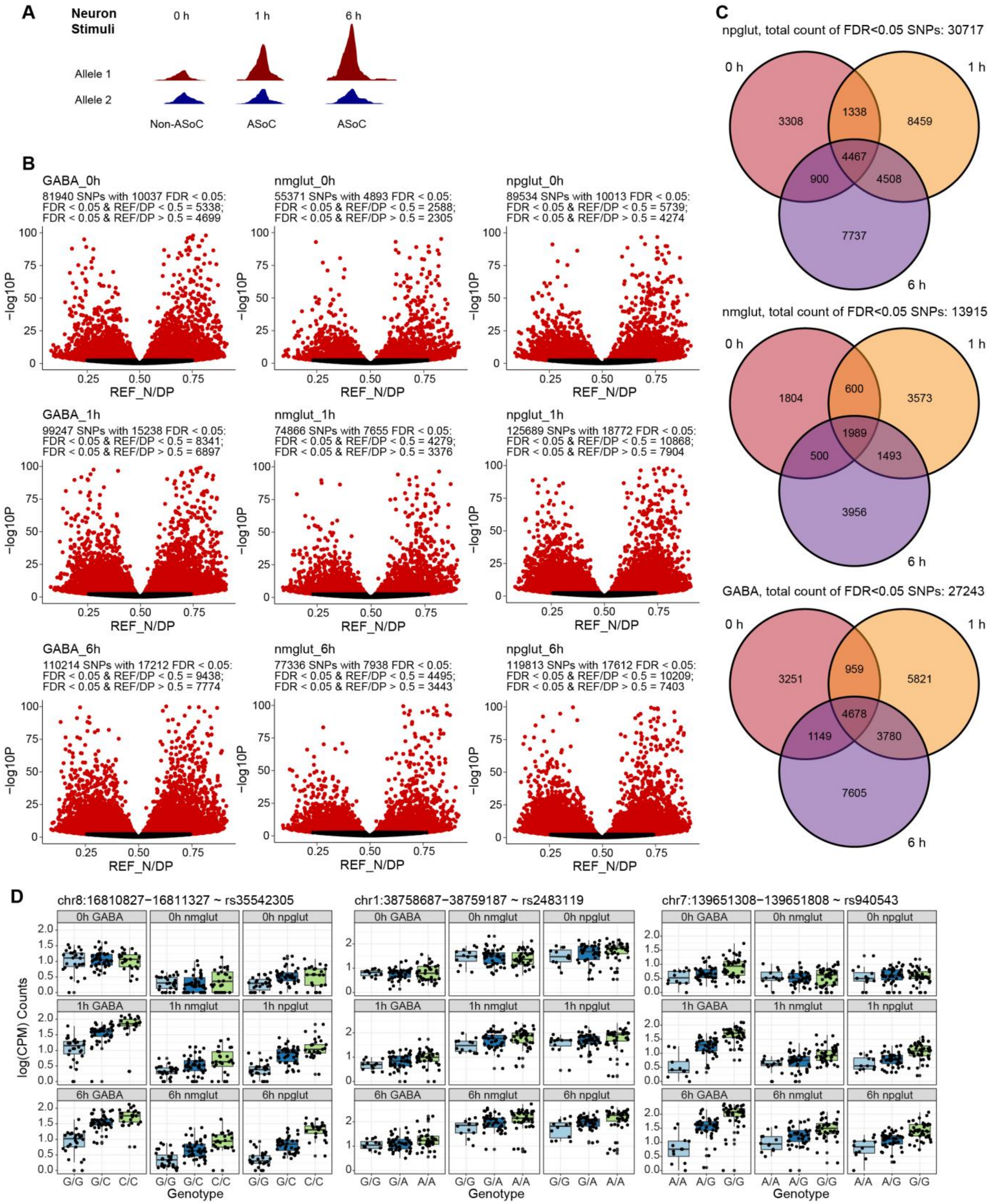
A
Neuron Stimuli
0 h
1 h
6 h
Allele 1
Allele 2
Non-ASoC
ASoC
ASoC
B
GABA_0h
81940 SNPs with 10037 FDR < 0.05:
FDR < 0.05 & REF/DP < 0.5 = 5338;
FDR < 0.05 & REF/DP > 0.5 = 4699
nmglut_0h
55371 SNPs with 4893 FDR < 0.05:
FDR < 0.05 & REF/DP < 0.5 = 2588;
FDR < 0.05 & REF/DP > 0.5 = 2305
npglut_0h
89534 SNPs with 10013 FDR < 0.05:
FDR < 0.05 & REF/DP < 0.5 = 5739;
FDR < 0.05 & REF/DP > 0.5 = 4274
GABA_1h
99247 SNPs with 15238 FDR < 0.05:
FDR < 0.05 & REF/DP < 0.5 = 8341;
FDR < 0.05 & REF/DP > 0.5 = 6897
nmglut_1h
74866 SNPs with 7655 FDR < 0.05:
FDR < 0.05 & REF/DP < 0.5 = 4279;
FDR < 0.05 & REF/DP > 0.5 = 3376
npglut_1h
125689 SNPs with 18772 FDR < 0.05:
FDR < 0.05 & REF/DP < 0.5 = 10868;
FDR < 0.05 & REF/DP > 0.5 = 7904
GABA_6h
110214 SNPs with 17212 FDR < 0.05:
FDR < 0.05 & REF/DP < 0.5 = 9438;
FDR < 0.05 & REF/DP > 0.5 = 7774
nmglut_6h
77336 SNPs with 7938 FDR < 0.05:
FDR < 0.05 & REF/DP < 0.5 = 4495;
FDR < 0.05 & REF/DP > 0.5 = 3443
npglut_6h
119813 SNPs with 17612 FDR < 0.05:
FDR < 0.05 & REF/DP < 0.5 = 10209;
FDR < 0.05 & REF/DP > 0.5 = 7403
-log10P
REF_N/DP
C
npglut, total count of FDR<0.05 SNPs: 30717
0 h
1 h
6 h
3308
1338
8459
4467
900
4508
7737
nmglut, total count of FDR<0.05 SNPs: 13915
1804
600
3573
1989
500
1493
3956
GABA, total count of FDR<0.05 SNPs: 27243
3251
959
5821
4678
1149
3780
7605
D
chr8:16810827−16811327 ~ rs35542305
chr1:38758687−38759187 ~ rs2483119
chr7:139651308−139651808 ~ rs940543
0h GABA
0h nmglut
0h npglut
1h GABA
1h nmglut
1h npglut
6h GABA
6h nmglut
6h npglut
log(CPM) Counts
G/G
G/C
C/C
G/G
G/A
A/A
A/A
A/G
G/G
Genotype

**Fig. S21. caQTL (including ASoC) mapping.**

(A) Schematics of neuron stimulation-specific (1 h and 6 h) ASoC SNP site. (B) Volcano plots show the ASoC SNPs (red dots) in each context (cell type × time point). Plotted are allele fractions of the reference alleles (Ref/total read depth) and -$\log_{10}P$ of the ASoC test. $FDR < 0.05$ for ASoC. (C) Venn diagrams show the overlaps of statistically significant ($FDR < 0.05$) ASoC SNPs of each cell type at three time points. (D) Box plots of three dynamic QTL, illustrating chromatin accessibility stratified by the genotype of the SNP, across time points for each cell type. Each dot represents log(CPM) (peak accessibility) of the SNP-associated cPeak of an individual cell line. CPM, counts per million reads.

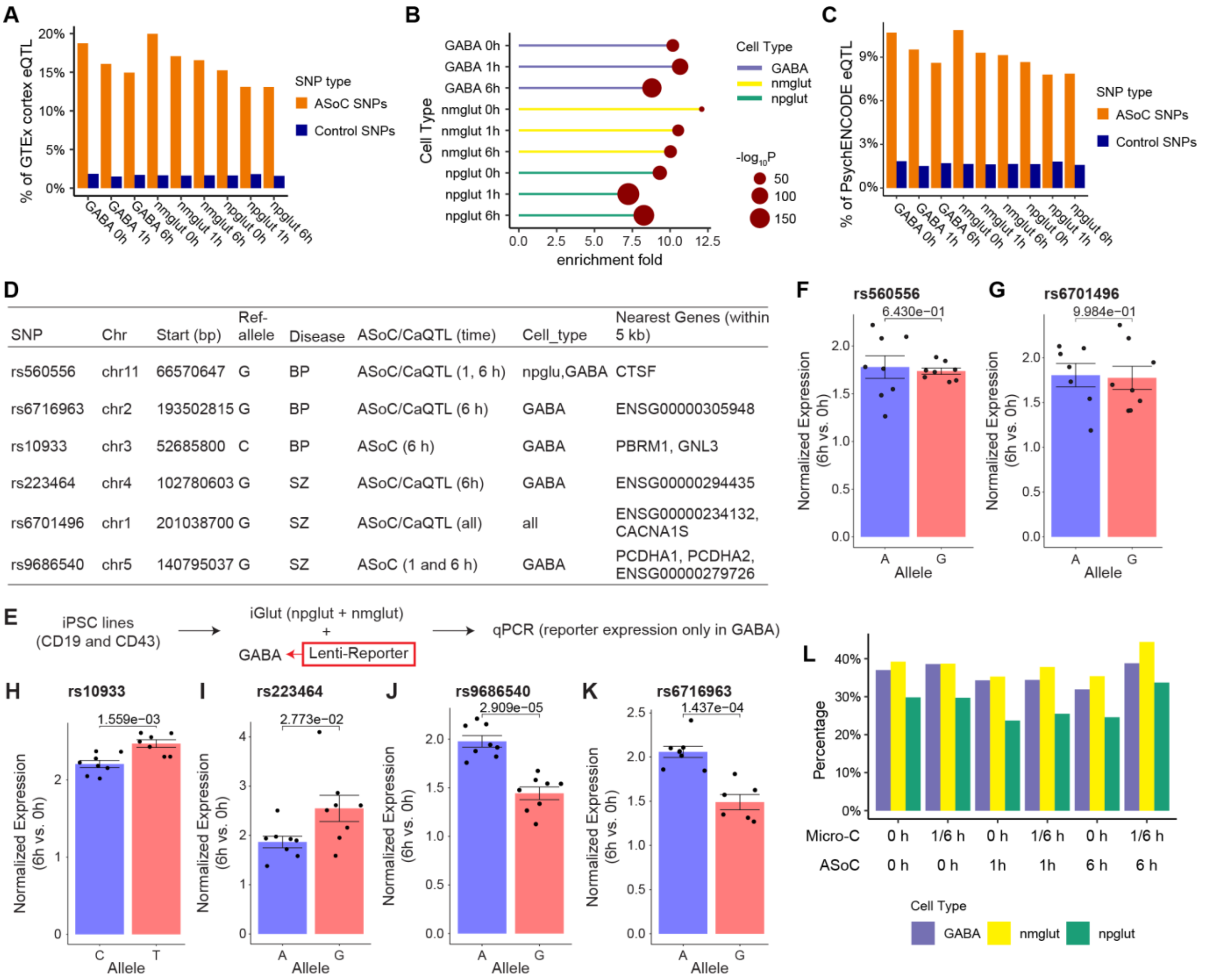

**Fig. S22. Functional validation of ASoC SNPs by comparing to brain eQTL, reporter gene assay, and Micro-C.**

(A) Bar plot shows the percentage of ASoC and control SNPs that are also GTEx cortex eQTL across contexts (cell types × time points). Control SNP sets contain random set of SNPs that are not ASoC. (B) The fold of enrichment and *P*-value for ASoC SNPs (vs. control non-ASoC SNPs) in (A). Fisher's exact test was used. (C) Bar plot shows the percentage of ASoC SNPs and control SNPs that are also PsychENCODE eQTL across contexts (cell types × time points). (D) to (K) Functional validation of ASoC SNPs by reporter gene assay. (D) The six selected ASoC SNPs that are also associated with SCZ and BP. Only ASoC SNPs in GABA were selected because of the technical challenge to distinguish npglut and nmglut in reporter gene assay. (E) The workflow for reporter gene assay. Two iPSC lines were used. GABA neurons were infected by Lenti-reporter before being co-cultured with iGlut. Oligos of 151bp (+/-75bp of the ASoC SNP site) were assayed for each allele. (F) to (K) QPCR quantification of the reporter gene expression for two alleles of an ASoC SNP for rs560556 (F), rs6701496 (G), rs10933 (H), rs223464 (I), rs9686540 (J), and rs6716963 (K). Data points, 3-4 biological replicates from each line. Shown are *P*-values from two-sided unpaired Student's *t*-test. (L) Proportion of ASoC SNPs that can be assigned to a *cis*-

target gene based on Micro-C chromatin contacts (gene TSS to OCRs) at matching context (cell type × time point).

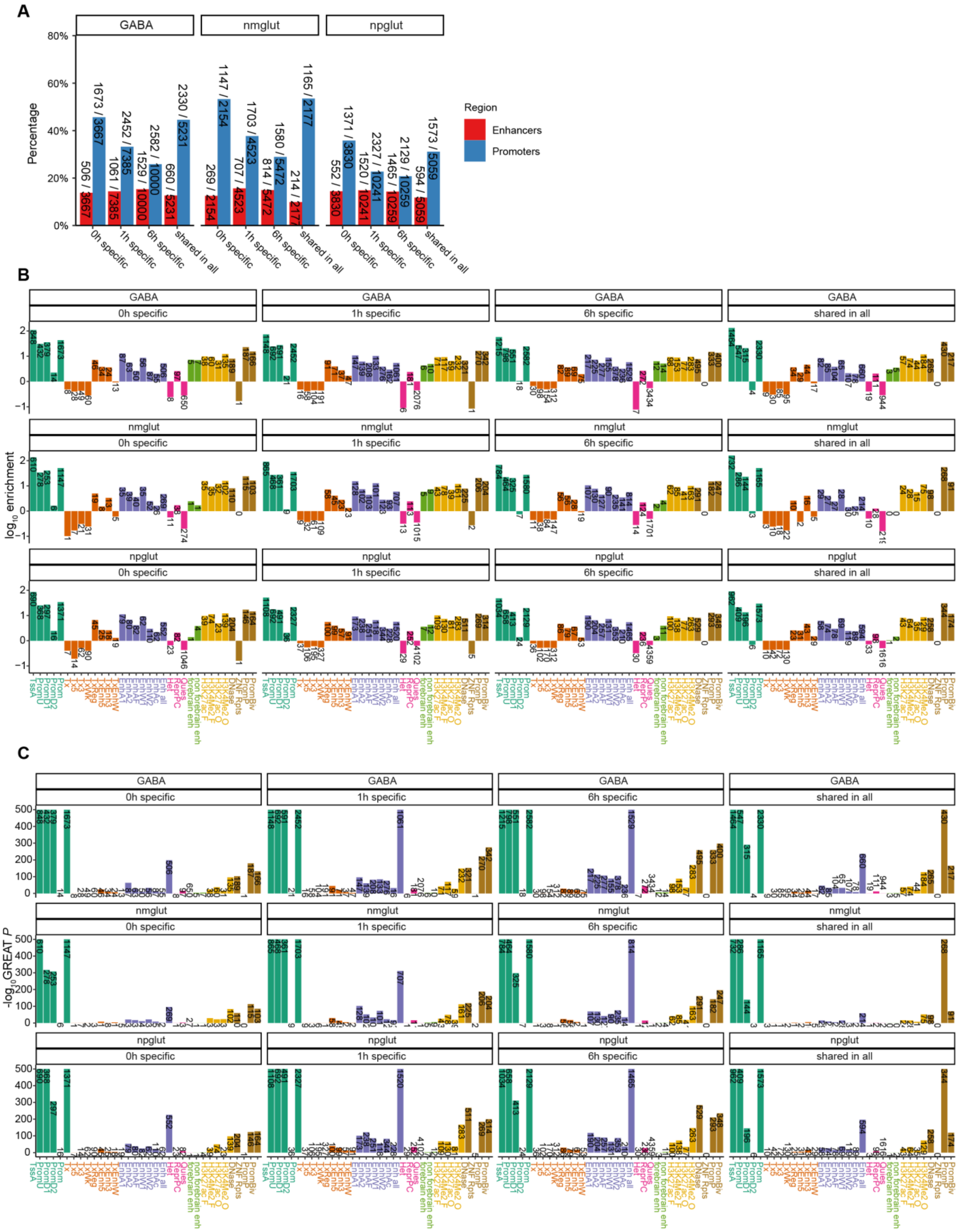

A
GABA
nmglut
npglut
Percentage
80%
60%
40%
20%
0%
Region
Enhancers
Promoters
0h specific
1h specific
6h specific
shared in all
B
log10 enrichment
C
-log10 GREAT P
TssA
PromU
PromD1
PromD2
Prom
Tx
Tx5
Tx3
TxWk
TxReg
TxEnh5
TxEnh3
TxEnhW
EnhA1
EnhA2
EnhAF
EnhW1
EnhW2
EnhAc
Enh all
Het
RepPC
Quies
forebrain enh
non forebrain enh
H3K27ac F
H3K27ac O
H3K4Me2 F
H3K4Me2 O
DNase
ZNF Rpts
PromP
PromBiv

**Fig. S23. Enrichment of ASoC SNPs in different regulatory sequence elements.**

(A) Bar plot shows the percentage and count of ASoC SNPs of each temporal category located in the annotated enhancer (red bars) or promoter (blue bars) regions. ASoC SNP categories in each cell type include those specific to each time point (0 h, 1 h, or 6 h) and shared by all time points. (B) and (C) Bar plots show the 25-way GREAT functional annotation of ASoC SNPs that are specific to or shared across time points in each cell type. (B) shows the fold of enrichment and SNP counts while (C) shows the GREAT enrichment *P*-value ($-\log_{10}P$) and SNP counts. The abbreviation of different chromatin states was as used in (*110*): TssA, Active TSS; PromU, Promoter Upstream TSS; PromD1, Promoter Downstream TSS with DNase; PromD2, Promoter Downstream TSS; Tx5', Transcription 5'; Tx, Transcription; Tx3', Transcription 3'; TxWk, Weak transcription; TxReg, Transcription Regulatory; TxEnh5', Transcription 5' Enhancer; TxEnh3', Transcription 3' Enhancer; TxEnhW, Transcription Weak Enhancer; EnhA1, Active Enhancer 1; EnhA2, Active Enhancer 2; EnhAF, Active Enhancer Flank; EnhW1, Weak Enhancer 1; EnhW2, Weak Enhancer 2; EnhAc, Enhancer Acetylation Only; Dnase, DNase only; ZNF/Rpts, ZNF genes & repeats; Het, Heterochromatin; PromP, Poised Promoter; PromBiv, Bivalent Promoter; ReprPC, Repressed PolyComb; Quies, Quiescent; Forebrain_enh, Forebrain enhancers; Non_forebrain_enh, Non-forebrain enhancers; H3K27acF, H3K27 acetylated regions found in the frontal lobe; H3K27acO, H3K27 acetylated regions found in the occipital lobe; H3K4me2F, H3K4me2 regions found in the frontal lobe; H3K4me2O, H3K4me2 regions found in the occipital lobe.

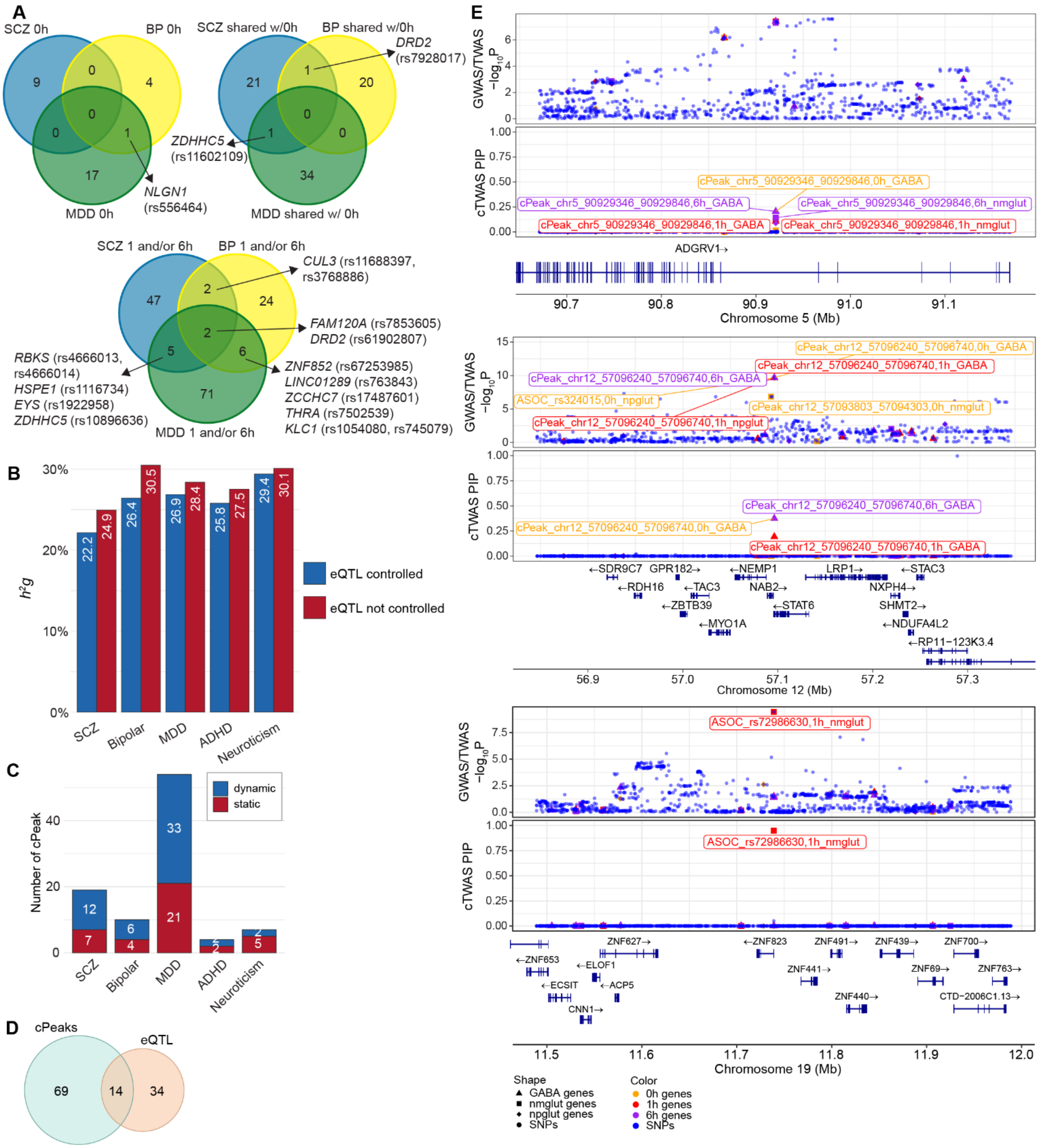

**Fig. S24. Integrative analysis of caQTL and GWAS of NPD phenotypes.**

(A) Venn diagrams showing the overlapping SCZ, BP, and MDD GWAS associated SNPs (and their nearest genes) that also showed ASoC. Top left, SNPs showing ASoC at 0 h only; top right, SNPs showing ASoC at 0 h and upon stimulation (i.e., shared); bottom, SNPs showing ASoC only at 1 h and/or 6 h (i.e., stimulation-specific). (B) The proportion of $h^2_g$ mediated by caQTL and

ASoCs with or without controlling for eQTL in cTWAS. (C) The number of risk peaks found by cTWAS, across the five NPD phenotypes stratified by peaks being dynamic or static. “Dynamic” or “Static” according to how PIPs are partitioned across contexts (see Fig. 4D caption). (D) The Venn diagram between risk peaks (from caQTL-based cTWAS) and risk genes (from eQTL-based cTWAS) with PIP over 0.5 for SCZ. They are overlapped if the corresponding caQTL/ASoC SNP is located within 500 kb distance from the eQTL SNP. (E) Locus plots of the SCZ risk cPeaks that can be assigned to a unique gene. For each locus plot, the top panel is GWAS *P* values of the SNPs and TWAS *P* values of the cPeaks (or ASoC SNP), the bottom panel is cTWAS PIP of cPeaks (or ASoC SNP). Note that the cPeak for each highlighted gene has combined PIP > 0.8 from all cellular contexts.

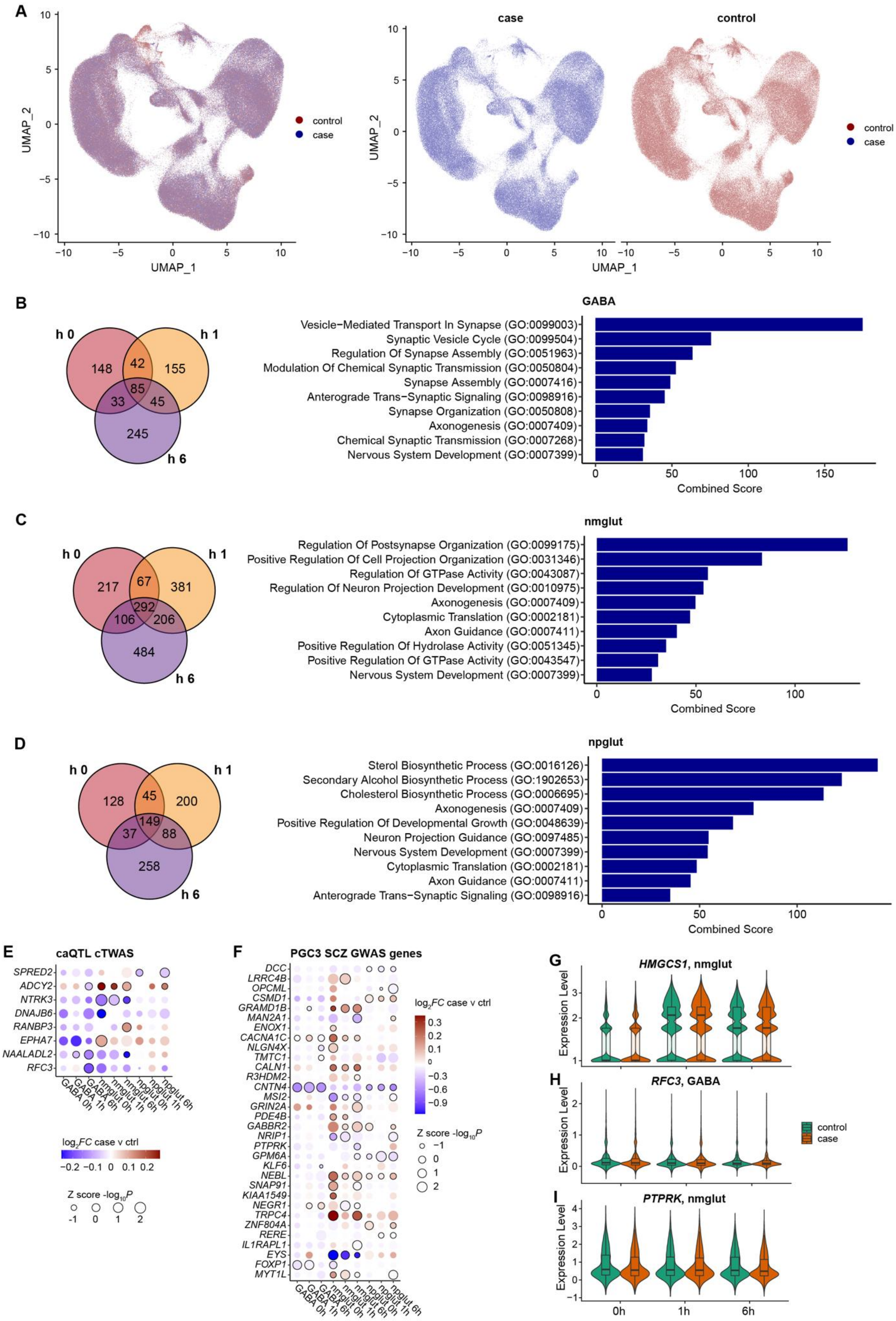
A
UMAP_1
UMAP_2
case
control
control
case
B
h 0
h 1
h 6
148
42
155
85
33
45
245
GABA
Vesicle−Mediated Transport In Synapse (GO:0099003)
Synaptic Vesicle Cycle (GO:0099504)
Regulation Of Synapse Assembly (GO:0051963)
Modulation Of Chemical Synaptic Transmission (GO:0050804)
Synapse Assembly (GO:0007416)
Anterograde Trans−Synaptic Signaling (GO:0098916)
Synapse Organization (GO:0050808)
Axonogenesis (GO:0007409)
Chemical Synaptic Transmission (GO:0007268)
Nervous System Development (GO:0007399)
Combined Score
C
h 0
h 1
h 6
217
67
381
292
106
206
484
nmglut
Regulation Of Postsynapse Organization (GO:0099175)
Positive Regulation Of Cell Projection Organization (GO:0031346)
Regulation Of GTPase Activity (GO:0043087)
Regulation Of Neuron Projection Development (GO:0010975)
Axonogenesis (GO:0007409)
Cytoplasmic Translation (GO:0002181)
Axon Guidance (GO:0007411)
Positive Regulation Of Hydrolase Activity (GO:0051345)
Positive Regulation Of GTPase Activity (GO:0043547)
Nervous System Development (GO:0007399)
Combined Score
D
h 0
h 1
h 6
128
45
200
149
37
88
258
npglut
Sterol Biosynthetic Process (GO:0016126)
Secondary Alcohol Biosynthetic Process (GO:1902653)
Cholesterol Biosynthetic Process (GO:0006695)
Axonogenesis (GO:0007409)
Positive Regulation Of Developmental Growth (GO:0048639)
Neuron Projection Guidance (GO:0097485)
Nervous System Development (GO:0007399)
Cytoplasmic Translation (GO:0002181)
Axon Guidance (GO:0007411)
Anterograde Trans−Synaptic Signaling (GO:0098916)
Combined Score
E
caQTL cTWAS
SPRED2
ADCY2
NTRK3
DNAJB6
RANBP3
EPHA7
NAALADL2
RFC3
GABA 0h
GABA 1h
GABA 6h
nmglut 0h
nmglut 1h
nmglut 6h
npglut 0h
npglut 1h
npglut 6h
log2FC case v ctrl
Z score -log10P
F
PGC3 SCZ GWAS genes
DCC
LRRC4B
OPCML
CSMD1
GRAMD1B
MAN2A1
ENOX1
CACNA1C
NLGN4X
TMTC1
CALN1
R3HDM2
CNTN4
MSI2
GRIN2A
PDE4B
GABBR2
NRIP1
PTPRK
GPM6A
KLF6
NEBL
SNAP91
KIAA1549
NEGR1
TRPC4
ZNF804A
RERE
IL1RAPL1
EYS
FOXP1
MYT1L
log2FC case v ctrl
Z score -log10P
G
HMGCS1, nmglut
Expression Level
H
RFC3, GABA
Expression Level
control
case
I
PTPRK, nmglut
Expression Level
0h
1h
6h

**Fig. S25. Differential expressed genes (DEGs) in neurons from SCZ cases.**

(A) UMAP projection of snRNA-seq data of neurons from the 28 SCZ cases and 28 controls matched by sex and age. (B)-(D) Venn diagram showing SCZ-associated DEGs at the three time points of neuronal activation (0 h, 1 h, and 6 h) (left) as well as the top 10 enriched GO terms (biological processes) (right) among stimulation-specific DEGs in three different neuron subtypes: GABA (B), nmglut (C), and npglut (D). All the listed GO terms have *FDR* < 0.05 and are ranked by their combined enrichment score from Enrichr. (E) and (F) Bubble plots show the $\log_2$FC and Z-scored -$\log_{10}P$ of DE from MAST test for cTWAS (caQTL-based) SCZ genes and PGC3 SCZ GWAS prioritized single genes, respectively. Only genes that are also SCZ-associated DEGs (*FDR*<0.05) at any context (cell type and time points) are included. (G) to (I) Violin plots of SCZ case-control single-cell expression of example genes in cholesterol metabolic gene set (HMGCS1) (G), SCZ cTWAS gene set (RFC3) (H), and SCZ GWAS prioritized single gene set (PTPTK) (I). Only the cell type showing the strongest DE in SCZ cases is shown for each gene.

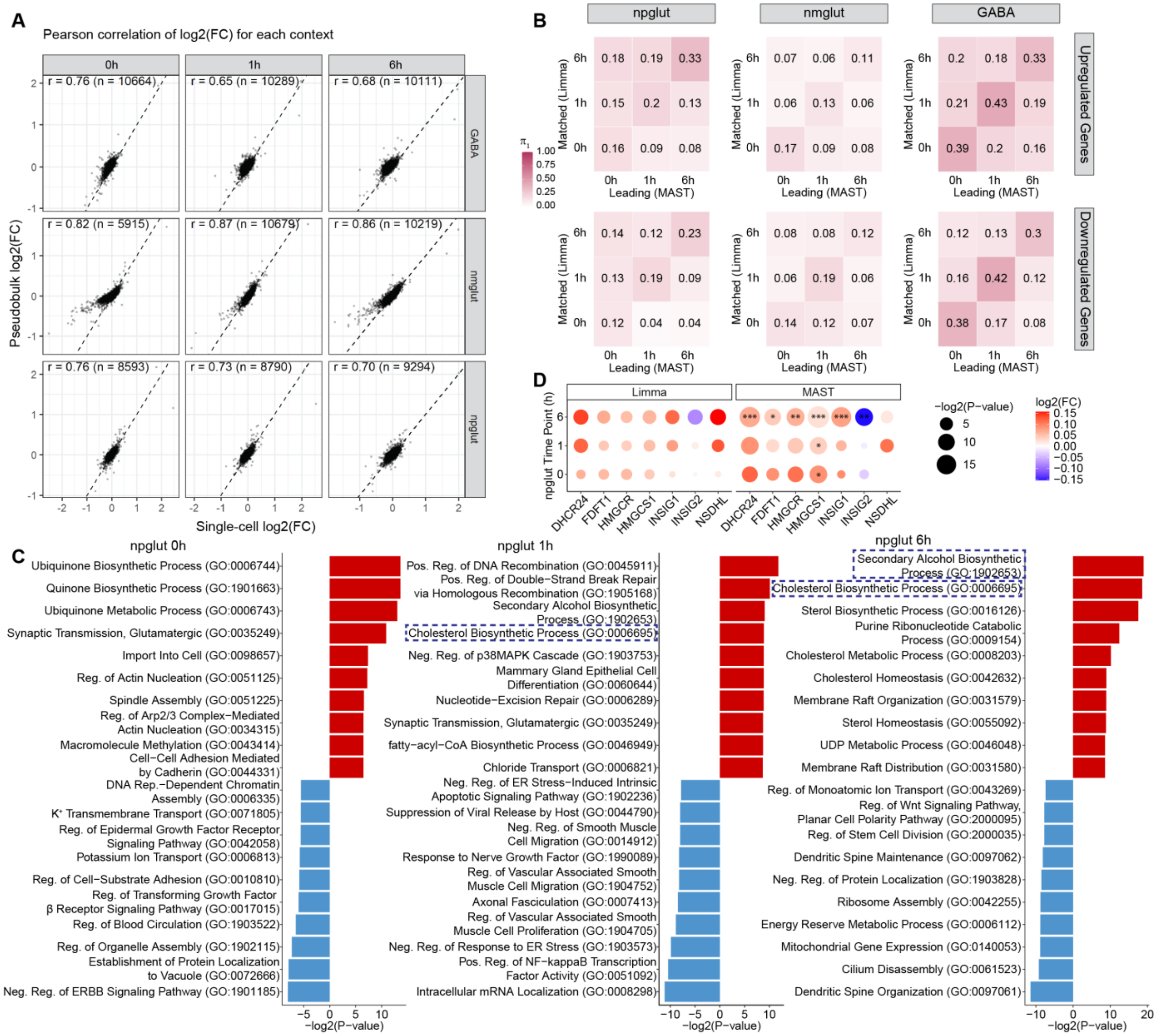

**Fig. S26. Consistent DEG results from using pseudobulk or single-cell data.**

(A) Strong correlation of $\log_2$FC between Pseudobulk-Limma and single-cell-MAST analyses. (B) Proportion of DEGs (*FDR* < 0.05) from MAST analysis that was shared with pseudobulk-Limma DEGs (*P* <0.05; only 2 DEGs with *FDR* < 0.05 across all contexts) in Pi1 analysis (Pi1 = 0.11 to 0.43; 13 to 18-fold enrichments; hypergeometric *P* values < $2.5 \times 10^{-29}$ to $7.9 \times 10^{-88}$). Note the larger Pi1 for the matching cellular contexts between sc-MAST results and pseudobulk-Limma results. (C) GO enrichment analysis of the pseudobulk DEGs (nominal *P* < 0.05) for each context. All the listed GO terms are ranked by their enrichment *P* values from Enrichr. Red bars, upregulated; blue bars, downregulated. Note the top-ranking cholesterol synthesis related GO among upregulated genes (highlighted in dashed boxes) upon stimulation. (D) Consistent DEG results for specific cholesterol synthesis genes from limma (Pseudobulk) and MAST (single-cell) analyses. *, *FDR* < 0.05; **, *FDR* < 0.01; ***, *FDR* < 0.001.

**List of Supplemental Tables**

(The supplementary tables have been included in the manuscript. These tables are also available at **https://doi.org/10.5061/dryad.0zpc8677w**.)

Table S1. Donor information of all the iPSC lines and their summary statistics of sn-multiomics data for all three time points of KCl stimulation.

Table S2. Summary statistics of each sequencing library before and after QC. Each sequencing library is for cells of a co-cultured 1-5 iPSC lines at a particular time point of stimulation. TSS, transcription starting site.

Table S3. SnRNA-seq post-QC summary data for all 100 lines.

Table S4. Differentially expressed genes upon KCl stimulation in each cell type.

Table S5. NPD gene sets used for enrichment analyses.

Table S6. Differentially accessible peaks upon KCl stimulation in each cell type.

Table S7. Genes included in each gene module (cluster).

Table S8. GO terms (biological process) enriched for each gene module.

Table S9. OCR-gene pairs identified through single-cell level peak-gene correlation (co-activation) analysis.

Table S10. Micro-C summary statistics.

Table S11. Micro-C chromatin contacts in each sample. Listed are 5 kb bins.

Table S12. Enhancer-gene pairs predicted by the ABC (Activity-by-Contact) model, with an ABC score ≥ 0.021. Predictions from all nine contexts are included.

Table S13. Differential TF motif activity analysis using a one-tailed Wilcoxon signed-rank test.

Table S14. Gene regulatory networks identified in three cell types.

Table S15. Target genes regulated by the ASD risk TFs in each cell type.

Table S16. GO-term (Biological Process) enrichment for four ASD risk-associated TFs. The *FDR* was recalculated to account for multiple testing across the four TFs.

Table S17. GO-term (Molecular Function) enrichment of target genes shared by at least three of the ASD risk TFs. Only "molecular function" terms are enriched.

Table S18. TFs with target genes enriched for ASD genes in each cell type.

Table S19. Significant eQTL in each cell type and time point.

Table S20. Dynamic cGenes in each cell type and time point.

Table S21. eQTL-based cTWAS results.

Table S22. GO term enrichment of cTWAS NPD genes.

Table S23. cPeak and caQTL identified by tensorQTL in each cell type and time point.

Table S24. ASoC SNPs in each cell type and time point.

Table S25. Dynamic caQTL for each cell type with permutation ($n$=1,000) testing result.

Table S26. Percentage of ASoC SNPs that can be assigned to a Micro-C based *cis*-target gene in a matching context.

Table S27. ASoC status of Schizophrenia (SCZ) GWAS risk SNPs and their LD proxies ($R^2$>0.8) in each cell type and time point. Brain eQTL and Micro-C target annotations are from column BN to BT.

Table S28. ASoC status of Bipolar disorder (BP) GWAS risk SNPs and their LD proxies ($R^2$>0.8) in each cell type and time point. Brain eQTL and Micro-C target annotations are from column BN to BT.

Table S29. ASoC status of Major depression disorder (MDD) GWAS risk SNPs and their LD proxies ($R^2$>0.8) in each cell type and time point. Brain eQTL and Micro-C target annotations are from column BN to BT.

Table S30. caQTL-based cTWAS results.

Table S31. Result of single cell DEG analysis in neurons of each context (Cell type × time point after stimulation) between 28 SCZ cases and matched controls. BH_*FDR* values were derived from the MAST *p*-value for each context.

Table S32. sgRNA sequences and PCR primer sequences as well as qPCR assays.

Table S33. GWAS datasets used in enrichment tests (MAGMA, TORUS, sLDSC, and cTWAS).

**References and Notes (only appeared in Supplementary Materials)**
